# Sex-specific regulation of binge drinking, social, and arousal behaviors by subcortical serotonin 5HT_2c_ receptor-containing neurons

**DOI:** 10.1101/2022.01.28.478036

**Authors:** M.E. Flanigan, O.J. Hon, S. D’Ambrosio, K.M. Boyt, L. Hassanein, M. Castle, H.L. Haun, M.M. Pina, T.L. Kash

## Abstract

Serotonin 5HT_2c_ receptors have been implicated in the pathophysiology of both mood disorders and alcohol use disorder, but the circuits mediating the effects of systemic pharmacological manipulations of this receptor on behavior have not been identified. Binge alcohol consumption induces discrete social and arousal disturbances in human populations, which are thought to promote increased drinking. However, whether models of binge drinking in rodents can induce these same long-term negative behavioral symptoms is unknown. In this study, we employed multiple anatomical, physiological, and behavioral approaches to identify two populations of neurons expressing serotonin 5HT_2c_ receptors, one in the lateral habenula (LHb_5HT2c_) and one in the bed nucleus of the stria terminalis (BNST_5HT2c_), that display coordinated *in-vivo* responses to social, arousal, and alcohol-related stimuli and are physiologically modulated by binge alcohol consumption in a sex-specific manner. Critically, these physiological changes were associated with sex-specific behavioral disturbances that closely mirror social and arousal symptoms observed in humans during abstinence from binge drinking. Specifically, we observed that one week of abstinence from three weeks of binge alcohol drinking induced social recognition deficits in females and potentiated acoustic startle responses in males. While both populations of neurons (BNST and LHb) and the 5HT_2c_ receptor itself contribute to the sex-specific effects of alcohol on social and arousal behaviors to some degree, the primary causal mechanism underlying these phenomena appears to be excessive activation of LHb_5HT2c_ neurons. These findings may have implications for the development of sex-specific treatments for mood and alcohol use disorders targeting the brain’s serotonin system.

## Introduction

Binge alcohol drinking is a significant cause of alcohol-related death, illness, and economic burden (SAMSHA, 2019). Moreover, repeated cycles of binge drinking and withdrawal increase the incidence of negative social and emotional states, which can promote further increases in drinking and the transition to alcohol dependence (Koob, 2021). Thus, understanding the neurobiological mechanisms mediating the negative effects of alcohol consumption on behavior may be critical for limiting subsequent escalations in alcohol intake. In humans, abstinence from binge alcohol consumption is strongly associated with both impaired social emotion recognition behavior and enhanced arousal (Castellano et al., 2015; Frigerio et al., 2002; Lannoy et al., 2018; Rupp et al., 2017), and repeated cycles of binge drinking and abstinence exacerbate these issues (Gorka, 2020; Kang et al., 2018; Krystal et al., 1997; Loeber et al., 2007; Marin et al., 2015; Moberg et al., 2017; Schellekens et al., 2012). However, whether binge alcohol consumption dysregulates social recognition and arousal behavior in adult rodents has not been explored.

The brain’s serotonin (5-hydroxytryptamine, 5-HT) system is a critical modulator of affect, motivation, social behavior, arousal, and metabolism (da Cunha-Bang and Knudsen, 2021; Georgescu et al., 2021; Hayes and Greenshaw, 2011; Vahid-Ansari and Albert, 2021). Dysregulation of 5-HT signaling has been implicated in the pathophysiology of numerous psychiatric disorders, including major depression, anxiety disorders, substance use disorders, obsessive-compulsive disorder, and schizophrenia (Nordquist and Oreland, 2010). 5-HT neurons are located primarily in hindbrain raphe nuclei, specifically the dorsal (DRN) and median (MRN) raphe nuclei. These neurons project widely throughout the brain, including to cortical, amygdalar, midbrain, and other hindbrain regions (for review, see (Commons, 2020)). A recent anatomical and functional mapping study of the DRN revealed that there are distinct functional populations of 5-HT neurons localized to discrete sub-regions of the DRN (Ren et al., 2018). While 5-HT is implicated in Alcohol Use Disorder (AUD) pathophysiology (Marcinkiewcz et al., 2016a), studies in humans and animal models report that manipulations of 5-HT signaling can both increase and decrease alcohol consumption and associated negative affect (Campbell et al., 2021; Filip et al., 2012; Fu et al., 2020; Marcinkiewcz et al., 2015; Marcinkiewcz et al., 2016a; Tabbara et al., 2021; Umhau et al., 2011). The heterogeneity of these observed effects is likely due to differential recruitment of discrete 5-HT circuits and receptor sub-types in AUD regulating specific motivational, cognitive, and social processes.

Both the lateral habenula (LHb) and the bed nucleus of the stria terminalis (BNST) receive inputs from serotonergic neurons located in the dorsal and caudal regions of the dorsal-raphe nucleus (Abrams et al., 2004; Commons, 2020; Muzerelle et al., 2016) and, according to the aforementioned study (Ren et al., 2018), may represent two key target regions of an “aversive” subcortical stream of serotonin signaling. Activation of LHb or BNST neurons generally promotes negative emotional states and can reduce motivation for natural and drug rewards (Avery et al., 2016; Baker et al., 2016; Hikosaka, 2010; Lawson et al., 2016; Lebow and Chen, 2016; Velasquez et al., 2014), and large subsets of neurons in both regions are depolarized by 5-HT through activation of the 5HT_2c_ receptor (Guo et al., 2009; Hammack et al., 2009; Marcinkiewcz et al., 2016b). 5HT_2c_ is a primarily Gq-protein-coupled receptor and is highly edited at the RNA level, resulting in a large number of unique isoforms that engage discrete intracellular signaling pathways (Berg et al., 2008). Systemic antagonism of 5HT_2c_ has been shown to reduce behaviors related to anxiety and depression in both alcohol-exposed and alcohol-naïve male rodents (Marcinkiewcz et al., 2015; Opal et al., 2014; Papp et al., 2020), but both 5HT_2c_ agonists and antagonists can reduce alcohol intake (Rezvani et al., 2014; Tabbara et al., 2021; Yoshimoto et al., 2012). While few studies have investigated the effects of functional modulation of LHb or BNST 5HT_2c_ receptors on behavior, antagonism of 5HT_2c_ in the LHb of male rats reduces both alcohol self-administration and withdrawal-induced anxiety-like behavior in the open field test (Fu et al., 2020). These studies support the potential role of 5HT_2c_ signaling in these structures in the regulation of alcohol drinking and affective behavior while also highlighting the need for future studies, particularly in female subjects.

In this study, we first characterized the anatomy, physiology, and behavioral function of the DRN-LHb and DRN-BNST projections in binge alcohol drinking, arousal behavior, and social behavior, with a specific focus on 5-HT inputs to 5HT_2c_-receptor containing neurons in the LHb and the BNST. We establish a model whereby three weeks of binge alcohol intake induces long-lasting behavioral and physiological adaptations that are similar to those observed in binge-drinking humans. Using complementary approaches for brain region-specific genetic deletion of 5HT_2c_ expression and chemogenetic manipulation of G-protein signaling in 5HT_2c_ neurons, we identify sex- and region-specific roles for 5HT_2c_ receptors themselves as well as 5HT_2c_-containing neurons in social and arousal behaviors in the context of binge alcohol consumption. Overall, our results suggest that binge alcohol consumption potentiates DRN-BNST and DRN-LHb circuitry to promote disruptions in affective behavior, but that excessive activation of the LHb_5HT2c_ neurons is the primary causal mechanism promoting behavioral dysfunction by alcohol.

## Results

### LHb_5HT2c_ and BNST_5HT2c_ neurons mount coordinated responses to rewarding and aversive stimuli

Given the potential role of 5HT_2c_ signaling in the LHb and BNST in regulating binge drinking and affective behavior, we first wanted to understand how neurons that contain these receptors respond to aversive and rewarding stimuli *in-vivo*. First, male and female 5HT_2c_-cre mice (Burke et al., 2016) were unilaterally injected with a cre-dependent AAV encoding the calcium sensor GCaMP7f in the LHb and BNST and optical fibers were implanted above these regions. Using fiber photometry we recorded calcium signals from both LHb_5HT2c_ and BNST_5H2c_ neurons during free social interaction with a novel juvenile same-sex conspecific, the acoustic startle test, voluntary water drinking, and voluntary alcohol drinking (Figure 1A, B). We focused our efforts on these behaviors and have been previously linked to 5HT_2c_ function in these regions and shown to be disrupted in human binge drinkers. Importantly, we were able to correct for any motion artifacts during recording by subtracting out signal from our isosbestic control wavelength channel (405nm). In general, both populations of 5HT_2c_ neurons in both sexes were activated by social interaction and acoustic startle stimuli, but inhibited by water and alcohol consumption (Figure 1C-T). However, males displayed more robust BNST_5HT2c_ responses to these stimuli. For example, relative to females, males displayed increased activation in response to social targets (Figure 1M, N), but stronger inhibition in response to alcohol (Figure 1Q, R) and water consumption (Figure 1S, T). In LHb_5HT2c_ neurons, however, females mounted a strong response to water consumption (Figure 1I, J) but not to alcohol consumption (Figure 1K, L), while males mounted similarly strong responses to both water (Figure 1S, T) and alcohol (Figure 1Q, R). Notably, mice were water deprived prior to voluntary drinking experiments in order to facilitate consumption while tethered to the fiber photometry apparatus. Under these conditions, we would expect that both water and alcohol consumption would be rewarding, yet only water consumption elicited a response in females, which may be suggestive of mixed ‘reward/aversion’ effects of alcohol in **binge**-naïve LHb. Remarkably, LHb_5HT2c_ and BNST_5HT2c_ neurons displayed robust overlap of their activity during these behaviors (Figure 1C, D), suggesting that their activity may be regulated by common upstream inputs.

**Figure 1:**
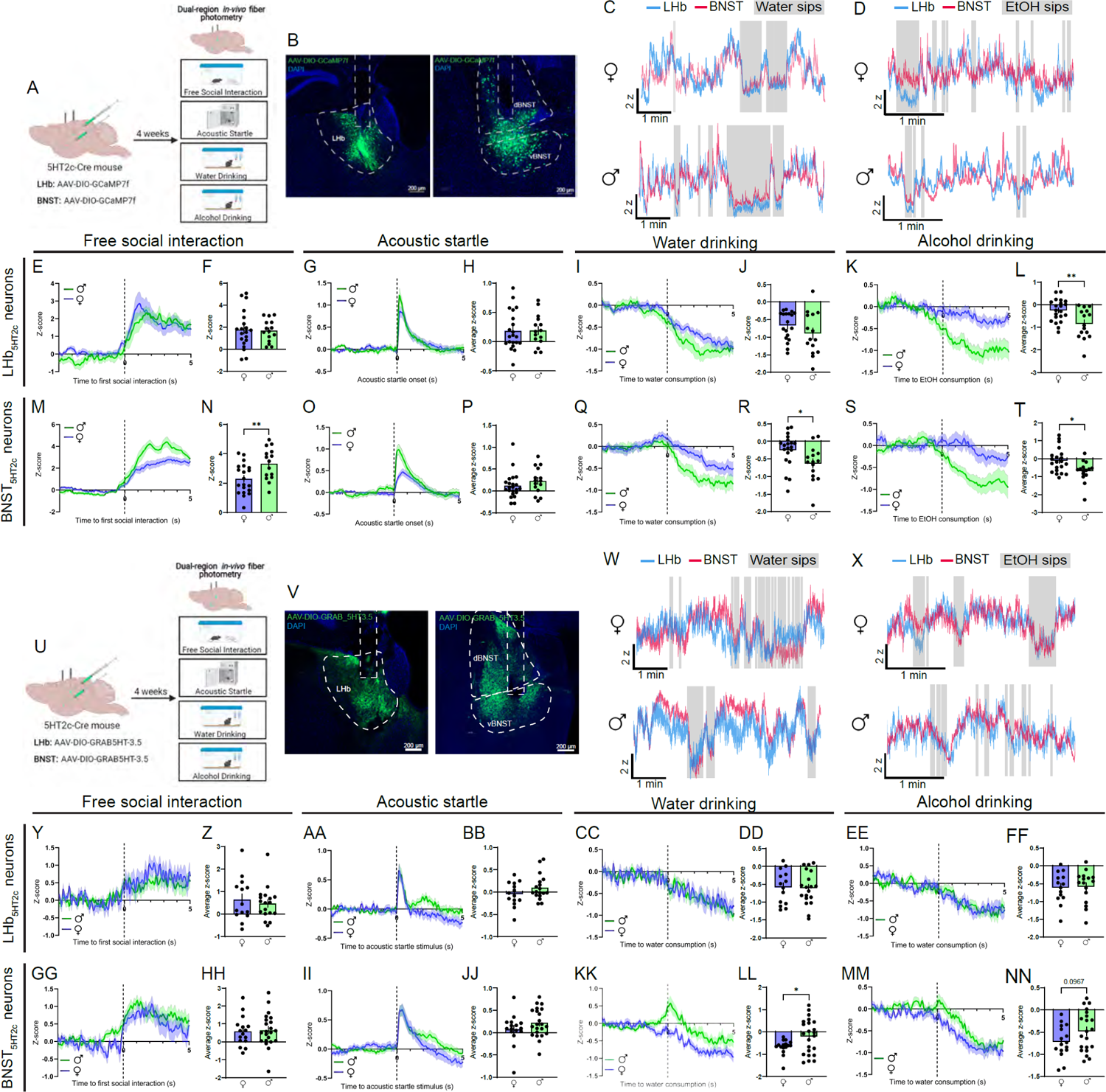
*In-vivo* LHb_5HT2c_ and BNST_5HT2c_ calcium and 5HT signals are modulated by exposure to affective stimuli. **A,** Surgical schematic and experimental timeline for GCaMP fiber photometry experiments. **B,** Representative viral infections and fiber placements in LHb (left) and BNST (right). **C,** Representative traces for water drinking session. **D,** Representative traces for alcohol drinking session. **E,** Peri-event plot of LHB_5HT2c_ GCaMP activity during the first interaction with a novel juvenile social target (one trial/mouse). **F,** Average z-score of LHb_5HT2c_ GCaMP signal for 0-5s post interaction (unpaired t-test, t(33)=0.1939, p=0.8474). **G,** Peri-event plot of LHb_5HT2c_ GCaMP activity during the acoustic startle test (average of 10 trials/mouse). **H,** Average z-score of LHb_5HT2c_ GCaMP signal for 0-5s post startle stimulus (unpaired t-test, t(34)=0.06731, p=0.94767). **I,** Peri-event plot of LHb_5HT2c_ GCaMP activity during voluntary water consumption (average of 1-3 bouts/mouse). **J**, Average z-score of LHb_5HT2c_ GCaMP signal for 0-5s post water drinking bout start (unpaired t-test, t(32)=1.333, p=0.192). **K,** Peri-event plot of LHb_5HT2c_ GCaMP activity during voluntary alcohol consumption (average of 1-3 bouts/mouse). **L,** Average z-score of LHb_5HT2c_ GCaMP signal for 0-5s post alcohol drinking bout start (unpaired t-test, t(34)=3.102, p=0.0038). **M,** Peri-event plot of BNST_5HT2c_ GCaMP activity during the first interaction with a novel juvenile social target (one trial/mouse). **N,** Average z-score of BNST_5HT2c_ GCaMP signal for 0-5s post interaction (n=15 males, n=20 females, unpaired t-test, t(33)=2.858, p=0.0073). **O,** Peri-event plot of BNST_5HT2c_ GCaMP activity during the acoustic startle test (average of 10 trials/mouse). **P,** Average z-score of BNST_5HT2c_ GCaMP signal for 0-5s post startle stimulus (unpaired t-test, t(35)=1.121, p=0.2698. **Q,** Peri-event plot of BNST_5HT2c_ GCaMP activity during voluntary water consumption (average of 1-3 bouts/mouse). **R,** Average z-score of BNST_5HT2c_ GCaMP signal for 0-5s post water drinking bout start (unpaired t-test, t(33)=2.274, p=0.0296. **S,** Peri-event plot of BNST_5HT2c_ GCaMP activity during voluntary alcohol consumption (average of 1-3 bouts/mouse). **T,** Average z-score of BNST_5HT2c_ GCaMP signal for 0-5s post water drinking bout start (unpaired t-test, t(34)=2.717, p=0.0103. **U,** Surgical schematic and experimental timeline for GRAB-5HT fiber photometry experiments. **V,** Representative viral infections and fiber placements in LHb (left) and BNST (right). **W,** Representative traces for water drinking session. **X,** Representative traces for alcohol drinking session. **Y,** Peri-event plot of LHB_5HT2c_ GRAB-5HT activity during the first interaction with a novel juvenile social target (one trial/mouse). **Z,** Average z-score of LHb_5HT2c_ GRAB-5HT signal for 0-5s post interaction (unpaired t-test, t(30)=0.5733, p=0.5707). **AA,** Peri-event plot of LHb_5HT2c_ GRAB-5HT activity during the acoustic startle test (average of 10 trials/mouse). **BB,** Average z-score of LHb_5HT2c_ GRAB-5HT signal for 0-5s post startle stimulus (unpaired t-test, t(30)=1.483, p=0.1486). **CC,** Peri-event plot of LHb_5HT2c_ GRAB-5HT activity during voluntary water consumption (average of 1-3 bouts/mouse). **DD**, Average z-score of LHb_5HT2c_ GRAB-5HT signal for 0-5s post water drinking bout start (unpaired t-test, t(31)=0.1558, p=0.8772). **EE,** Peri-event plot of LHb_5HT2c_ GRAB-5HT activity during voluntary alcohol consumption (average of 1-3 bouts/mouse). **FF,** Average z-score of LHb_5HT2c_ GRAB-5HT signal for 0-5s post alcohol drinking bout start (unpaired t-test, t(28)=0.1442, p=0.8864). **GG,** Peri-event plot of BNST_5HT2c_ GRAB-5HT activity during the first interaction with a novel juvenile social target (one trial/mouse). **HH,** Average z-score of BNST_5HT2c_ GRAB-5HT signal for 0-5s post interaction (unpaired t-test, t(36)=0.238, p=0.8133). **II,** Peri-event plot of BNST_5HT2c_ GRAB-5HT activity during the acoustic startle test (average of 10 trials/mouse). **JJ,** Average z-score of BNST_5HT2c_ GRAB-5HT signal for 0-5s post startle stimulus (unpaired t-test, t(37)=1.079, p=0.2874. **KK,** Peri-event plot of BNST_5HT2c_ GRAB-5HT activity during voluntary water consumption (average of 1-3 bouts/mouse). **LL,** Average z-score of BNST_5HT2c_ GRAB-5HT signal for 0-5s post water drinking bout start (unpaired t-test, t(37)=2.214, p=0.0331. **MM,** Peri-event plot of BNST_5HT2c_ GRAB-5HT activity during voluntary alcohol consumption (average of 1-3 bouts/mouse). **NN,** Average z-score of BNST_5HT2c_ GRAB-5HT signal for 0-5s post water drinking bout start (unpaired t-test, t(35)=1.707, p=0.0967. n=14-23 mice/group. All data are represented as mean ± SEM. *p<0.05, **p<0.01.

To characterize 5-HT dynamics, a separate cohort of 5HT_2c_-cre male and female mice were unilaterally injected with a cre-dependent AAV encoding the 5-HT sensor GRAB-5HT in the LHb and BNST and optical fibers were implanted above these regions (Figure 1U, V). This sensor fluoresces upon binding of 5-HT, which allows us to visualize 5-HT dynamics with millisecond temporal resolution (Wan et al., 2021). We determined that motion artifacts were not occurring during these recordings by performing recordings in a separate cohort of mice using a cre-dependent GFP control virus (Figure S7). This is necessary because the isosbestic control wavelength for this sensor has not yet been verified. Similar to the calcium responses observed in LHb_5HT2c_ and BNST_5HT2c_ neurons, 5-HT release onto these neurons was generally increased in response to social interaction and acoustic startle stimuli but decreased during water and alcohol consumption (Figure 1W-MM). The only difference in 5-HT release between males and females during these tasks was observed in the BNST_5HT2c_ responses to water consumption (Figure 1KK, LL), where females displayed a stronger reduction in 5-HT release. Similar to calcium signals in these neurons, BNST_5HT2c_ and LHb_5HT2c_ neurons displayed robust overlap of 5-HT signals during these behaviors (Figure 1W, X). Together, these experiments suggest that although BNST_5HT2c_ neurons are more responsive to affective stimuli in males, this phenomenon is largely not associated with sex differences at the level of 5-HT release onto these neurons.

To investigate the anatomy and neurochemical identity of the DRN-LHb and DRN-BNST projections, we next performed dual region retrograde tracing combined with immunohistochemistry for 5-HT (Figure 2A-D). In both males and females, a vast majority of DRN neurons projecting to the LHb and BNST were positive for 5-HT (Figure 2M, N). In addition, we observed that an extremely high percentage of DRN neurons project to both BNST and LHb (Figure 2O). We hesitated to directly compare percentages of these overlapping neurons between males and females, as differences in infection spread and uptake are challenging to normalize in these experiments. However, it may be the case that males have fewer DRN-BNST neurons that also project to the LHb than females (Figure 2P), and this should be investigated further in future studies using more quantitative tracing methods. Consistent with previous reports (Petit et al., 1995; Ren et al., 2018), the anatomical localization of these BNST- and LHb-projecting 5-HT neurons in the DRN was largely in dorsal and caudal regions. Subsequent *in-situ* hybridization experiments revealed that BNST_5HT2c_ neurons are primarily co-localized with the GABAergic marker vGAT, while LHb_5HT2c_ neurons are primarily co-localized with the glutamatergic marker vGlut2 (Figure 2Q-T). Notably, the percentages of BNST_5HT2c_/vGAT and LHb_5HT2c_/vGlut2 were comparable between sexes (Figure 2U-W).

**Figure 2:**
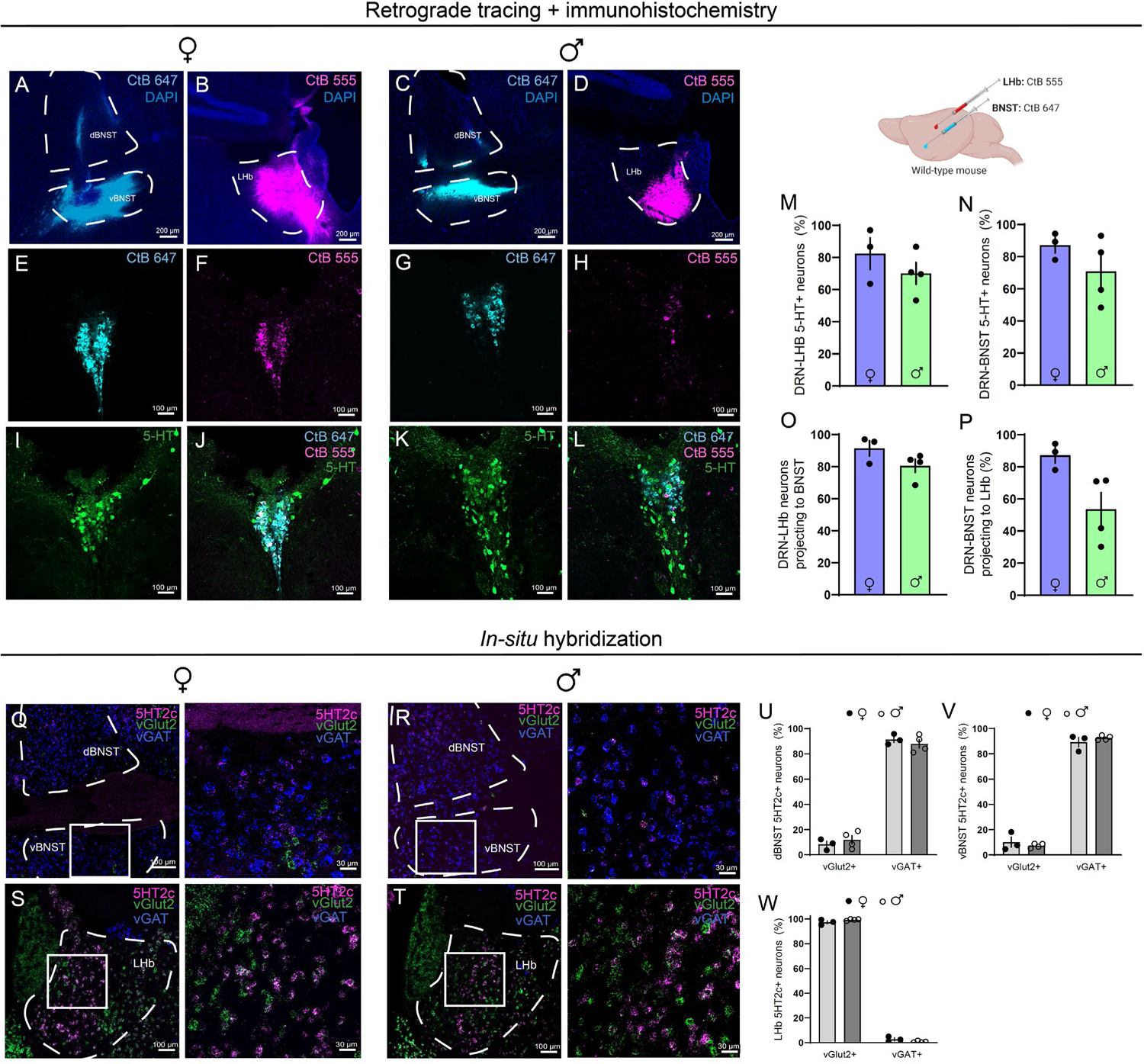
Anatomical and neurochemical characterization of DRN-LHb and DRN-BNST circuits. **A**, Representative CtB 647 infection in BNST, females. **B,** Representative CtB 555 infection, females. **C,** Representative CtB 647 infection in BNST, males. **D,** Representative CtB 555 infection in LHb, males. **E,** Representative CtB 647 labeling in DRN, females. **F,** Representative CtB 555 labeling in DRN, females.**G,** Representative CtB 647 infection in BNST, males. **H,** Representative CtB 555 infection, males. **I,** Representative 5HT labeling in DRN, females. **J,** Representative labeling in DRN for CtB 647, CtB 555, and 5HT, females. **K,** Representative 5HT labeling in DRN, males. **L,** Representative labeling in DRN for CtB 647, CtB 555, and 5HT, males. **M,** Percent of DRN-LHb neurons positive for 5HT (unpaired t-test, t(5)=1.07, p=0.3336). **N,** Percent of DRN-BNST neurons positive for 5HT (unpaired t-test, t(5)=2.275, p=0.2582). **O,** Percent of CtB 555 neurons that co-express CtB 647 in DRN (unpaired t-test, t(5)=1.171, p=0.1466). **P,** Percent of CtB 647 neurons that co-express CtB 555 in DRN. **Q,** *In-situ* hybridization for 5HT_2c_, vGAT, and vGlut2 in BNST, females. **R,** *In-situ* hybridization for 5HT_2c_, vGAT, and vGlut2 in BNST, males. **S,** *In-situ* hybridization for 5HT_2c_, vGAT, and vGlut2 in LHb, females. **T,** *In-situ* hybridization for 5HT_2c_, vGAT, and vGlut2 in LHb, males. **U,** Dorsal BNST 5HT2c neuron overlap with vGAT or vGlut2 (unpaired t-tests, vGlut2: t(5)=0.8020, p=0.4589; vGAT: t(5):0.7741, p=0.4739). **V,** Ventral BNST 5HT2c neuron overlap with vGAT or vGlut2 (unpaired t-tests, vGlut2: t(5)=0.337, p=0.4741; vGAT: t(5)=1.032, p=0.3942).**W,** LHb 5HT2c neuron overlap with vGAT or vGlut2 (unpaired t-tests, vGlut2: t(5)=1.676, p=0.1376; vGAT: t(5)=1.767, p=0.1376). n=3-4 mice/group, 2-4 slices/mouse. All data are represented as mean ± SEM. *p<0.05.

### Gq signaling in BNST_5HT2c_ and LHb_5HT2c_ modulates social behavior, arousal behavior, and binge alcohol consumption

Given that the 5HT_2c_ receptor primarily signals through Gq-coupled signaling mechanisms (Roth et al., 1998), we next asked how activation of Gq signaling in LHb_5HT2c_ and BNST_5HT2c_ would impact binge drinking, social, and arousal behavior in alcohol-naïve mice. 5HT_2c_-cre mice were injected in the LHb or BNST with a cre-dependent AAV encoding either the Gq-coupled DREADD hm3Dq or a mCherry control virus (Figure 3A, K) and three weeks later were tested in the 3-chamber social test, acoustic startle test, open field test, and sucrose preference test **before being tested in a binge alcohol consumption test**. To evaluate binge alcohol consumption, we employed the widely used Drinking in the Dark (DiD) model. In the DiD paradigm, animals are given free access to 20% alcohol in water for two hours on Monday-Wednesday and four hours on Thursday and this pattern is repeated for 3 weeks (cycles). While female mice generally consume greater amounts of alcohol than male mice in this model, both sexes voluntarily drink to achieve intoxication at blood alcohol concentrations (BACs) of at least 80 mg/dl (Radke et al., 2021). An acute intraperitoneal dose of 3 mg/kg clozapine-n-oxide (CNO) was administered to all mice 30 minutes prior to each individual behavioral task, while CNO treatment for the binge drinking test occurred only on the last day of DiD. We verified this CNO treatment significantly increased expression of the activity-dependent marker cFos in DREADD-treated animals (Figure S1). Critically, aside from a small effect on alcohol preference, CNO treatment alone in the absence of viral expression did not impact the behaviors tested (Figure S2). Activation of Gq signaling in BNST_5HT2c_ neurons did not affect social behavior in the 3-chamber social preference or social recognition tasks in males, but an interaction between sex and virus treatment was observed for social preference such that females displayed lower social preference in DREADD-treated animals compared water treated animals and males displayed higher social preference in DREADD-treated animals compared to water treated animals (Figure 3C, D). Activation of Gq signaling in BNST_5HT2c_ neurons increased acoustic startle responses in both sexes (Figure 3E-G) but reduced sucrose and alcohol drinking only in females (Figure 3H-J). DREADD-mediated activation of BNST_5HT2c_ neurons had no effect on open field exploratory or locomotor behavior in either sex (Figure S3). These findings indicate an important sex difference in the behavioral role of Gq signaling in BNST_5HT2c_ neurons: that it primarily promotes arousal in males, whereas it promotes arousal and reduces consumption of rewards in females.

**Figure 3:**
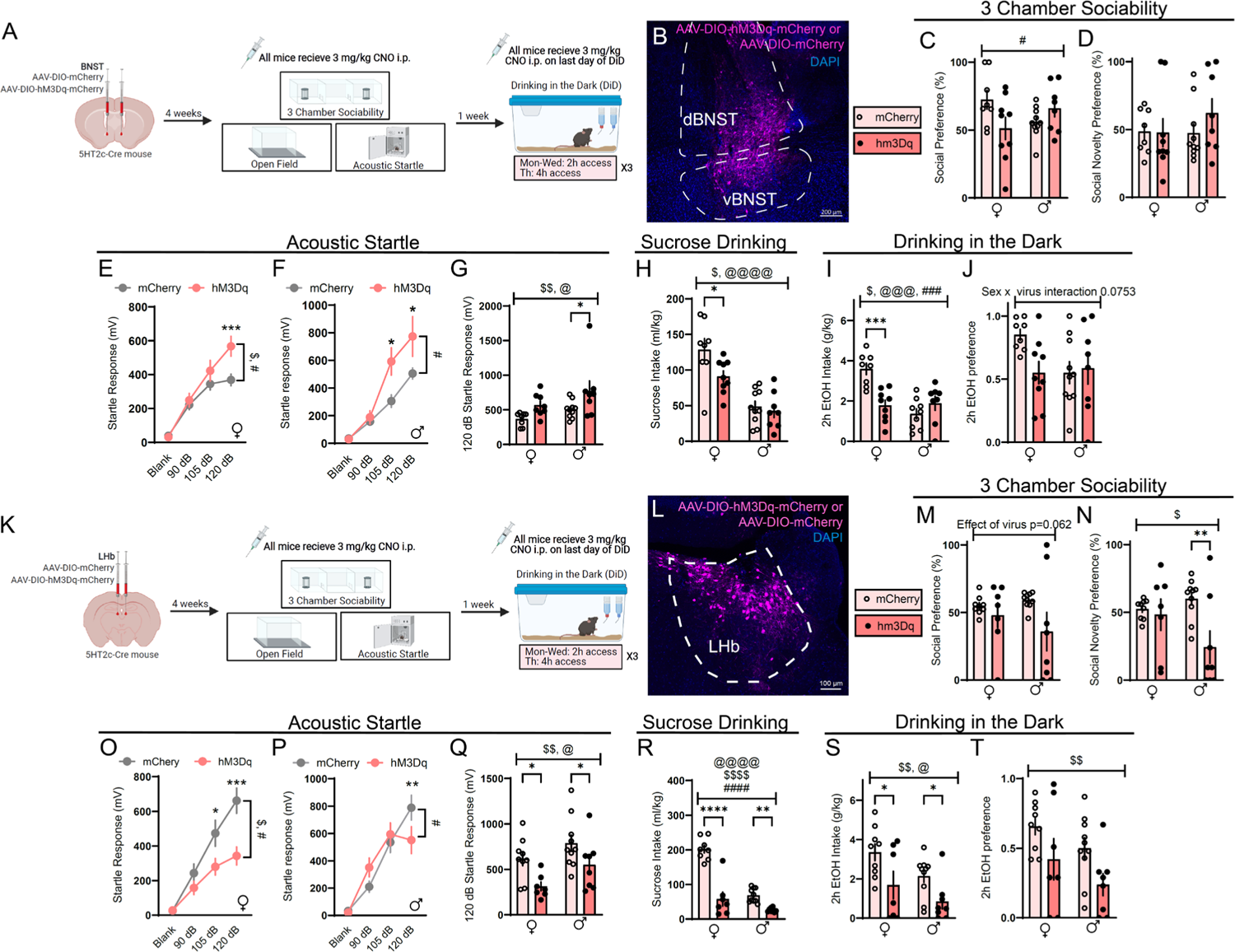
Chemogenetic activation of LHb_5HT2c_ or BNST_5HT2c_ neurons modulates affective behaviors and binge alcohol consumption. **A**, Surgical schematic and experimental timeline for BNST_5HT2c_ chemogenetic activation experiments. **B,** Representative viral infection in BNST. **C,** Social preference in the 3-chamber sociability test (two-way ANOVA, interaction: F(1,31)=6.103, p=0.0192, sex: F(1,31)=0.01504, p=0.9302, virus: F(1,31)=0.7581, p=0.3906). **D,** Social novelty preference in the 3-chamber sociability test (two-way ANOVA, interaction: F(1,31)=0.8310, p=0.3690, sex: F(1,31)=0.6015, p=0.4439, virus: F(1,31)=0.6529, p=0.4525). **E,** Acoustic startle behavior in females (two-way repeated measures ANOVA F(3,42)=4.244, p=0.0105, stimulus strength F(3,42)=75.06, p<0.0001, virus: F(1,14)=3.525, p=0.0814; Holm-Sidak post-hoc, mCherry vs. hm3Dq 120 dB p=0.0025). **F,** Acoustic startle behavior in males (two-way repeated measures ANOVA, interaction: F(3,48)=4.127, p=0.0111 stimulus strength: F(3,48)=51.95, p<0.0001, virus: F(1,16)=6.153, p=0.0246; Holm-Sidak post-hoc, mCherry vs. hm3Dq 105 dB p=0.0071, 120 dB p=0.0099). **G,** Acoustic startle behavior 120 dB only (two-way ANOVA, interaction: F(1,30)=0.1974, sex: F(1,30)=4.764, virus: F(1,30)=8.842; Holm-Sidak post-hoc, mCherry males vs. hm3Dq males p=0.0373). **H,** Sucrose consumption (two-way ANOVA, interaction: F(1,31)=2.375, p=0.1334 sex: F(1,31)=39.79, p<0.0001, virus: F(1,31)=4.661, p=0.0387, Holm-Sidak post-hoc females mCherry vs hm3Dq: p=0.0291). **I,** Alcohol intake in DiD (two-way ANOVA, interaction: F(1,30)=17.0, p=0.0003, sex: F(1,30)=14.14, p=0.0007, virus: F(1,30)=5.167, p=0.0303; Holm-Sidak post-hoc, females mCherry vs hm3dq: p=0.0.02). **J,** Alcohol preference in DiD (two-way ANOVA, interaction: F(1,31)=3.386, p=0.0753 sex: F(1,31)=2.113, p=0.1542, virus: F(1,31)=2.135, p=0.1541). **K,** Surgical schematic and experimental timeline for LHb_5HT2c_ chemogenetic activation experiments. **L,** Representative viral infection in the LHb. **M,** Social preference in the 3-chamber sociability test (two-way ANOVA, interaction: F(1,30)=1.069, p=0.3094, sex: F(1,30)=0.2553, p=0.6171, virus: F(1,30)=3.775, p=0.0621). **N,** Social novelty preference in the 3-chamber sociability test (two-way ANOVA, interaction: F(1,30)=3.876, p=0.0583, sex: F(1,30)=1.037, p=0.3166, virus: F(1,30)=6.198, p=0.0186; Holm-Sidak post-hoc, males mCherry vs males hm3dq: p=0.0316, males mCherry vs males hm3dq: p=0.0168). **O,** Acoustic startle behavior, females (two-way repeated measures ANOVA, interaction: F(3,42)=4.911, p=0.0051, stimulus strength F(3,42)=39.46, p<0.0001, virus: F(1,14)=9.377, p=0.0084; Holm-Sidak post-hoc, mCherry vs. hm3Dq, 105 dB: p=0.0162, 120 dB: p=0.0003). **P,** Acoustic startle behavior, males (two-way repeated measures ANOVA, interaction: F(3,48)=4.052, p=0.0120, stimulus strength: F(3,48)=51.74, p<0.0001, virus: F(1,16)=0.03768, p=0.8485). **Q,** Acoustic startle behavior, 120 dB only (two-way ANOVA, interaction: F(1,30)=0.1678, p=0.6850, sex: F(1,30)=5.828, p=0.0221, virus: F(1,30)=10.35, p=0.0031; Holm-Sidak post-hoc, females hM3dq vs mCherry p= 0.0367, males hM3dq vs mCherry p=0.0494). **R,** Sucrose consumption (two-way ANOVA, interaction: F(1,29)=22.62, p<0.0001, sex: F(1,29)=57.49, p<0.0001, virus: F(1,29)=73.83, p<0.0001; Holm-Sidak post-hoc, females mCherry vs hm3dq: p>0.0001, males mCherry vs males hm3dq: p=0.0082). **S,** Alcohol consumption in DiD (two-way ANOVA, interaction: F(1,30)=0.1607, p=0.6913, sex: F(1,30)=5.185, p=0.0301, virus: F(1,30)=10.80, p=0.0026; Holm-Sidak post-hoc females mCherry vs hM3dq p=0.0333, males mCherry vs. hM3dq p=0.0438). **T,** Alcohol preference in DiD (two-way ANOVA, interaction: F(1,30)=0.02081, p=0.8863, sex: F(1,30)=3.542, p=0.0696, virus: F(1,30)=7.582, p=0.0099). n=7-10 mice/group. All data are presented as mean ± SEM. $ denotes effect of virus, @ denotes effect of sex, # denotes interaction (sex x virus or virus x stimulus strength), * denotes post-hoc effects.

While DREADD-mediated activation of LHb_5HT2c_ neurons primarily altered behaviors in a similar fashion to DREADD-mediated activation of BNST_5HT2c_ neurons, sex differences in these behavioral effects were not observed. Indeed, activation of LHb_5HT2c_ neurons reduced social preference, sucrose consumption, and alcohol consumption in both sexes (Figure 3K-T). However, contrary to what we observed in the BNST, activation of LHb_5HT2c_ neurons reduced responses in the acoustic startle test rather than enhanced them. These results are in agreement with an established literature describing a role for the LHb in mediating passive coping to threats (Andalman et al., 2019; Berger et al., 2018; Coffey et al., 2020), yet highlight an important divergence in the behavioral functions of Gq signaling in LHb_5HT2c_ versus BNST_5HT2c_ neurons in alcohol-naïve mice. Of further relevance in this context, we observed that chemogenetic activation of LHb_5HT2c_ neurons reduced exploratory behavior, but not locomotion, in an open field (Figure S3).

### Binge alcohol consumption produces sex-specific affective disturbances and increases 5-HT_2c_ receptor expression in the LHb

Our results thus far suggest that BNST_5HT2c_ and LHb_5HT2c_ neurons play similar, though in part sex-dependent, roles in modulating social, arousal, and consummatory behaviors in naïve mice. Next, we wanted to ask: how does an insult that disrupts social and arousal behaviors impact the BNST_5HT2c_ and LHb_5HT2c_ systems? Studies in humans and animal models suggest that a history of alcohol consumption reduces social behaviors and increases anxiety-like behaviors, which may involve alcohol-induced adaptations in LHb and/or BNST 5HT_2c_ receptor expression and signaling (Campbell et al., 2021; Fu et al., 2020; Marcinkiewcz et al., 2015; Umhau et al., 2011).To investigate this possibility, we subjected mice to three weeks of DiD, and then after one week of abstinence assessed their behavior in the 3-chamber sociability task, the open field test, and the acoustic startle test (Figure 4A). Notably, this behavioral battery was designed in consideration of robust binge drinking-induced dysregulation of facial emotion recognition and acoustic startle responses reported in humans (Hone et al., 2020; Lannoy et al., 2019; Lannoy et al., 2018; Leganes-Fonteneau et al., 2020; Tolomeo et al., 2021). Consistent with previous reports, both sexes reached significant levels of intoxication as determined by their BACs at 4h (Figure 4B-E). It should be noted that in the DiD paradigm, 4h BACs may not be fully reflective of peak intoxication, as mice tend to front-load their alcohol consumption, drinking the majority of their total volume in the first two hours (Wilcox et al., 2014). DiD had no impact on social preference, but there was an interaction between sex and DiD such that social recognition was blunted in DiD females but not DiD males (Figure 4F, G). On the other hand, DiD had no impact on startle in female mice but markedly increased startle in males (Figure 4L-N). DiD similarly reduced exploratory behavior in both sexes in the open field test, although this effect was not consistent across behavioral cohorts (Figure 4H-K, also see S8). When we re-tested these mice four weeks following their last DiD exposure, we observed a normalization of female social recognition deficits as well as male and female open field exploratory deficits (Figure S4). However, the enhancement of startle responses in male DiD mice persisted out to this four week time point. Remarkably, when we tested the same male mice again at eight weeks post-DiD exposure, we still observed an enhancement in their startle responses (Figure S4). These data indicate that binge alcohol consumption uniquely alters affective behaviors in males and females, with male startle behavior being enhanced for a period far exceeding that of the actual alcohol exposure.

**Figure 4:**
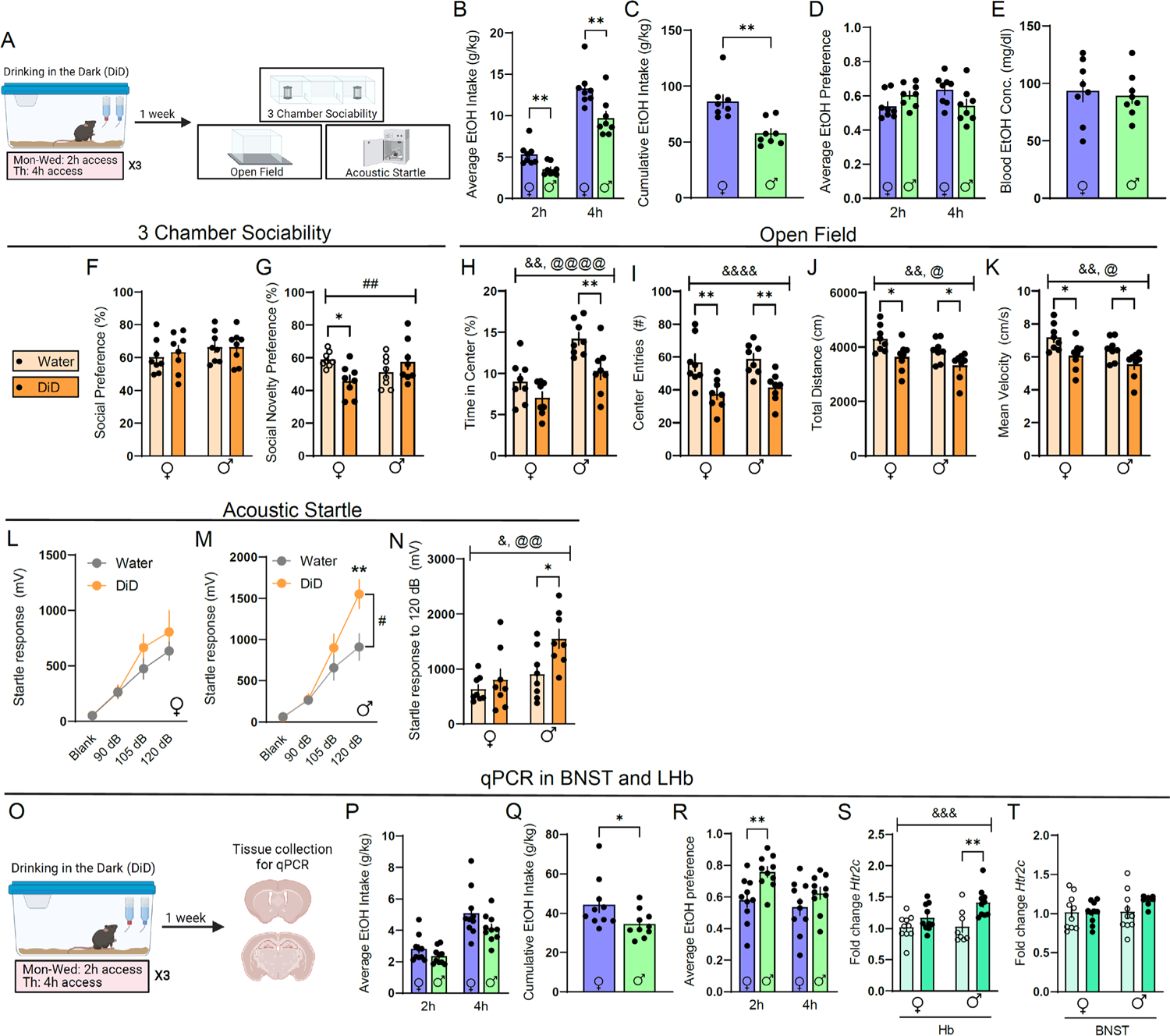
DiD induces unique affective disturbances in males and females. **A**, Experimental timeline for DiD behavioral studies. **B,** Average alcohol intake in DiD (unpaired t-tests, 2h: t(14)=3.140, p=0.0042, 4h: t(14)=3.166, p=0.0069). **C,** Cumulative alcohol intake in DiD (unpaired t-test, t(14)3.869, p=0.0017). **D,** Average alcohol preference in DiD (unpaired t-tests, 2h: t(14)=1.859, p=0.0841, 4h: t(14)=1.974, p=0.0685). **E,** Blood alcohol concentration immediately following 4h of DiD (unpaired t-test, t(14)=0.3529, p=0.7294). **F,** Social preference in the 3-chamber sociability test (two-way ANOVA, interaction: F(1,28)=0.1576, p=0.6944, sex: F(1,28)=1.567, p=0.2210, DiD: F(1,28)=0.1458, p=0.7054). **G,** Social novelty preference in the 3-chamber sociability test (two-way ANOVA, interaction: F(1,28)=8.165, p=0.008, sex: F(1,28)=0.3469, p=0.5606, DiD: F(1,28)=0.012, p=0.3231; Holm-Sidak post-hoc, water females vs. DiD females p=0.0214). **H,** Percent time in center of open field (two-way ANOVA, interaction: F(1,28)=1.257, p=0.2718, sex: F(1,28)=24.25, p<0.0001, DiD: F(1,28)=11.71, p=0.0019; Holm-Sidak post-hoc, male water vs. male DiD: p=0.0066). **I,** Center entries open field (two-way ANOVA, interaction: F(1,28)=0.5364, p=0.8185, sex: F(1,28)=0.6305, p=0.4228, DiD: F(1,28)=23.33, p<0.0001; Holm-Sidak post-hoc, female water vs. female DiD: p=0.0026, male water vs. male DiD: p=0.003). **J,** Total distance in open field (two-way ANOVA, interaction: F(1,28)=0.1195, p=0.7321, sex: F(1,28)=4.816, p=0.0366, DiD: F(1,28)=12.78, p=0.0013; Holm-Sidak post-hoc, female water vs. female DiD: p=0.0195, male water vs. male DiD: p=0.0302). **K,** Mean velocity in open field, (two-way ANOVA, interaction: F(1,28)=0.1191, p=0.07326, sex: F(1,28)=4.817, p=0.0366, DiD F(1,28)=12.77, p=0.0013; Holm-Sidak post-hoc water females vs. DiD females p=0.0195, water males vs. DiD males p=0.0302). **L,** Acoustic startle behavior, females (two-way repeated measures ANOVA, interaction: F(3,42)=1.096, p=0.3616, stimulus strength: F(3,42)=37.42, p<0.0001, DiD: F(1,14)=0.7868, p=0.3901). **M,** Acoustic startle behavior, males (two-way repeated measures ANOVA, interaction: F(3,42)=4.848, p=0.0055, stimulus strength: F(3,42)=58.67, p<0.0001, DiD: F(1,14)=2.944, p=0.0821; Holm-Sidak post-hoc water vs DiD p=0.0013). **N,** Acoustic startle behavior, 120 dB only (two-way ANOVA, interaction: F(1,28)=2.173, p=0.1516, sex: F(1,28)=10.28, p=0.0034, DiD: F(1,28)=6.514, p=0.0164; Holm-Sidak post-hoc, female water vs male DiD: p=0.0021, female DiD vs. male DiD: p=0.0128, male water vs male DiD: p=0.0323). **O,** Experimental timeline for DiD qPCR studies. **P,** Average alcohol intake in DiD (unpaired t-tests, 2h: t(18)=1.470, p=0.1589; 4h: t(18)=1.728, p=0.1012). **Q,** Cumulative alcohol intake in DiD (unpaired t-test, t(18)=2.146, p=0.0458). **R,** Average alcohol preference in DiD (unpaired t-tests, 2h: t(18)=3.161, p=0.0054, 4h: t(18)=1.342, p=0.1964). **S,** Relative expression of Htr2c mRNA in Hb (two-way ANOVA, interaction: F(1,35)=2.833, p=0.1012, sex: F(1,35)=3.037, p=0.0902, DiD: F(1,35)=14.63, p=0.0005; Holm-Sidak post-hoc females water vs males DiD: p=0.0019, males water vs males did: p=0.0025). **T,** Relative expression of *Htr2c* mRNA in BNST (two-way ANOVA, interaction: F(1,34)=1.157 p=0.2266, sex: F(1,34)=1.769, p=0.1923, DiD: F(1,34)=1.205, p=0.2801). All data are presented as mean ± SEM. & denotes effect of DiD, @ denotes effect of sex, # denotes interaction (sex x DiD or DiD x stimulus strength), and * denotes post-hoc effects.

To investigate whether abstinence from binge alcohol consumption is associated with altered expression of 5HT_2c_, we next subjected a separate cohort of mice to the DiD paradigm and collected brain tissue punches of the LHb and the BNST for qPCR-based analysis of *Htr2c* mRNA expression (Figure 4O-T). While DiD did not alter *Htr2c* mRNA levels in the BNST (Figure 4T), it induced an increase of *Htr2c* mRNA levels in the LHb, which post-hoc analysis revealed was particularly pronounced in males (Figure 4S). Thus, affective disturbances induced by DiD are accompanied by increased expression of 5HT_2c_ in the LHb, at least at the mRNA level.

### Binge alcohol consumption enhances the intrinsic excitability of LHb_5HT2c_ neurons in both sexes

Previous studies have demonstrated that alcohol exposure can enhance the activity of neurons in the LHb and the BNST during subsequent abstinence (Fu et al., 2020; Marcinkiewcz et al., 2015). To investigate whether this is specifically the case for 5HT_2c_-containing neurons in these regions, we subjected 5HT_2c_-Ai9 reporter mice to DiD and performed whole-cell patch clamp electrophysiology in the LHb and the BNST of the same mice (Figure 5A). Of note, recordings in BNST_5HT2c_ neurons were performed in the ventral BNST, while recordings in LHb_5HT2c_ neurons were performed in both the medial and lateral aspects of the LHb. We did not observe any differences in resting membrane potential (RMP) of LHb_5HT2c_ neurons between water and DiD groups or between males and females (Figure 5E). However, female BNST_5HT2c_ neurons had on average higher RMPs than those of maless (Figure 5K). Rheobase (current required to elicit an action potential) (Figure 5G, M), action potential threshold (Figure 5H, N), or number of current-elicited action potentials of BNST_5HT2c_ neurons (Figure 5O, P) did not differ between groups. However, DiD induced a trend for a reduction in voltage-elicited current (p=0.07) selectively in females that was particularly pronounced at higher voltages (Figure 5S). For LHb_5HT2c_ neurons, males displayed an increased number of current-elicited action potentials in DiD mice compared to water mice (Figure 5J). Although females did not display differences in rheobase between water and DiD groups when considering the entire LHb, when only medial LHb neurons were included in the analysis it was decreased in DiD mice, indicating greater excitability (Figure 5U-V). Splitting male cells into medial vs. lateral sub-divisions did not yield region-specific differences (data not shown). Thus, females may display region-specific increases in LHb_5HT2c_ neuronal excitability, whereas males display these increases across LHb_5HT2c_ neurons. This region-specificity of DiD-induced adaptations in females may explain why we did not observe a strong effect of DiD on total levels of LHb *Htr2c* mRNA in females, as our tissue punches included the entirety of the LHb. Notably, there was no effect of DiD the proportions of tonically active, silent, intermittently active/bursting, or depolarization blocked cells (silent at >-40 mV) in either region (Figure 5W-Z).

**Figure 5:**
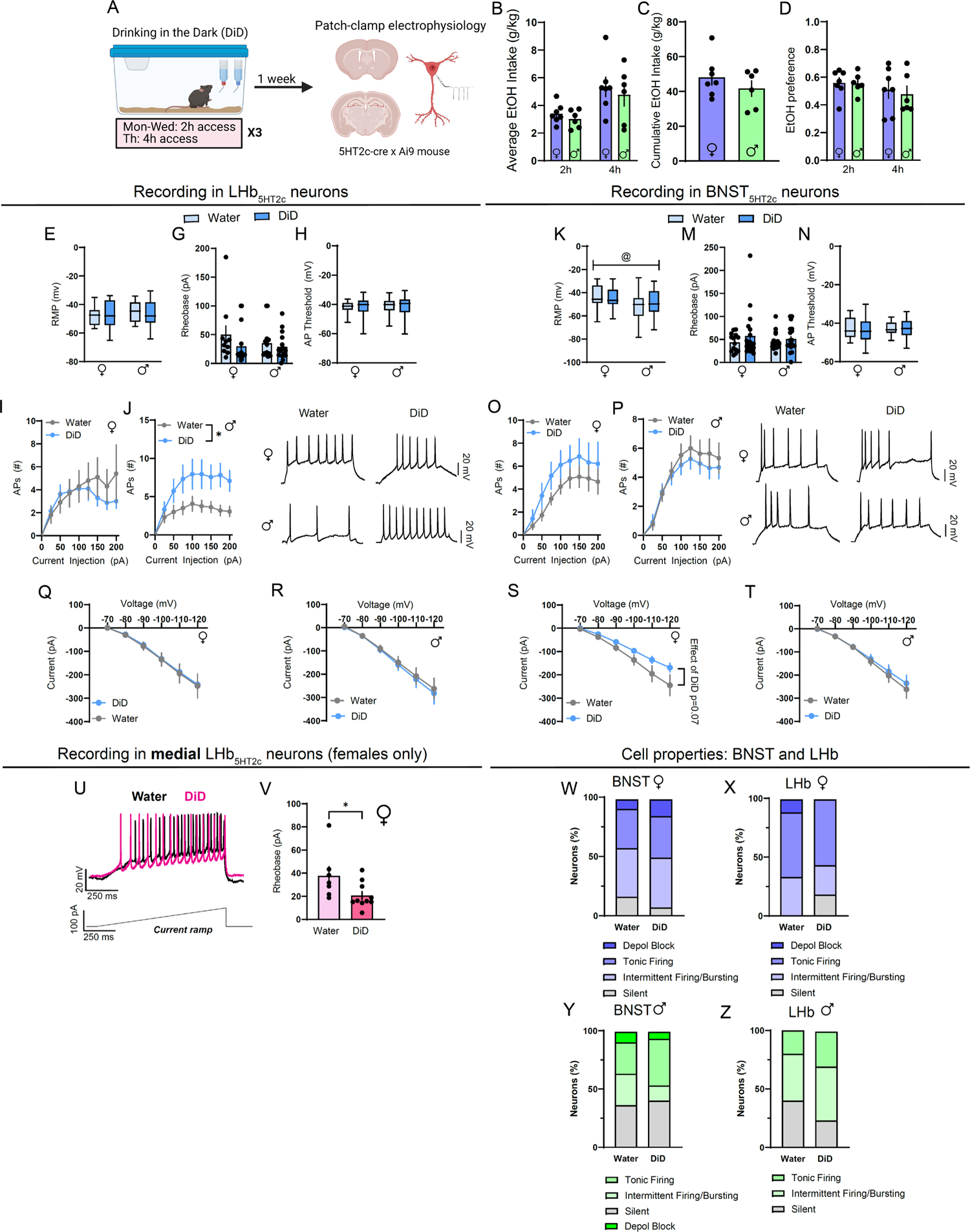
DiD induces physiological adaptations in LHb_5HT2c_ and BNST_5HT2c_. **A**, Experimental timeline for DiD electrophysiology studies. **B,** Average alcohol intake in DiD (unpaired t-tests, 2h: t(11)=1.009, p=0.3347, 4h: t(11)0.553, p=0.6043). **C,** Cumulative alcohol intake in DiD (unpaired t-test, t(11)=1.024, p=0.3278). **D,** Average alcohol preference in DiD (unpaired t-tests, 2h: t(11)=0.0738, p=0.9425, 4h: t(11)=0.3734, p=0.7159). **E,** Resting membrane potential (RMP) in LHb_5HT2c_ neurons (two-way ANOVA, interaction: F(1,57)=0.3589, p=0.5515, sex: F(1,57)=0.3115, p=0.5789, DiD: F(1,57)=0.0380, p=0.8455). **F,** LHb_5HT2c_ RMP vs. average alcohol intake in DiD (linear regression, F(1,10)=0.1769, R^2^=0.0174, p=0.683). **G,** LHb_5HT2c_ rheobase (two-way ANOVA, interaction: F(1,57)=0.5515, p=0.4757, sex: F(1,57)=1.11, p=0.2963, DiD: F(1,57)=1.964, p=0.1665). **H,** LHb_5HT2c_ action potential (AP) threshold (two-way ANOVA, interaction: F(1,57)=0.0851, p=0.7716, sex: F(1,57)=0.02227, p=0.8819 DiD: F(1,57)=0.01765, p=0.8948). **I,** LHb_5HT2c_ elicited APs, females (two-way repeated measures ANOVA, interaction: F(8,232)=1.301, p=0.2438, current: F(2.672, 77.49)=9.40, p<0.0001, DiD: F(1,29)=0.2629, p=0.6120). **J,** LHb_5HT2c_ elicited APs, males (two-way repeated measures ANOVA, interaction: F(8,240)=1.968, p=0.0512, current: F(2.483,74.76)=12.09, p<0.0001, DiD: F(1,30)=5.691, p=0.0236). **K,** Average RMP in BNST_5HT2c_ neurons (two-way ANOVA, interaction: F(1,67):0.6314, p=0.4297, sex: F(1.67)=5.777, p=0.0190, DiD: F(1.67)=0.08616, p=0.7700). **L,** BNST_5HT2c_ RMP vs. average alcohol intake (linear regression, F(1,11)=6.043, R^2^=0.3546, p=0.0318). **M,** BNST_5HT2c_ rheobase (two-way ANOVA, interaction: F(1,67)=0.2568, p=0.6140, sex: F(1,67)=0.08456, p=0.7721, DiD: F(1,67)=1.633, p=0.2057). **N,** BNST_5HT2c_ AP threshold (two-way ANOVA, interaction: F(1,67)=0.02599, p=0.8724, sex: F(1,67)=0.002352, p=0.9615, DiD: F(1,67)=0.5019, p=0.4811). **O,** BNST_5HT2c_ elicited APs, females (two-way repeated measures ANOVA, interaction: F(8,248)=0.4389, p=,0.8970, current: F(2.142,66.39)=19.98, p<0.0001, DiD: F(1.36)=1.049, p=0.3137). **P,** BNST_5HT2c_ elicited APs, males (two-way repeated measures ANOVA, interaction: F(8,288)=0.3469, p=0.9464, voltage: F(2.285,82.26)=32.94, p<0.0001, DiD: F(1,36)=0.3564, p=0.5542). **Q,** Voltage-Current plot for LHb_5HT2c_ females (two-way repeated measures ANOVA, interaction: F(5,115)=0.01066, p=0.99, current: F(5,115)=55.51, p<0.0001, DiD: F(1,23)=0.01647, p=0.8990). **R,** Voltage-Current plot for LHb_5HT2c_ males (two-way repeated measures ANOVA, interaction: F(1,150)=0.0789, p=0.994, voltage: F(5,150)=70.8, p<0.0001, DiD: F(1,30)=0.08315, p=0.7751). **S,** Current-Voltage plot for BNST_5HT2c_ females (two-way repeated measures ANOVA, interaction: F(5,120)=2.099, p=0.0701, voltage: F(5,120)=67.37, p<0.0001, DiD: F(1,24)=2.870, p=0.1032; Holm-Sidak posthoc −120 mV water vs. DiD p=0.0399). **T,** Current-Voltage plot for BNST_5HT2c_ males (two-way repeated measures ANOVA, interaction: F(5,180)=0.2865, p=0.92, voltage: F(5,180)=86.20, p<0.0001, DiD: F(1,36)=0.1527, p=0.6983). **U,** Representative trace in medial LHb_5HT2c_ during rheobase test. **V,** Quantification of rheobase values in medial LHb_5HT2c_ in females (student’s unpaired t-test, t(15)=2.180, p=0.0456). **V,** Breakdown of cell properties at RMP, BNST females. **W,** Breakdown of cell properties at RMP, LHb females. **Y,** Breakdown of cell properties at RMP, BNST males. **Z,** Breakdown of cell properties at RMP, LHb males. Depol block = silent at greater than −40 mV. For females, n=5-7 mice/group, 1-4 cells/mouse. For males, n=6-7/group and 2-4 cells/mouse. All data except for box plots are represented as mean ± SEM. For box plots, center line is the mean, box limits are 25^th^-75^th^ percentiles, whiskers are min to max. & denotes effect of DiD, @ denotes effect of sex, # denotes interaction (sex x DiD or DiD x current), and * denotes post-hoc effects.

### Binge alcohol consumption modulates responses of DRN-LHb_5HT2c_ and DRN-BNST_5HT2c_ circuits to affective stimuli

In our next set of experiments, we sought to determine if social and arousal behaviors impacted by binge alcohol consumption are accompanied by alterations in the responses of LHb_5HT2c_ and BNST_5HT2c_ neurons to relevant stimuli. Following their first round of recordings in an alcohol-naïve state, the same mice used in the fiber photometry experiments of Figure 1 were randomly assigned to DiD or water groups. After three weeks of DiD or water exposure followed by one week of abstinence, we again performed either GCaMP7f or GRAB-5HT recordings in LHb_5HT2c_ and BNST_5HT2c_ during behavior (Figure 6, S5, S6, S7, S8). DiD had no effect on female BNST_5HT2c_ or LHb_5HT2c_ calcium responses to social targets or acoustic startle stimuli (Figure 6E, F, I, J, M, N, Q, R). However, DiD increased male BNST_5HT2c_ and LHb_5HT2c_ calcium responses to acoustic startle stimuli (Figure 6O, P, S, T). In addition, DiD exposure was associated with the emergence of a negative LHb_5HT2c_ calcium response to alcohol in females, suggestive of a negative reinforcement signal in these neurons as a consequence of binge alcohol drinking experience (Figure S5). Interestingly, DiD had no impact on 5-HT release onto BNST_5HT2c_ or LHb_5HT2c_ neurons in males for any of the behaviors tested (Figure 6AA, BB, EE, FF, II, JJ, MM, NN; S6). Together with our electrophysiology findings suggesting increased intrinsic excitability of LHb_5HT2c_ neurons, these results indicate that the increased calcium responses to acoustic startle stimuli observed in DiD-exposed males are likely occurring post-synaptically and independently of acute fluctuations in 5-HT release. Females exposed to DiD, on the other hand, displayed a robust increase in 5-HT release onto BNST_5HT2c_ neurons in response to social interaction relative to water females (Figure 6CC, DD), suggesting that binge alcohol consumption alters the pre-synaptic component of this circuit in the female BNST. Together with our electrophysiology findings in BNST_5HT2c_, these results may suggest a model whereby alcohol consumption enhances 5-HT release onto BNST_5HT2c_, which then results in a compensatory blunting of BNST_5HT2c_ excitability to influence subsequent behavior in females. We were not able to detect any significant alterations in 5-HT release onto LHb_5HT2c_ neurons as a consequence of DiD for either sex in any of the behaviors tested (Figure 6, S7). Taken together, these findings suggest that subcortical 5-HT release may be more subject to alcohol-induced modulation in female mice.

**Figure 6:**
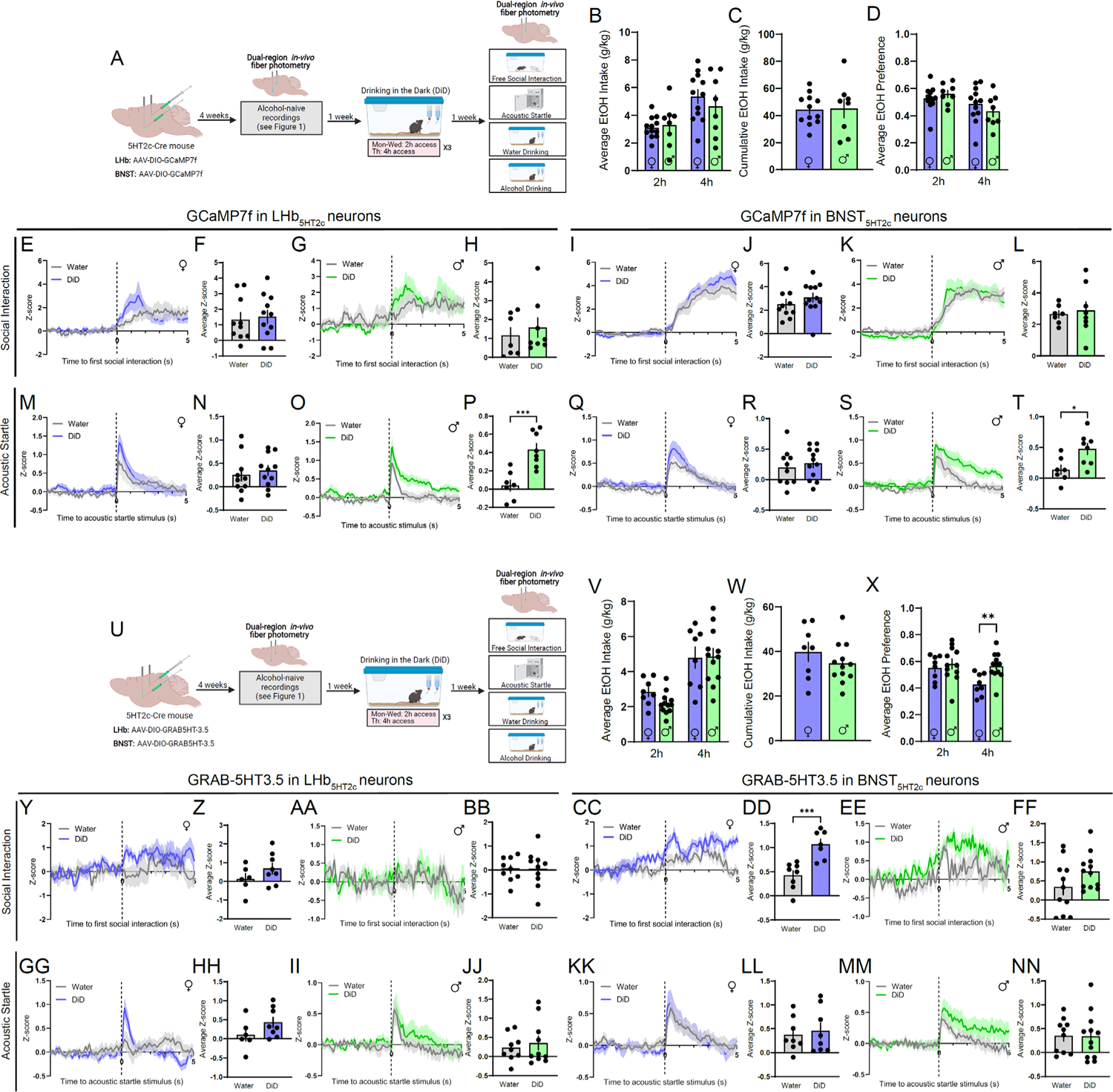
DiD modulates the responses of LHb_5HT2c_ and BNST_5HT2c_ neurons to affective stimuli. **A**, Surgical schematic and experimental timeline for LHb_5HT2c_ and BNST_5HT2c_ GCaMP fiber photometry experiments in DiD-exposed mice. **B,** Average alcohol consumption in DiD (unpaired t-tests, 2h: t(18)=0.3492, p=0.7310; 4h: t(18)=0.7865, p=0.4418). **C,** Cumulative alcohol intake in DiD (unpaired t-test, t(18)=0.1215, p=0.9047). **D,** Average alcohol preference in DiD (unpaired t-tests, 2h: t(18)=0.8352, p=0.4145; 4h: t(18)=0.9315, p=0.3639). **E,** Peri-event plot of LHb_5HT2c_ GCaMP activity during the first interaction with a novel social target, females (1 trial/mouse). **F,** Average z-score of LHb_5HT2c_ GCaMP signal for 0-5s post interaction, females (unpaired t-test, t(19)=0.2983, p=0.7687). **G,** Peri-event plot of LHb_5HT2c_ GCaMP activity during the first interaction with a novel social target, males (1 trial/mouse). **H,** Average z-score of LHb_5HT2c_ GCaMP signal for 0-5s post interaction, males (unpaired t-test, t(13)=0.6248, p=0.5429). **I,** Peri-event plot of BNST_5HT2c_ GCaMP activity during the first interaction with a novel social target, females (1 trial/mouse). **J,** Average z-score of BNST_5HT2c_ GCaMP signal for 0-5s post interaction, females (unpaired t-test, t(20)=0.9793, p=0.3391). **K,** Peri-event plot of BNST_5HT2c_ GCaMP activity during the first interaction with a novel social target, males (1 trial/mouse). **L,** Average z-score of BNST_5HT2c_ GCaMP signal for 0-5s post interaction, males (unpaired t-test, t(13)=0.3939, p=0.7001). **M,** Peri-even plot of LHb_5HT2c_ GCaMP activity during the acoustic startle test, females (average of 10 trials/mouse). **N,** Average z-score of LHb_5HT2c_ GCaMP signal for 0-5s post acoustic startle stimulus, females (unpaired t-test, t(19)=0.555, p=0.5854). **O,** Peri-event plot of LHb_5HT2c_ GCaMP activity during the acoustic startle test, males (average of 10 trials/mouse). **P,** Average z-score of LHb_5HT2c_ GCaMP signal for 0-5s post acoustic startle stimulus, males (unpaired t-test, t(13)=4.367, p=0.0008). **Q,** Peri-event plot of BNST_5HT2c_ GCaMP activity during the acoustic startle test, females (average of 10 trials/mouse). **R,** Average z-score of BNST_5HT2c_ GCaMP signal for 0-5s post acoustic startle stimulus, females (unpaired t-test, t(20)=0.5621). **S,** Peri-event plot of BNST_5HT2c_ GCaMP activity during the acoustic startle test, males (average of 10 trials/mouse). **T,** Average z-score of BNST_5HT2c_ GCaMP signal for 0-5s post acoustic startle stimulus (unpaired t-test, t(13)=2.776, p=0.0157). **U,** Surgical schematic and experimental timeline for LHb_5HT2c_ and BNST_5HT2c_ GRAB-5HT fiber photometry experiments in DiD-exposed mice. **V,** Average alcohol consumption in DiD (unpaired t-tests, 2h: t(18)=2.069, p=0.0532; 4h: t(18)=0.0925, p=0.9273). **W,** Cumulative alcohol intake in DiD (unpaired t-test, t(18)=1.091, p=0.2897). **X,** Average alcohol preference in DiD (unpaired t-tests, 2h: t(18)=0.5740, p=0.5731; 4h: t(18)=3.312, p=0.0039). **Y,** Peri-event plot of LHb_5HT2c_ GRAB-5HT activity during the first interaction with a novel social target, females (1 trial/mouse). **Z,** Average z-score of LHb_5HT2c_ GRAB-5HT signal for 0-5s post interaction, females (unpaired t-test, t(12)=1.328, p=0.2089). **AA,** Peri-event plot of LHb_5HT2c_ GRAB-5HT activity during the first interaction with a novel social target, males (1 trial/mouse). **BB,** Average z-score of LHb_5HT2c_ GRAB-5HT signal for 0-5s post interaction, males (unpaired t-test, t(17)=0.0798, p=0.9379). **CC,** Peri-event plot of BNST_5HT2c_ GRAB-5HT activity during the first interaction with a novel social target, females (1 trial/mouse). **DD,** Average z-score of BNST_5HT2c_ GRAB-5HT signal for 0-5s post interaction, females (unpaired t-test, t(13)=4.376, p=0.0008). **EE,** Peri-event plot of BNST_5HT2c_ GRAB-5HT activity during the first interaction with a novel social target, males (1 trial/mouse). **FF,** Average z-score of BNST_5HT2c_ GRAB-5HT signal for 0-5s post interaction, males (unpaired t-test, t(21)=1.689, p=0.106). **GG,** Peri-even plot of LHb_5HT2c_ GRAB-5HT activity during the acoustic startle test, females (average of 10 trials/mouse). **HH,** Average z-score of LHb_5HT2c_ GRAB-5HT signal for 0-5s post acoustic startle stimulus, females (unpaired t-test, t(12)=1.534, p=0.151). **II,** Peri-event plot of LHb_5HT2c_ GRAB-5HT activity during the acoustic startle test, males (average of 10 trials/mouse). **JJ,** Average z-score of LHb_5HT2c_ GRAB-5HT signal for 0-5s post acoustic startle stimulus, males (unpaired t-test, t(17)=0.5353, p=0.5993). **KK,** Peri-event plot of BNST_5HT2c_ GRAB-5HT activity during the acoustic startle test, females (average of 10 trials/mouse). **LL,** Average z-score of BNST_5HT2c_ GRAB-5HT signal for 0-5s post acoustic startle stimulus, females (unpaired t-test, t(14)=0.4082, p=0.6893). **MM,** Peri-event plot of BNST_5HT2c_ GRAB-5HT activity during the acoustic startle test, males (average of 10 trials/mouse). **NN,** Average z-score of BNST_5HT2c_ GRAB-5HT signal for 0-5s post acoustic startle stimulus (unpaired t-test, t(21)=0.08344, p=0.9343). n=6-12 mice/group. All data represented as mean ± SEM. *p<0.05, **p<0.01, ***p<0.0001.

### Genetic deletion of LHb or BNST 5HT_2c_ receptors partially normalizes DiD-induced social and arousal disturbances

To directly investigate the role of BNST and LHb 5HT_2c_ receptors in affective behaviors of alcohol-exposed vs. alcohol-naïve mice, we performed site-specific deletion of 5HT_2c_ receptors. We injected *Htr2c^lox/lox^* mice in the BNST or the LHb with AAVs encoding cre fused to GFP or GFP alone. Expression of cre excises 5HT_2c_ receptor DNA from the genome of tranduced cells via recombination at loxP sites (Burke et al., 2016), while expression of GFP alone keeps 5HT_2c_ expression intact. Four weeks following surgery, half of the mice were subjected to DiD and half continued to drink only water. One week following the last DiD session, mice were subsequently assessed in the 3-chamber sociability test, acoustic startle test, sucrose consumption test, and open field test (Figures 7 and S10). Critically, in these experiments enough time was given between surgery and the start of DiD for genetic recombination to occur, and thus 5HT_2c_ expression was repressed prior to alcohol exposure (Figure 7, S10). With this in mind, we found that deletion of 5HT_2c_ in the BNST increased binge alcohol consumption in females, but had no effect in males (Figure S10). This is consistent with what we observed in our chemogenetic experiments, where acute engagement of Gq signaling in BNST_5HT2c_ neurons reduced alcohol consumption in alcohol-experienced females but not males (Figure 3). However, contrary to the findings of our chemogenetic experiments, deletion of 5HT_2c_ from the BNST had no impact on sucrose consumption in either sex (Figure S10). This suggests that in females BNST 5HT_2c_ specifically regulates alcohol consumption in females and not simply the consumption of rewarding liquids *per se*. Deletion of 5HT_2c_ in the BNST had opposing effects on social recognition in alcohol-exposed versus alcohol-naïve females such that there was a significant interaction between virus treatment and DiD exposure. Specifically, alcohol-naïve cre females displayed reduced social recognition compared to alcohol-naïve GFP females, but alcohol-exposed cre females displayed increased social recognition compared to alcohol-exposed GFP females (Figure 7M-N). Interestingly, this manipulation also significantly reduced social preference in females regardless of alcohol history. There was no effect of 5HT_2c_ deletion in the BNST on social behavior in males (Figure 7O-P) or on startle behavior in females (Figure 7Q-R). However, regardless of alcohol experience, BNST 5HT_2c_ deletion somewhat reduced startle behavior in males, although this effect was not statistically significant (Figure 7S-T). It is notable that we did not observe the typical increase in startle behavior in male GFP mice following DiD exposure here, but this may be explained by the relatively low levels of baseline alcohol consumption in Htr_2c_*^lox/lox^* mice compared to wild type C57BL6/J mice (see Figure 4B-D). Exploratory behavior in the open field and locomotion were largely unaffected by BNST 5HT_2c_ deletion, except for an interaction between virus treatment and DiD exposure on time spent in the center of the open field in females (Figure S8). Together, these results suggest that 5HT_2c_ receptor signaling in the BNST partially contributes to alcohol-induced affective disturbances in a sex-specific manner while also reducing binge alcohol consumption selectively in females.

**Figure 7:**
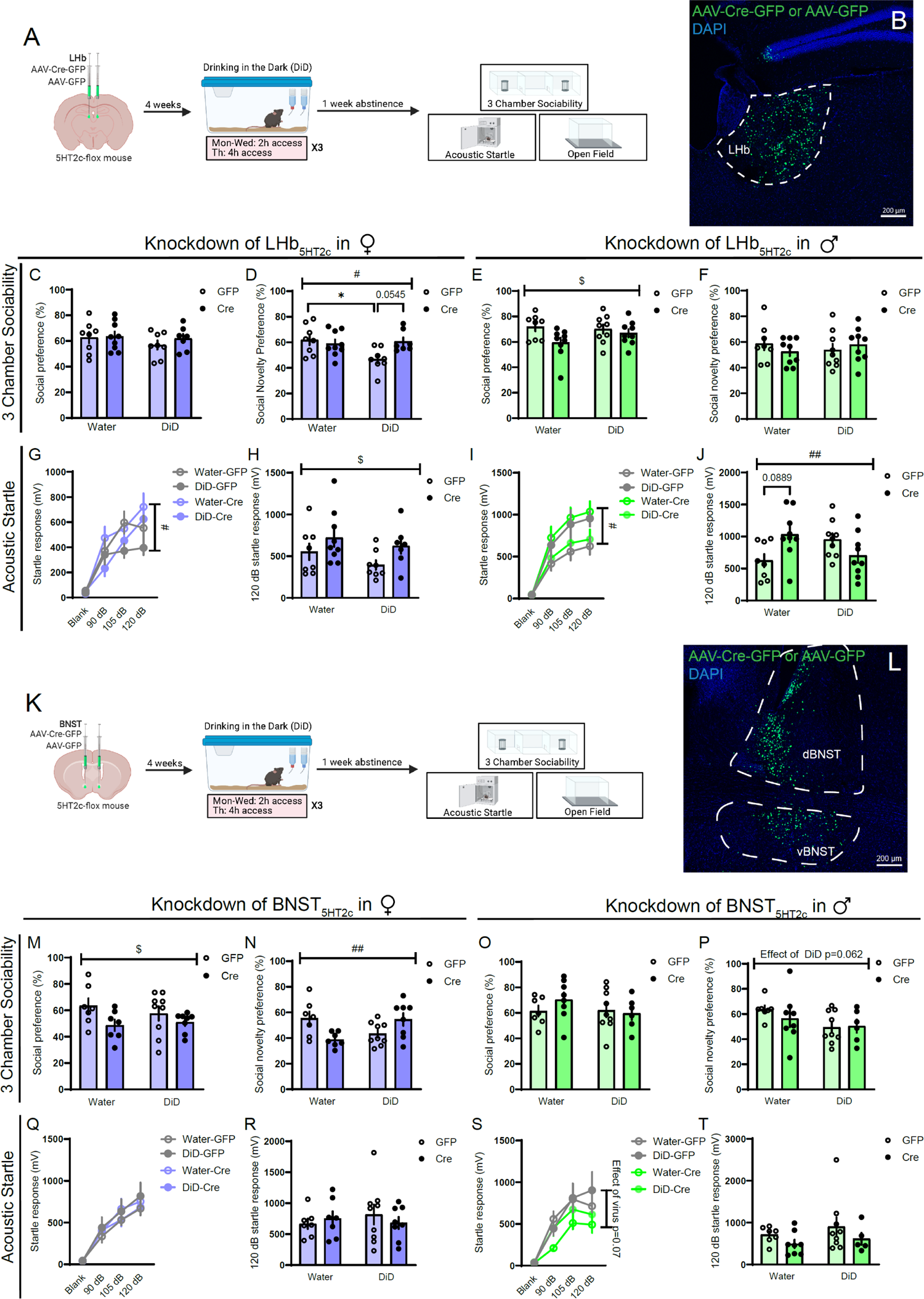
Deletion of 5HT_2c_ in the BNST or LHb partially normalizes alcohol-induced social and startle disturbances. **A**, Surgical schematic and experimental for LHb 5HT_2c_ deletion. **B,** Representative viral infection for LHb 5HT_2c_ deletion experiments. **C,** Social preference in the 3-chamber sociability test, females (two-way ANOVA, interaction: F(1,28)=0.3025, p=0.5867, DiD: F(1,28)=0.9430, p=0.3398, virus: F(1,28)=0.6131, p=0.4402). **D,** Social novelty preference in the 3-chamber sociability test, females (two-way ANOVA, interaction: F(1,28)=5.5877, p=0.0220, DiD: F(1,28)=3.631, p=0.0670, virus: F(1,28)=2.367, p=0.1352; Holm-Sidak posthoc water GFP vs. water GFP p=0.0277, water GFP vs. DiD cre p=0.0545). **E,** Social preference in 3-chamber sociability test, males (two-way ANOVA, interaction: F(1,31)=1.650, p=0.2085, virus: F(1,31)=0.6106, p=0.4405, virus: F(1,31)=4.958, p=0.0334). **F,** Social novelty preference in 3-chamber sociability test, males (two-way ANOVA, interaction: F(1,31)=1.518, p=0.2271, DiD: F(1,31)=0.01187, p=0.9131, virus: F(1,31)=0.0272, p=0.8701). **G,** Acoustic startle behavior, females (three-way repeated measures ANOVA, stimulus strength x virus: F(3,84)=3.394, p=0.0112, virus x DiD: F(1,28)=0.0007, p=0.9784, stimulus strength x virus x DiD: F(3,84)=2.260, p=0.0873, stimulus strength: F(3,84)=88.73, p<0.0001, virus: F(1,27)=0.7470, p=0.3948, DiD: F(1,28)=3.068, p=0.0908). **H,** Acoustic startle behavior, 120 dB only, females (two-way ANOVA, interaction: F(1,28)=0.1064, p=0.7467, DiD: F(1,28)=1.867, p=0.1827, virus: F(1,28)=4.558, p=0.0417). **I,** Acoustic startle behavior, males (three-way repeated measures ANOVA, stimulus strength x virus: F(3,93)=0.3065, p=0.8206, stimulus strength x DiD: F(3,93)=0.01384, p=0.9936, stimulus strength x DiD x virus: F(3,93)=4.391, p=0.0062, virus x DiD: F(1,31)= 7.432, p=0.0104, stimulus strength: F(3,93)=101.2, p<0.0001, DiD: F(1,31)=0.0005, p=0.9817, virus: F(1,31)=0.6014, p=0.4439). **J,** Acoustic startle behavior 120 dB only, males (two-way ANOVA, interaction: F(1,31)=8.744, p=0.0059, DiD: F(1,31)<0.0001, p=0.9980, virus: F(1,31)=5242, p=0.4745; Holm-Sidak posthoc water GFP vs. water cre p=0.0889). n=6-9/group. All data are presented as mean ± SEM. & denotes effect of DiD, $ denotes effect of virus, # denotes interaction (DiD x virus or virus x stimulus strength), * denotes post-hoc effects. **K,** Surgical schematic and experimental for BNST 5HT_2c_ deletion. **L,** Representative viral infection for BNST 5HT_2c_ deletion experiments. **M,** Social preference in the 3-chamber sociability test, females (two-way ANOVA, interaction: F(1,27)=0.7258, p=0.4017, DiD: F(1,27)=0.1379, p=0.7133, virus: F(1,27)=5.056, p=0.0329). **N,** Social novelty preference in the 3-chamber sociability test, females (two-way ANOVA, interaction: F(1,27)=11.90, p=0.0019, DiD: F(1,27)=0.2076, p=0.6523, virus: F(1,27)=0.4260, p=0.5195). **O,** Social preference in 3-chamber sociability test, males (two-way ANOVA, interaction: F(1,26)=1.154, p=0.2927, DiD: F(1,26)=0.9184, p=0.3647, virus: F(1,26)=0.3924, p=0.5365). **P,** Social novelty preference in 3-chamber sociability test, males (two-way ANOVA, interaction: F(1,26)=0.6155, p=0.4398, DiD: F(1,26)=3.794, p=0.0623, virus: F(1,26)=0.3552, p=0.5563). **Q,** Acoustic startle behavior, females (three-way repeated measures ANOVA, stimulus strength x virus: F(3,81)=0.2699, p=0.8470, virus x DiD: F(1,27)=0.7596, p=0.3911, stimulus strength x virus x DiD: F(3,81)=0.7636, p=0.5175, stimulus strength: F(3,81)=83.27, p<0.0001, virus: F(1,27)=0.00009, p=0.9924, DiD: F(1,27)=0.07816, p=0.7819). **R,** Acoustic startle behavior, 120 dB only, females (two-way ANOVA, interaction: F(1,27)=0.7727, p=0.3871, DiD: F(1,27)=0.1018, p=0.7521, virus: F(1,27)=0.04304, p=0.8372). **S,** Acoustic startle behavior, males (three-way repeated measures ANOVA, stimulus strength x virus: F(3,75)=1.695, p=0.1753, stimulus strength x DiD: F(3,75)=0.5282, p=0.6643, stimulus strength x DiD x virus: F(3,75)=1.176, p=0.3247, virus x DiD: F(1,25)= 0.4236, p=0.5211, stimulus strength: F(3,75)=58.08, p<0.0001, DiD: F(1,25)=0.8615, p=0.3622, virus: F(1,25)=3.583, p=0.07). **T,** Acoustic startle behavior 120 dB only, males (two-way ANOVA, interaction: F(1,25)=0.0464, p=0.8312, DiD: F(1,25)=0.9134, p=0.3484, virus: F(1,25)=2.475, p=0.1282). n=6-9/group. All data are presented as mean ± SEM. & denotes effect of DiD, # denotes interaction (DiD x virus), * denotes post-hoc effects.

Deletion of 5HT_2c_ in the LHb had only weak effects on binge alcohol consumption in DiD, with both sexes showing trends towards increased overall alcohol preference in cre mice (Figure S10) as well as interactions between viral treatment and day of DiD such that in later DiD sessions cre mice began to drink more than their GFP counterparts (Figure S10). However, there was no effect of LHb 5HT_2c_ deletion on sucrose consumption in either sex (Figure S10). Similar to the effects in BNST, there was an interaction between virus treatment and DiD exposure on social recognition in females, suggesting that 5HT_2c_ in the BNST and the LHb weakly promotes alcohol-induced disruptions in social recognition (Figure 7C-D). Curiously, deletion of LHb 5HT_2c_ reduced social preference in males regardless of alcohol history but had no effect on social recognition (Figure 7E-F). In females, deletion of LHb 5HT_2c_ increased startle behavior regardless of alcohol history, whereas in males there was an interaction between virus treatment and DiD exposure (Figure 7G-J). Specifically, 5HT_2c_ deletion in alcohol-naïve males increased startle behavior while in alcohol-exposed males it reduced startle behavior. Together, these results suggest that in both the BNST and the LHb, 5HT_2c_ does not appear to be the predominant mechanism by which alcohol induces behavioral dysregulation, as receptor deletion only partially normalized social and startle behaviors. Nevertheless, 5HT_2c_ in the BNST does appear to be a powerful regulator of alcohol intake in females.

### Chemogenetic inhibition of LHb_5HT2c_ fully normalizes DiD-induced social and arousal disturbances

To determine whether acute silencing of LHb_5HT2c_ or BNST_5HT2c_ could also modulate DiD-induced social and arousal disturbances, we performed chemogenetic inhibition of LHb_5HT2c_ or BNST_5HT2c_. We injected male and female 5HT_2c_-cre mice in the LHb or BNST with cre-dependent AAVs encoding either mCherry alone or the inhibitory DREADD hm4Di fused to mCherry. Four weeks following surgery, we split mice into water or DiD groups and let half of them consume alcohol freely for three cycles without engagement of hm4Di. On the last day of DiD, hm4Di and mCherry mice were treated with 3 mg/kg CNO prior to their drinking session. Interestingly, neither chemogenetic inhibition of BNST_5HT2c_ nor LHb_5HT2c_ impacted alcohol drinking in males. In females, however, chemogenetic inhibition of LHb_5HT2c_ increased alcohol intake (Figure S11). These effects in females were specific to alcohol, as chemogenetic inhibition of LHb_5HT2c_ did not impact sucrose consumption (Figure S11). Together with our 5-HT_2c_ deletion studies, our findings indicate that in the BNST, 5-HT_2c_ itself regulates alcohol consumption in females; in the LHb, however, the activity of 5-HT_2c_-containing neurons regulates alcohol consumption in females in a manner that may be only partly dependent on 5-HT_2c_ receptor signaling. Chemogenetic inhibition of BNST_5HT2c_ during the 3-chamber sociability test did not modulate deficits induced by DiD in females (Figure 8N), nor did it impact social behavior in males (Figure 8O-P). However, social novelty preference was enhanced in LHb_5HT2c_ hm4Di males regardless of alcohol history (Figure 8F). Critically, inhibition of LHb_5HT2c_ fully rescued DiD-induced social recognition deficits in female mice, indicating that increased activity of LHb_5HT2c_ neurons drives DiD-induced social impairment. Inhibition of LHb_5HT2c_ also fully normalized startle behavior in DiD males. Together, these results identify excessive activation of LHb_5HT2c_ as a critical mechanism for promoting sex-specific behavioral disturbances induced by binge alcohol consumption. Unfortunately, our BNST_5HT2c_ chemogenetic inhibition experiments were more difficult to interpret in the context of startle, as we observed a DiD-induced increase in startle in females, but not males. While chemogenetic inhibition of BNST_5HT2c_ fully normalized startle behavior in DiD females, startle behavior was unaffected in males of either DiD or Water groups. Considered together with the results of our chemogenetic activation studies and our 5HT_2c_ deletion studies, these results may simply reflect a floor effect of BNST_5HT2c_ inhibition on startle behavior in cases when startle behavior is not enhanced by alcohol. The factors determining whether females develop enhanced startle behavior as a consequence of DiD should be investigated in future studies.

**Figure 8:**
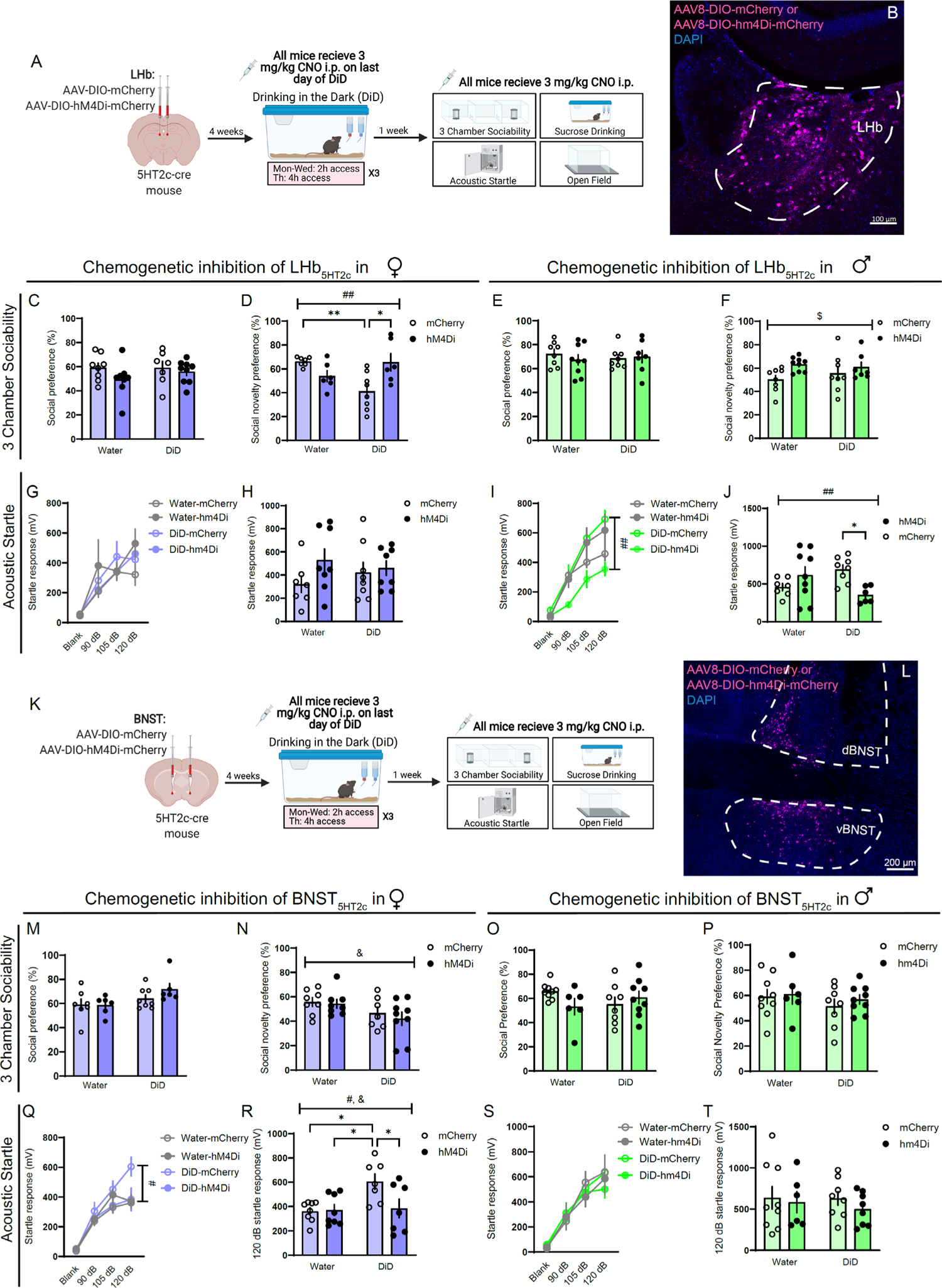
Chemogenetic inhibition of LHb_5HT2c_ and BNST_5HT2c_ normalizes DiD-induced social and arousal disturbances. **A**, Surgical schematic and experimental timeline. **B,** Representative image of viral infection in LHb. **C,** Social preference in the 3-chamber sociability test, LHb females (two-way ANOVA, interaction: F(1,23)=1.014, p=0.3244, DiD: F(1,23)=5.232, p=0.0317, virus: F(1,23)=0.8160, p=0.3757). **D,** Social novelty preference in the 3-chamber sociability test, LHb females (two-way ANOVA, interaction: F(1,23)=12.83, p=0.0016, DiD: F(1,23)1.723, p=0.2023, virus: F(1,23)=1.410, p=0.2472; Holm-Sidak test for multiple comparisons water mCherry vs. DiD mCherry p=0.0077, DiD mCherry vs. DiD hm4di p=0.0116). **E,** Social preference in 3-chamber sociability test, LHb males (two-way ANOVA, interaction: F(1,28)=0.6281, p=0.4347, DiD: F(1,28)=0.02297, p=0.088, virus: F(1,28)=0.1749, p=0.6790). **F,** Social novelty preference in the 3-chamber sociability test, LHb males (two-way ANOVA, interaction: F(1,28)=0.9090, p=0.3486, DiD: F(1,28)=0.1287, p=0.7225, virus: F(1,28)5.239, p=0.0298). **G,** Acoustic startle behavior, LHb females (three-way repeated measures ANOVA, stim strength: F(3, 69)=25.27, p>0.0001, DiD: F(1,23)=0.02758, p=0.8696, virus: F(1,23)=0.07716, p=0.7837, stim strength x DiD: F(3,69)=0.9065, p=0.4426, stim strength x virus: F(3,69)=0.7932, p=0.5018, DiD x virus: F(1,23)=0.9761, p=0.3334, stim strength x DiD x virus: F(3,69)=1.036, p=0.3823). **H,** Acoustic startle behavior 120 dB only, LHb females (two-way ANOVA, interaction: F(1,23)=0.2752, p=0.6049, DiD: F(1,23)=0.4492, p=0.5094, virus: F(1,23)=0.6380, p=0.4326). **I,** Acoustic startle behavior, LHb males (three-way repeated measures ANOVA, stim strength: F(3,78)=82.19, p<0.0001, DiD: F(1,26)=0.5068, p=0.4829, virus: F(1,26)=2.580, p=0.1033, stim strength x DiD: F(3,78)=0.9405, p=0.4253, stim strength x virus: F(3,78)=0.8116, p=0.4913, DiD x virus: F(1,26)=10.95, p=0.0028, stim strength x DiD x virus: F(3,78)=4.755, p=0.0043). **J,** Acoustic startle behavior 120 dB only, LHb males (two-way ANOVA, interaction: F(1,26)9.745, p=0.0044, DiD: F(1,26)=0.0319, p=0.8596, virus: F(1,26)=0.23=63, p=0.2713; Holm-Sidak test for multiple comparisons DiD mCherry vs. DiD hm4Di p=0.0438). **K,** Surgical schematic and experimental timeline. **L,** Representative image of viral infection in BNST. **M,** Social preference in 3-chamber sociability test, BNST females (two-way ANOVA, interaction: F(1,28)=0.5537, p=0.4630, DiD: F(1,28)=00.5258, p=0.4744, virus: F(1,28)=2.218, p=0.1476). **N,** Social novelty preference in 3-chamber sociability test, BNST females (two-way ANOVA, interaction: F(1,28)=0.1255, p=0.7258, DiD: F(1,28)=5.582, p=0.0253, virus: F(1,28)=0.4324, p=0.5162). **O,** Social preference in the 3-chamber sociability test, BNST males (two-way ANOVA, interaction: F(1,28)=3.310, p=0.0796, DiD: F(1,28)=0.0902, p=0.7661, virus: F(1,28)=0.5163, p=0.4784). **P,** Social novelty preference in 3-chamber sociability test, BNST males (two-way ANOVA, interaction: F(1,28)=0.0942, p=0.7612, DiD: F(1,28)=1.093, p=0.3047, virus: F(1,28)=0.4362, p=0.5143). **Q,** Acoustic startle behavior, BNST females (three-way repeated measures ANOVA, stim strength: F(2,78)=118.2, p<0.0001, virus: F(1,26)=2.602, p=0.1188, DiD: F(1,26)=1.113, p=0.3012, stim strength x virus: F(3,78)=3.156, p=0.0294, stim strength x DiD: F(3,78)=2.596, p=0.0583, virus x DiD: F(1,26)=4.396, p=0.0459, stim strength x DiD x virus: F(3,78)=2.808, p=0.045). **R,** Acoustic startle behavior 120 dB only, BNST females (two-way ANOVA, interaction: F(1,26)=4.481, p=0.044, DiD: F(1,26)=5.582, p=0.0259, virus: F(1,26)=3.744, p=0.0640; Holm-Sidak test for multiple comparisons water mCherry vs. DiD mCherry p=0.0232, water hm4Di vs. DiD mCherry p=0.0265, DiD mCherry vs. DiD hm4Di p=0.0398). **S,** Acoustic startle behavior, BNST males (three-way repeated measures ANOVA, stim strength: F(3,81)=57.59, p<0.0001, DiD: F(1,27)=0.000017, p=0.9897, virus: F(1,27)=0.3338, p=0.5682, stim strength x DiD: F(3,81)=0.2623, p=0.8254, stim strength x virus: F(3,81)=0.9185, p=0.4358, DiD x virus: F(1,27)=0.00729, p=0.9326, stim strength x DiD x virus: F(3,81)=0.2617, p=0.8528). **T,** Acoustic startle behavior 120 dB only, BNST males (two-way ANOVA, interaction: F(1,27)=0.1176, p=0.7343, DiD: F(1,27)=0.1697, p=0.6836, virus: F(1,27)=0.6640, p=0.4223). For all panels, n=6-9 mice/group. All data are presented as mean ± SEM. & denotes effect of DiD, # denotes interaction (DiD x virus), $ denotes effect of virus, * denotes post-hoc effects.

## Discussion

Previous anatomical and functional evidence suggests that the DRN is comprised of parallel sub-systems that differ in their input-output connectivity, physical localization within the DRN, neurotransmitter co-release properties, and roles in behavior (Commons, 2020; Gagnon and Parent, 2014; Huang et al., 2019; Muzerelle et al., 2016; Okaty et al., 2020; Ren et al., 2018; Sengupta and Holmes, 2019). Using a combination of anatomy and *in-vivo* biosensor measurement of 5-HT release, we found that many single 5-HT neurons in the caudal dorsal DRN target both the BNST and the LHb and appear to have coordinated 5-HT release in these regions. We demonstrate that this release is associated with increased calcium signaling in BNST_5H2c_ and LHb_5HT2c_ populations. Chemogenetic stimulation of BNST_5HT2c_ and LHb_5HT2c_ neurons in alcohol-naïve mice revealed many common functions of these neurons in promoting dysregulation of social and arousal behaviors. Perhaps reflecting previous data supporting the BNST as a sexually dimorphic structure (Lebow and Chen, 2016), a number of sex differences were observed in the behavioral functions of BNST_5HT2c_ neurons. We also found that binge alcohol consumption, which is associated with the development of negative affective states during abstinence, alters DRN-BNST_5HT2c_ and DRN-LHB_5HT2c_ *in-vivo* circuit physiology distinctly in each sex, with males displaying enhanced calcium signaling in both BNST_5HT2c_ and LHb_5HT2c_ neurons in response to acoustic startle stimuli, and females displaying robust enhancement of 5-HT release onto BNST_5HT2c_ neurons in response to novel social targets. Furthermore, DiD was associated with increased expression of LHb *Htr2c* mRNA and LHb_5HT2c_ *ex-vivo* excitability, particularly in males. DiD also modestly reduced BNST_5HT2c_ excitability in females. Genetic deletion of 5-HT_2c_ in the LHb, and to a lesser extent the BNST, partially normalized social and startle disturbances induced by DiD. However, chemogenetic inhibition of LHb_5HT2c_ neurons fully rescued sex-specific social and startle disturbances, indicating that 5-HT_2c_ itself may only partially contribute to excessive activation of LHb_5HT2c_ following alcohol exposure (see visual summary of findings in Figure S12).

Our results suggest that LHb_5HT2c_ neurons and BNST_5HT2c_ neurons are activated by aversive stimuli (acoustic startle stimuli) and inhibited by rewarding stimuli (water or alcohol consumption) in both males and females. One exception to this appears to be interaction with a novel juvenile social target, which is purported to be a rewarding stimulus for both adult males and adult females (Dölen et al., 2013; Venniro et al., 2018). Indeed, first contact with a juvenile social target in a neutral context (not the experimental mouse’s home cage) elicited activation in both LHb_5HT2c_ and BNST_5HT2c_ neurons, although this activation was greater in the BNST for males compared to females. Recent non cell-type specific fiber photometry studies performed in male mice indicate that LHb neurons are not notably modulated by non-aggressive interactions with familiar adult conspecifics (Wang et al., 2017), but *in-vivo* recordings of LHb neurons during interactions with juveniles have not been performed, nor have recordings in females.

To our knowledge, *in-vivo* recordings in the BNST have not previously been performed during non-aggressive social interactions in either sex. Crucially, for both sexes the degree of BNST_5HT2c_ neuronal activation in response to the first interaction with a novel juvenile social target was greater than that of any other stimulus tested, suggesting that this may be reflective of a salience-driven signal, as opposed to a valence-driven signal. This is consistent with a recent report illustrating that a large proportion of individual DRN-BNST neurons respond to a variety of valenced stimuli, including social stimuli, electric shock, and contextual threat (Paquelet et al., 2022). Only in our voluntary alcohol consumption fiber photometry experiments did we observe a notable divergence between patterns of 5-HT release onto LHb_5HT2c_ and BNST_5HT2c_ neurons and patterns of calcium activity in these neurons. In naïve females, alcohol consumption did not coincide with decreased calcium activity, yet 5-HT release was reduced during alcohol consumption (Figure 1). In males, however, both calcium activity and 5-HT release were reduced upon alcohol consumption. While the precise reasons for this are not clear, these results could reflect differences in post-synaptic 5-HT receptor signaling in females compared to males. More generally, these findings are in keeping with other studies supporting the BNST as a critical site for integration of salience and value, and extend these prior findings by demonstrating the *in-vivo* dynamics of 5-HT release and calcium activity during behavior.

Our chemogenetic activation studies in BNST_5HT2c_ neurons highlight important sex differences in the function of Gq signaling on affective behaviors and binge alcohol consumption. Chemogenetic activation of BNST_5HT2c_ neurons increased acoustic startle responses in both sexes, an effect that is in agreement with previous findings from our group showing that intra-BNST infusion of the 5HT_2c_ agonist mCPP increases acoustic startle responses in male mice (Mazzone et al., 2018). However, no other behavior tested was altered by chemogenetic activation of BNST_5HT2c_ neurons in males. Binge alcohol consumption, sucrose consumption, and to a lesser degree social preference, were reduced by chemogenetic activation of BNST_5HT2c_ neurons in females, indicating that these neurons play a more significant role in reward consumption in females compared to males. The potential mechanisms underlying these sex differences in reward consumption are numerous, but could be due to variations in the density or synaptic strength of downstream projections of BNST_5HT2c_ neurons, perhaps to regions like the ventral tegmental area or the DRN.

In contrast to the effects of chemogenetic activation of BNST_5HT2c_, chemogenetic activation of LHb_5HT2c_ did not produce sex-specific effects on affective behaviors. Instead, in both sexes, this manipulation had an inhibitory effect on liquid reward consumption (sucrose and alcohol), responses to acoustic startle stimuli, social behavior, and exploratory behavior. There was no inhibition of locomotion induced by chemogenetic activation of LHb_5HT2c_ neurons, indicating that these effects on affective behavior are not due to general behavioral inhibition. In regards to social behavior, these results are consistent with previous studies in males and extend the findings to females. Indeed, non-conditional chemogenetic activation of LHb neurons in male rats reduced social preference in the 3-chamber test in a recent study (Benekareddy et al., 2018). While no previous studies have investigated the effects of non-conditional chemogenetic activation of LHb neurons on alcohol-related behaviors, non-conditional chemogenetic inhibition of LHb neurons in male rats reduces withdrawal-related anxiety-like behavior (Kang et al., 2017) and voluntary alcohol consumption in an intermittent access model (Li et al., 2017). However, in a recent study employing chronic intermittent ethanol vapor followed by intermittent two-bottle choice access, non-conditional chemogenetic inhibition of LHb neurons in male rats did not impact alcohol drinking behaviors (Nentwig et al., 2022). Our results are broadly concordant with these previous findings and suggest that the LHb plays more of a role in behavior disruption induced by alcohol than alcohol consumption itself in males.

This study reveals an important distinction between the behavioral roles LHb_5HT2c_ neurons versus BNST_5HT2c_ neurons: LHb_5HT2c_ neurons serve to reduce active responses to stressors (reduced acoustic startle responses with chemogenetic activation), whereas BNST_5HT2c_ neurons serve to increase active responses to stressors (increased acoustic startle responses with chemogenetic activation). Despite these opposing behavioral functions, our fiber photometry experiments showed that the delivery of acoustic startle stimuli increased calcium activity and 5-HT release in BNST_5HT2c_ and LHb_5HT2c_ neurons. Thus, LHb_5HT2c_ and BNST_5HT2c_ neurons drive opposing behavioral responses to stressful stimuli, yet these neurons display similar cellular responses to these stimuli. In human patients, reduced responses to acoustic startle stimuli are observed in patients with major depression, while increased responses to these stimuli are observed in patients with anxiety disorders (Kaviani et al., 2004). Given that the LHb has been highly implicated in human major depression (Gold and Kadriu, 2019) and the BNST has been implicated in multiple sub-types of human anxiety disorders (Avery et al., 2016), it is then reasonable to speculate that regionally selective alterations of 5HT_2c_-containing neurons in LHb and BNST could play a role these behavioral observations.

Previous studies suggest that neuronal activity is enhanced in both the BNST and the LHb as a consequence of alcohol exposure in males, in at least a partly 5HT_2c_-dependent manner (Fu et al., 2020; Marcinkiewcz et al., 2015). However, no studies to date have investigated the effects of alcohol consumption on 5HT_2c_-mediated signaling in the LHb or the BNST in females. Withdrawal from intermittent access to alcohol in males increases cFos (a marker of neuronal activation) and 5HT_2c_ protein expression in the LHb, and this increase in cFos is reversed with intra-LHb infusion of a 5HT_2c_ antagonist (Fu et al., 2020). Consistent with these findings, we found that abstinence from binge alcohol drinking increases both the expression of LHb *Htr2c* mRNA as well as neuronal excitability. However, both of these effects were more pronounced in males compared to females. In the male BNST, withdrawal from alcohol vapor exposure also increases cFos activation that can be reversed with systemic treatment with a 5HT_2c_ antagonist (Marcinkiewcz et al., 2015). Furthermore, although not performed selectively in BNST_5TH2c_ neurons, slice electrophysiology results from this same study indicated that alcohol vapor exposure enhances the intrinsic excitability of BNST neurons in a 5HT_2c_-dependent fashion in males. However, our slice electrophysiology recordings yielded no differences in BNST_5HT2c_ neuronal excitability between water and DiD males. The discrepancy with previous studies in males could be the result of differences in degrees of intoxication achieved through voluntary DiD versus involuntary alcohol vapor exposure. In addition to displaying blunted voltage-current relationships compared to water females, DiD females had greater BNST_5HT2c_ RMPs than those of males. This is somewhat concordant with findings in BNST neurons expressing corticotrophin releasing factor (CRF), a subset of which express 5HT_2c_ (Marcinkiewcz et al., 2016b). In these neurons, binge alcohol consumption significantly increases the proportion of male cells that are in a depolarization block such that they begin to resemble the high proportions observed in females (Levine et al., 2021).

Our electrophysiology combined with our fiber photometry findings suggest that female mice may undergo only subtle changes in the physiological properties of BNST_5HT2c_ neurons as a consequence of binge alcohol consumption. However, binge alcohol consumption appears to strongly increase 5-HT release onto BNST_5HT2c_ neurons in females. This effect is stimulus-dependent, as the difference from water mice was only observed upon social interaction. Importantly, differences in stimulus-dependent activity between DiD and water mice were only observed for those behaviors which were sex-specifically disrupted as a consequence of DiD (see Figure 4). Males, which displayed increased startle behavior as a result of DiD, showed heightened calcium responses to startle but not to social interaction. Females, which displayed reduced social behavior as a result of DiD, showed enhanced *in-vivo* 5-HT responses to social interaction but not to startle. Taken together, these results suggest that female subjects could be more responsive to manipulations of 5-HT release to improve alcohol-induced behavioral deficits and highlight the importance of examining both male and female subjects in preclinical research.

Perhaps the most striking findings from our 5HT_2c_ deletion and chemogenetic inhibition experiments were those indicating that these manipulations could have opposing effects on social and arousal behaviors in alcohol exposed vs. alcohol naïve individuals. In females, deletion of 5HT_2c_ in the BNST or chemogenetic inhibition of BNST_5HT2c_ partly normalized social recognition in DiD mice but disrupted it in water mice. In males, deletion of 5HT_2c_ in the LHb or chemogenetic inhibition of LHb_5HT2c_ normalized startle behavior in DiD mice but enhanced it in in water mice. These data may suggest the existence of an inverted-U type relationship between levels of 5HT_2c_ expression/neural activity and specific affective behaviors in males and females such that too little 5HT_2c_/neural activation in the case of water cre-treated (5HT_2c_ deleted) mice or excessive 5HT_2c_/neural activation in the case of DiD GFP-treated (5HT_2c_ intact) mice dysregulates social and arousal processing.

Together with our chemogenetic experiments, our 5-HT_2c_ deletion experiments also highlight important dissociations between cellular manipulations of Gq/Gi signaling broadly and manipulations directly affecting 5-HT_2c_. For example, stimulation of Gq signaling in BNST_5HT2c_ neurons increased startle behavior in females and stimulation of Gi signaling reduced it, but deletion of BNST 5-HT_2c_ had no impact on startle behavior in females. Similarly, stimulation of Gq signaling in BNST_5HT2c_ neurons in females reduced sucrose consumption, but deletion of 5-HT_2c_ or stimulation of Gi signaling in BNST_5HT2c_ had no impact on sucrose consumption. Furthermore, stimulation of Gq or Gi signaling in LHb_5HT2c_ neurons had robust effects on alcohol consumption in females, but deletion of 5-HT_2c_ did not markedly affect this behavior. These data may suggest that in the LHb and BNST, 5-HT_2c_ itself plays less of an overall role in regulating social and arousal behaviors than the general activity of 5-HT_2c_-containing neurons.

There are several limitations of the present work that warrant discussion and/or further investigation. Previous work from our laboratory suggests that heavy alcohol exposure is associated with increased excitability of ventral, but not dorsal, BNST (vBNST, dBNST) neurons in male mice (Marcinkiewcz et al., 2015). Importantly, dorsal and ventral sub-regions of the BNST are known to be distinct in their circuit architecture and connectivity with regions outside of the BNST (Lebow and Chen, 2016). Therefore, we chose to perform our electrophysiology experiments specifically in the vBNST. While we also targeted our viral injections and optic fiber placements to the vBNST in subsequent photometry, receptor deletion, and chemogenetic experiments, we were not able to fully restrict viral expression to the vBNST and in some cases did observe spread to the dBNST. Thus, all animals had vBNST viral expression and some had both vBNST and dBNST viral expression. Future studies should aim to determine whether there is a functional dissociation between dBNST and vBNST 5HT2c-containing neurons, perhaps by selecting non-overlapping outputs of dBNST and vBNST sub-regions and injecting a retrograde virus to achieve sub-region-selective viral expression. Other important limitations of this study involve the design of our fiber photometry experiments. In order to facilitate recordings during voluntary alcohol consumption under tethered conditions (attached to a patch cord), we were forced to briefly water deprive experimental mice prior to testing. In an attempt to tease apart the rewarding effects of thirst quenching to the rewarding effects of alcohol, we also performed voluntary water drinking experiments under water deprivation conditions. However, it is possible that given increased thirst, the rewarding value of both water and alcohol were increased under these conditions. Whether the signals we observed would look different without prior water deprivation should be investigated in future studies, perhaps by performing more long-term home cage photometry recordings. Another potential caveat of our photometry experiments is that all mice were run through the battery of social, startle, and drinking behaviors twice, which could have impacted the results we observed in the second phase of testing. However, if this were the case, we would have expected to see changes in both DiD and Water groups. Rather, the photometry signals in Water mice during the second phase of testing closely resembled those obtained during the first phase, while DiD mice displayed marked differences from pre-DiD signals. The relatively long time period (1 month) between testing phases also reduces the likelihood that the post-DiD signals were impacted by pre-DiD testing.

In conclusion, our study suggests that the LHb and BNST represent two critical targets of an aversive DRN 5-HT sub-system that are physiologically impacted by binge alcohol consumption in sex-specific ways. Functionally, it appears that the primary mechanism by which alcohol promotes sex-specific expression of negative affect is through increased activation of LHb_5HT2c_, which is likely only partially 5HT_2c_-dependent. These data may have important implications for the development of novel, sex-specific treatments for AUD and comorbid mood disorders.

## Acknowledgements

This work was supported by grants from the National Institutes of Health’s (NIH) National Institute of Alcohol Abuse and Alcoholism (NIAAA) (M.F.: T32 AA007573-21; T.K.: R01 AA019454-12). Figures for this manuscript were created using BioRender.com. We also thank Dr. Dipanwita Pati for their feedback over the course of this project.

## Author Contributions

M.E.F. and T.L.K. conceptualized experiments. M.E.F. performed behavior, electrophysiology, fiber photometry, chemogenetics, histology, and analyzed data. L.H. assisted with behavioral experiments. H.L.H., M.M.P., and S.D. assisted with drinking experiments. K.M.B. performed *in-*situ hybridization experiments and qPCR. M.C. analyzed *in-situ* hybridization images. O.J.H. assisted with fiber photometry data collection and analysis. M.E.F. and T.L.K. wrote the paper with editing contributions from all authors.

## Online Methods

### Animals

Male and female wild-type C57BL6/J (Stock #: 000664, Jackson Laboratories), transgenic 5HT_2c_-Cre (provided by Dr. Laura Heisler’s lab, from Burke et al. 2016), transgenic 5HT_2c_ x Ai9 (Ai9 Stock #007909, 5HT_2c_-cre x Ai9 breeding performed in our animal facility), or transgenic *Htr_2c_^lox/lox^* (provided by Dr. Joel Elmquist, University of Texas Southwestern) adult mice aged 2-5 months at the start of the experiment were used as experimental animals. Male and female albino C57BL6/J (Stock #: 000058, Jackson Laboratories) adolescent mice aged 5-6 weeks were used as social targets. 5HT_2c_-Cre, and *Htr_2c_^lox/lox^*, and 5HT_2c_-Cre x Ai9 strains were bred in-house at UNC School of Medicine, while wild-type C57BL6/J and albino C5BL6/J mice were ordered from Jackson Laboratories and allowed to acclimate to our animal facility for one week prior to testing. Social target mice were group housed. Experimental mice were group housed for all experiments until the beginning of Drinking in the Dark (DiD), at which point they were single housed until experiment completion. All mice were housed in polycarbonate cages (GM500, Techniplast) under a 12:12h reverse dark-light cycle where lights turned off at 7:00 am. Mice had ad-libitum access to food (Prolab Isopro RMH 3000, LabDiet) and water unless otherwise stated. All experiments were approved by the UNC School of Medicine Institutional Animal Care and Use Committee (IACUC) and in accordance with the NIH guidelines for the care and use of laboratory animals.

### Drinking in the Dark (DiD)

Experimental mice were singly housed at least three days before initiation of the Drinking in the Dark (DiD) procedure and remained singly housed throughout the completion of the experiment. During this procedure, mice were given free access to both water and 20% (w/v) ethanol bottles in the home cage from 10:00 AM to 12:00 PM on Mondays, Tuesdays, and Wednesdays, and from 10:00 AM to 2:00 PM on Thursdays. At all other times, mice were given access to water alone. Water and ethanol bottles were weighed at the 2h time point on Mondays, Tuesdays, and Wednesdays, and at the 2h and 4h time point on Thursdays. Ethanol and water bottle positions were alternated daily to account for any inherent side preference in the animals. This weekly DiD access schedule was repeated for three weeks total, after which mice remained abstinent until behavioral testing. Confirmation of binge levels of intoxication was performed by measuring the blood ethanol concentration of tail blood collected at the end of a 4h DiD drinking session using the AM1 Analox Analyzer (Analox Instruments).

### 3-Chamber Sociability Test

The 3-chamber sociability test was performed using an apparatus consisting of a plexiglass rectangle measuring 20 cm x 40.5 cm x 22 cm that was divided into 3 equally-sized spaces by two panels with small square openings at the base to allow for movement between the chambers. Before the test began, social target mice were habituated for 10 minutes to a 10.5 cm diameter metal holding cage consisting of vertical bars with gaps between them. Importantly, mice outside of the holding cages can see, smell, and touch mice inside the holding cages but cannot freely interact wth them. Holding cages were placed in opposite corners of the two outermost chambers during all phases of the test (one in each of the two chambers). The test consisted of three 10 minute phases run one right after the other to total 30 minutes: 1. Habituation, 2. Social preference, and 3. Social novelty preference. For all phases, an experimental mouse was placed in the center chamber and allowed to move freely between chambers. In the habituation phase, the holding cages in the outermost chambers were empty. In the social preference phase, one holding cage contained a novel, same-sex, adolescent social target and the other holding cage contained a novel object (plastic toy mouse). In the social novelty preference phase, one holding cage contained the novel social target from the social preference phase (now familiar, on opposite side from social preference phase position) and the other holding cage contained a second novel social target. For all phases, the experimental mouse’s movements were tracked and the time spent interacting with the holding cages was measured using Ethovision XT 14.0 (Noldus). The social preference ratio was determined by calculating: (time spent with novel mouse)/(time spent with novel mouse + time spent with novel object) x 100 (for a % preference). The social novelty preference ratio was determined by calculating: (time spent with novel mouse)/(time spent with novel mouse + time spent with familiar mouse) x 100. Mice that did not enter both outermost chambers at least once were excluded from analysis for that phase of the test.

### Open Field Test

The open field test was performed in a plexiglass square arena with dimensions 50 cm x 50 cm x 40 cm. An experimental mouse was placed in the corner of the arena and allowed to freely explore for 10 minutes. Behavior was recorded with an overhead video camera. Mouse position data was acquired from videos using Ethovision XT (Noldus, Inc.). The total distance traveled, mean velocity, time spent in the 10×10 cm center zone, and number of entries to the center zone were calculated.

### Acoustic Startle Test

The acoustic startle test was performed with the SR-LAB Startle Response System (SD Instruments). The system consisted of a sound-attenuating isolation cabinet with dimensions 38 cm x 36 cm x 46 cm containing a small plastic cylindrical enclosure with dimensions 4 cm (diameter) x 13 cm (length). The isolation cabinet was lit by low intensity white light throughout the test. At the start of the test, an experimental mouse was placed in the plastic enclosure and allowed to habituate for 5 minutes. After habituation, the mouse was presented with one of four different acoustic stimuli ten different times (for a total of 40 trials): 90 dB, 105 dB, 120 dB, and 0 dB. Acoustic stimuli were delivered for 40 ms each trial and the startle response was measured for 200 ms following stimulus delivery. Startle stimuli were delivered in a random order with random inter-stimulus intervals lasting 30-50 seconds. For each mouse, maximum responses for each stimulus type were averaged across the ten trials.

### Home Cage Sucrose Consumption Test

The sucrose consumption test was performed in the home cage of singly-housed experimental animals. Mice were given 4hs of access to both water and a 2% (w/v) sucrose solution and bottles were weighed at the 2h and 4h time points. Sucrose consumption was normalized to body weight (ml/kg consumed).

### Stereotaxic Surgery

Adult mice (>7 weeks of age) were anesthetized with isoflurane (1-3%) in oxygen (1-2 L/min) and positioned in a stereotaxic frame using ear cup bars (Kopf Instruments). The scalp was sterilized with 70% ethanol and betadine and a vertical incision was made before using a drill to burr small holes in the skull directly above the injection targets. Using a 1 µl Neuros Hamilton Syringe (Hamilton, Inc.), viruses were then microinjected at a 0° angle into the LHb (mm relative to bregma: AP: −1.5, ML: ± 0.5, DV: −2.95) and/or the BNST (mm relative to bregma: AP: + 0.7, ML: ± 0.9, DV: −4.60) at a volume of 200 nl of virus per injection site. For fiber photometry experiments, optic fibers were implanted in the LHb and BNST at the same DV as the viral injections. Optic fibers were secured to the skull using Metabond dental cement (Parkell, Inc.). For chemogenetics experiments, 5HT_2c_-Cre mice were injected with either AAV8-hSyn-DIO-mCherry, AAV8-hSyn-DIO-hm3Dq-mCherry, or AAV8-hSyn-DIO-hm4Di-mCherry (Addgene). For 5HT_2c_ knockdown experiments, *Htr_2c_^lox/lox^* mice were injected with either AAV8-hSyn-GFP or AAV8-hSyn-Cre-GFP (UNC Vector Core). For fiber photometry experiments measuring calcium in 5HT_2c_ neurons, 5HT_2c_-Cre mice were injected with AAV8-hSyn-DIO-GCaMP7f (Addgene). For fiber photometry experiments measuring 5HT in 5HT_2c_ neurons, 5HT_2c_-Cre mice were injected with AAV9-hSyn-DIO-5HT3.5 (Dr. Yulong Li). For retrograde tracing experiments, wild type C57BL6/J mice were injected with both Cholera Toxin-B 555 (LHb) and Cholera Toxin-B 647 (BNST) (Thermo Fisher Scientific). Mice were given acute buprenorphine subcutaneously (0.1 mg/kg) on the day of surgery and access to Tylenol in water for 3 days post-op. Mice were allowed to recover in their home cages for at least 4 weeks before the start of experiments.

### Chemogenetic Activation of LHb_5HT2c_ or BNST_5HT2c_ neurons

Starting at least 4 weeks after stereotaxic surgery, LHb_5HT2c_ mCherry/hM3Dq and BNST_5HT2c_ mCherry/hM3Dq mice were subjected to a battery of behavioral tests performed in the following order: three chamber sociability, acoustic startle, and open field. One test was performed each day for 3 successive days, and both mCherry and hM3Dq groups received 3 mg/kg CNO intraperitoneally (i.p.) 30 minutes before the start of each behavioral test. One week after the open field test, mice were subjected to three weeks of DiD with no treatment (baseline). On the last day of DiD, both mCherry and hM3Dq mice were given 3 mg/kg CNO i.p. 30 minutes before a 4h drinking session. One week following DiD, mice were given one day of 4h access to either water or a 2% (w/v) sucrose solution with no treatment (baseline). The next day, both mCherry and hM3Dq mice were given 3 mg/kg CNO i.p. 30 minutes before a 4h drinking session where both water and a 2% sucrose solution were available. One week after sucrose testing, mice were perfused and viral placement was verified.

### Chemogenetic Inhibition of LHb_5HT2c_ or BNST_5HT2c_ neurons

Starting at least 4 weeks after stereotaxic surgery, half of LHb_5HT2c_ mCherry/hm4Di and BNST_5HT2c_ mCherry/hm4Di mice were subjected to three weeks of DiD. 30 mins before the last drinking session of the last day of DiD, mice were injected with 3 mg/kg CNO i.p. One week after the last DiD session, mice were subjected to three chamber sociability, acoustic startle, and open field testing. One test was performed each day for 3 consecutive days, and all mCherry and hm4Di mice were treated with 3 mg/kg CNO i.p. 30 minutes before the start of each behavioral test. One week after these tests, mice were subjected to a sucrose consumption test, again with both mCherry and hm4Di mice receiving acute CNO injections. Mice were then perfused and viral placement was verified.

### Viral-mediated genetic deletion of 5HT_2c_ in LHb and BNST

Starting 4 weeks after stereotaxic surgery, half of LHb_5HT2c_ GFP/Cre and BNST_5HT2c_ GFP/Cre mice were subjected to three weeks of DiD and half continued to have free access to only water. One week following the last DiD session, all mice were subjected to a battery of behavioral tests performed in the following order: three chamber sociability, acoustic startle, and open field. One test was performed each day for 3 days. One week following the open field test, mice were given one day of 4h access to either water or a 2% (w/v) sucrose solution. One week after sucrose testing, mice were perfused and viral placement was verified.

### Fiber Photometry

#### Hardware

Fiber photometry was performed using a commercially available system from Neurophotometrics, Inc. To record either GCaMP7f or GRAB-5HT signals in LHb_5HT2c_ and BNST_5HT2c_ neurons simultaneously, light from a 470 nm LED was bandpass filtered, collimated, reflected by a dichroic mirror, and focused by a 20x objective into a multi-branch patch cord. Excitation power was adjusted to obtain 75-120 uW of 470 nm light at the tip of the patch cord. Emitted fluorescence was then bandpass filtered and focused on the sensor of a CCD camera and images of the patch cord ROIs corresponding to LHb and BNST were captured at a rate of 40 Hz. 415 nm LED light was also delivered in a similar fashion alternatingly with 470 nm LED light to serve as an isosbestic control channel. To align photometry signals with mouse behaviors, the open-source software Bonsai was used to trigger LEDs simultaneously with behavioral video recording. In a subset of experiments (drinking and startle), TTL pulses were also sent to Bonsai during recordings to identify timestamps of relevant stimuli in real time through an Arduino-based setup.

#### Data Collection

Starting at least 4 weeks after stereotaxic surgery, mice were habituated to patch cords for at least 2 days prior to fiber photometry recordings. Ethanol-naïve male and female mice expressing either GCaMP7f or GRAB-5HT in LHb_5HT2c_ and BNST_5HT2c_ neurons were then subjected to a battery of behavioral tests in the following order: free social interaction, acoustic startle, water drinking, and ethanol drinking. One test was performed each day for 4 successive days, and fiber photometry signals were recorded throughout all behavioral tests. One week following the last behavioral test, half of these mice went through 3 weeks of DiD while the other half continued to only drink water. One week after the last DiD session, all mice were subjected to the same tests in the same order as in the pre-DiD behavioral battery and GCaMP7f or GRAB-5HT signals were recorded and compared between groups.

The free social interaction test was performed in a mouse polycarbonate shoebox cage with dimensions 19 cm x 29 cm x 13 cm (without bedding). Prior to testing, both experimental mice and same-sex, adolescent social targets were habituated to social interaction test cages for 10 minutes (separately). This habituation was done so that the shoebox cage would be a neutral, but not novel, territory for both mice. For the test, an experimental mouse was placed in the social interaction cage first, then one minute later a social target was added to the cage. The mice were then allowed to freely interact for 10 minutes and the interaction was recorded with an overhead video camera. The average z-scored ΔF/F was quantified for the 5 seconds before and the 5 seconds after the first social interaction with the juvenile target mouse. Timestamps for the start of the first social interaction were generated manually from videos.

Acoustic startle experiments were performed as described above, except that animals were exposed to only 0 and 120 dB acoustic stimuli rather than 0, 90, 105, and 120 dB acoustic stimuli. The average z-score was quantified for either the 5 seconds before and the 5 seconds after each trial stimulus or the 1 second before and 1 second after each trial stimulus and averaged across trial type for each mouse (n=10 blank trials and n=10 120 dB trials). TTL pulses corresponding to trial stimulus onset timestamps were delivered to Bonsai from the startle apparatus using an Arduino device.

For ethanol and water drinking experiments, an automated Arduino sipper device (Godynyuk et al. 2019) was used to deliver TTL pulses corresponding to lick timestamps to Bonsai. To habituate the mice to the sipper, a dummy sipper device was placed in the home cage of experimental animals one week prior to testing (mice must drink water from the sipper but the sipper was not hooked up to the Arduino device). On the day of testing, the sipper was placed into the home cage and mice were allowed to freely drink from it until they reached a criterion of at least 2 isolated drinking bouts more than 10 seconds apart and lasting at least 5 seconds each. Mice that did not drink from the sipper within 40 minutes were excluded from the experiment. For each test, one bottle of either water or 20% (w/v) ethanol was placed in the sipper (separate test days for water vs. ethanol). The average z-score was quantified for the 5 seconds before and 5 seconds after the start of each drinking bout and averaged across trials for each liquid.

#### Analysis

Fiber photometry signals were analyzed using a custom MATLAB (MathWorks, Inc.) script. Briefly, 470 nm and 415 nm signals were de-interleaved, background fluorescence was subtracted for each ROI, and the data was low-pass filtered at 2 Hz. Data was then fit to a biexponential curve and the fit was subtracted from the signal to correct for baseline drift. Next, ΔF/F (%) was calculated for 470 nm and 415 nm signals as 100*(signal-fitted signal)/(fitted signal). For GCaMP7f recordings, the 470 nm signal for the entire recording session was then z-scored and fit using non-negative robust linear regression and the 415 nm signal was fit to the resulting 470 nm signal. The fit 415 nm signal was next subtracted from the z-scored 470 nm signal to yield a motion-corrected recording. For GRAB-5HT recordings, the 470 nm signal was z-scored for the entire recording session without fitting and subtracting the 415 signal, as the 415 nm signal is not an appropriate isosbestic channel for this sensor. We performed additional recordings using AAV-DIO-GFP in 5HT2c-cre mice to ensure a lack of motion artifacts during the behaviors tested (See Figure S7).

#### Patch-clamp electrophysiology

Whole-cell patch clamp electrophysiology recordings were obtained from LHb_5HT2c_ and BNST_5HT2c_ neurons using a 5HT_2c_ x Ai9 reporter mouse, which expresses tdTomato in 5HT_2c_-containing neurons. Recordings from both regions were obtained in the same animals 7-10 days following their last DiD session. Mice were rapidly decapitated under isoflurane anesthesia and brains were quickly removed and immersed in a chilled carbogen (95% O_2_/5% CO_2_)-saturated sucrose artificial cerebrospinal fluid (aCSF) cutting solution: 194 mM sucrose, 20 mM NaCl, 4.4 mM KCl, 2 mM CaCl_2_, 1 mM MgCl_2_, 1.2 mM NaH_2_PO_4_, 10 mM D-glucose, and 26 mM NaHCO_3_. Coronal slices containing the LHb or the BNST were prepared on a vibratome at 300 µm and slices were transferred to a holding chamber containing a heated oxygenated aCSF solution: 124 mM NaCl, 4.4 mM KCl, 1 mM NaH_2_PO_4_, 1.2 MgSO_4_, 10 mM D-glucose, 2 mM CaCl_2_, and 26 mM NaHCO_3_. After equilibration for >30 mins, slices were placed in a submerged recording chamber superfused with 30-35 °C oxygenated aCSF (2 mL/min). Cells were visualized under a 40x water immersion objective with video-enhanced differential interference contrast, and a 555 LED was used to visualize fluorescently labeled 5HT_2c_ neurons. Signals were acquired using an Axon Multiclamp 700B amplifier (Molecular Devices) digitized at 10 kHz and filtered at 3 kHz, and subsequently analyzed in pClamp 10.7 or Easy Electrophysiology. Series resistance (R_a_) was monitored and cells were excluded from analysis when changes in R_a_ exceeded 20%. Cells were also excluded from analysis if the current to hold the V_m_ of the cell at 0 mV in current clamp mode exceeded −100 pA.

Potentials were recorded in current-clamp mode with a potassium gluconate-based intracellular solution: 135 mM K-gluconate, 5 mM NaCl, 2 mM MgCl_2_. 10 mM HEPES, 0.6 mM EGTA, 4 mM Na_2_ATP, 0.4 mM Na_2_GTP, pH 7.3, 289-292 Osm. To hold cells at a similar membrane potential for excitability experiments, V_m_ was adjusted to −70 mV by constant current injection. Current injection-evoked action potentials were evaluated by measuring rheobase (minimum current required to evoke an action potential) and number of action potentials fired at linearly increasing current steps (25 pA increments, −100 to 375 pA).

#### Immunohistochemistry

To prepare tissue for immunohistochemistry, mice were anesthetized with an overdose of Avertin (1 ml, i.p.) and transcardially perfused with chilled 0.01 M phosphate-buffered saline (PBS) followed by 4% paraformaldehyde (PFA) in PBS. Brains were extracted and post-fixed in 4% PFA for 24hs and then stored in PBS at 4°C for long-term storage. 45 µm coronal sections were collected using a Leica VT1000S vibratome (Leica Microsystems) and stored in 0.02% Sodium Azide (Sigma Aldrich) in PBS until immunohistochemistry was performed.

To perform immunohistochemistry, tissue was washed for 3×10 minutes in PBS, permeabilized for 30 minutes in 0.5% Triton-X-100 in PBS, and immersed in blocking solution for one hour (0.1% Triton-X-100 + 10% Normal Donkey Serum in PBS). Next, the tissue was incubated overnight at 4°C in primary antibody diluted in blocking solution (anti-RFP 1:500; anti-5-HT 1:1000; anti-cFos 1:1000). The next day, tissue was washed 4×10 minutes in PBS before being incubated for two hours in secondary antibody diluted in PBS (all at 1:200). The tissue was then washed 3×10 minutes in PBS, mounted on slides, and allowed to dry overnight before cover slipping with Vecta-Shield Hardset Mounting Medium with DAPI (Vector Laboratories).

### In-situ Hybridization

To prepare tissue for *in-situ* hybridization (ISH), mice were anesthetized with isoflurane and rapidly decapitated. Brains were extracted and flash frozen in isopentene (Sigma Aldrich) before being stored at −80°C until sectioning. 18 µm coronal sections were collected with a Leica CM3050 S cryostat (Leica Microsystems) and mounted directly on slides. ISH was performed to fluorescently label mRNA for mouse serotonin receptor _2c_ (Mm-Htr_2c_, probe#: 401001), vesicular GABA transporter (Mm-Slc32a1, probe#:319191), and vesicular glutamate transporter 2 (Mm-Slc17a6, probe#: 319171) using the RNAscope Fluorescence Multiplex Assay Kit (Advanced Cell Diagnostics) according to the manufacturer’s instructions. Following ISH, the slides were cover slipped with Prolong-Diamond Mounting Medium with DAPI (Thermo Fisher Scientific). For analysis, a minimum of 5 puncta per cell was used as the criteria for a positive cell for any one mRNA marker. 1-4 images were analyzed per animal per experiment.

### Confocal Microscopy

All fluorescent images were acquired with a Zeiss 800 upright confocal microscope using Zen Blue software (Carl Zeiss). Validation images of viral expression and optic fiber placement were acquired with a 10x objective, while all other immunohistochemistry and ISH images were acquired using a 20x objective. Images were processed and quantified in FIJI (Schindelin et al., 2012).

### Real-Time Quantitative PCR (qPCR)

To prepare tissue for qPCR, mice were anesthetized with isoflurane and rapidly decapitated. Brains were extracted and immediately placed on ice in PBS buffer, then placed in a brain cutting block and sectioned into 1 mm thick slices. Brain punches containing the LHb (one punch) or the BNST (bilateral punches) were collected from 1mm sections using a 1mm diameter tissue micro-punch and immediately flash frozen in a tube block chilled with dry ice. Tissue punches were kept in a −80 °C freezer until processing. RNA was extracted from brain tissue using the RNAeasy Kit (Qiagen) per the manufacturer’s instructions and eluted in water. Following extraction, RNA concentration and quality was assessed using a NanoDrop Spectrophotometer (Thermo Fisher). RNA concentrations were then normalized to 1 ng/µL. Reverse transcription of cDNA from total RNA was performed using SuperScript IV VILO Master Mix with ezDNase according to the manufacturer’s instructions (Thermo Fisher). For each qPCR reaction, 1 ng of cDNA was combined with 10 µl of Taqman Advanced Master Mix (Thermo Fisher), 0.5 µl each of Taqman mouse *Htr2c* (cat. 401001) and *GAPDH* primers (cat. 4331182), and topped off with water to total 20 µl for each reaction. For each mouse, 3 biological replicates were included for each brain region. *GAPDH* and *Htr2c* expression was assessed within single replicates using a multiplex technique with FAM (*Htr2c*) and VIC (*GAPDH*) dyes. Real-time qPCR was performed using the QuantStudio3 System (Thermo Fisher) and analyzed using the 2^-ΔΔCT^ method with *GAPDH* as a housekeeping gene for normalization of *Htr2c* (Livak and Schmittgen, 2001).

### Statistics

Single-variable comparisons between two groups were made using paired or unpaired two-tailed t-tests. Group comparisons were made using one-way ANOVA, two-way ANOVA, or two-way mixed-model ANOVA (depending on the number of independent and within-subject variables). Following significant interactions or main effects, post-hoc pairwise t-tests were performed using Holm-Sidak’s test to control for multiple comparisons. All data are expressed as mean ± standard error of the mean (SEM), with significance defined as p<0.05. All data were analyzed with GraphPad Prism 9 (GraphPad Software).

### Excluded Data

Data points were only excluded from these analyses for the following reasons: missed targeting of viral injections or optic fiber placements, clogged or spilled water/sucrose/alcohol bottles, malfunctions in fiber photometry hardware affecting quality of data collected, data points that were statistically significant outliers (as determined using Grubb’s test, used only once per dataset to identify single outliers), or cells in electrophysiology experiments that did not meet inclusion criteria for health (see electrophysiology section above for these criteria).

## Data Availability

All data in this manuscript are freely available from the corresponding author Dr. Kash via email. This includes raw and processed photometry data, raw behavioral videos and video analysis output, microscopy images and quantification, raw electrophysiology data and analysis, and raw qPCR data and quantification.

## Code Availability

Custom MATLAB scripts developed to process and analyze fiber photometry data are freely available from the corresponding author Dr. Kash via email.

## Extended Data

**Figure S1 (accompanies Figure 3).**
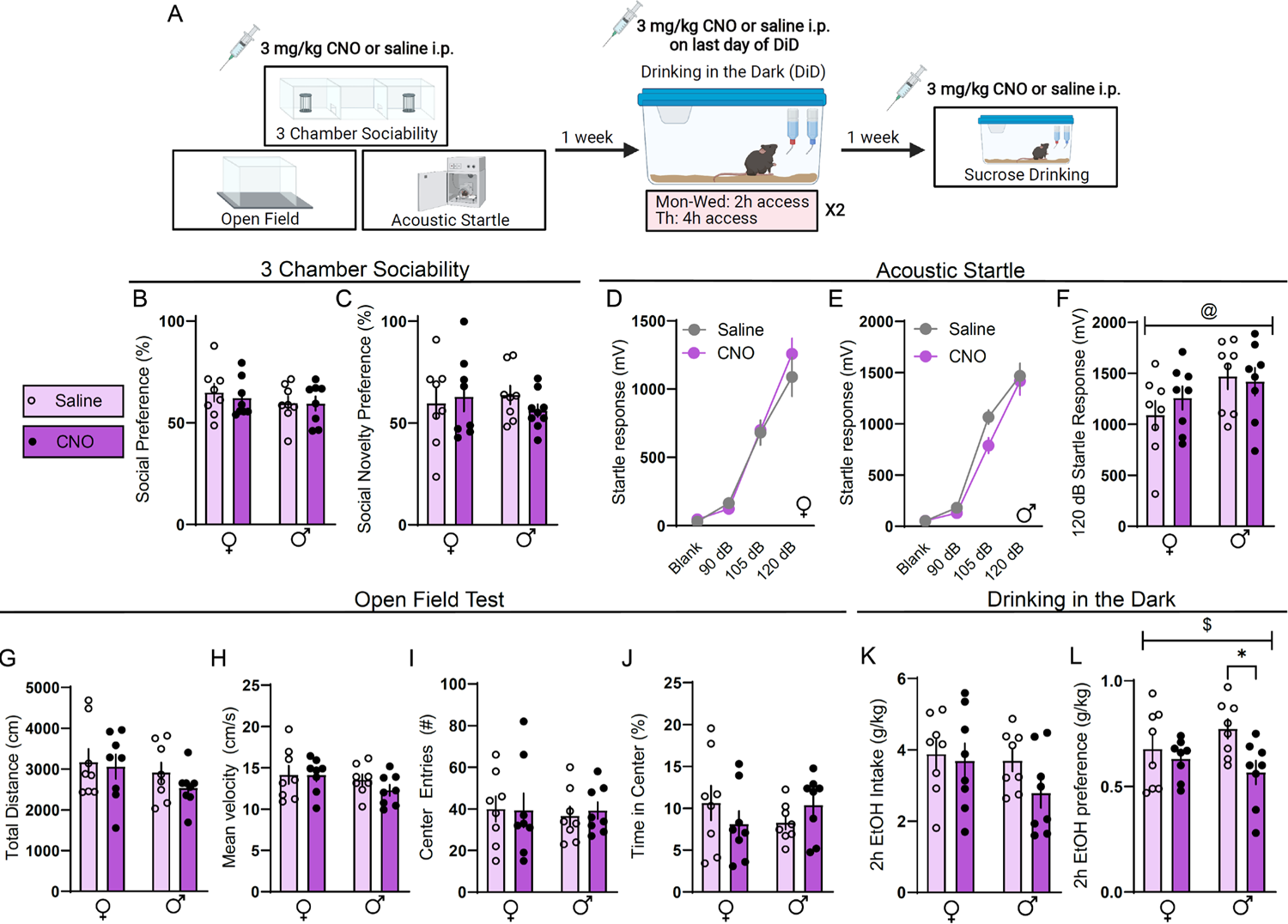
Systemic CNO treatment effects on behavior in wild-type mice. **A,** Experimental timeline for CNO vs. vehicle studies. **B,** Social preference in the 3-chamber sociability test (two-way ANOVA, interaction: F(1,28)=0.1204, p=0.7312, sex: F(1,28)=1.162, p=0.2902, CNO: F(1,28)=0.1526, p=0.6990). **C,** Social novelty preference in 3-chamber sociability test (two-way ANOVA, interaction: F(1,28)=0.6185, p=0.4382, sex: F(1,28)=0.004338, p=0.9480, CNO: F(1,28)=0.05018, p=0.8244). **D,** Acoustic startle behavior, females (two-way repeated measures ANOVA, interaction: F(3,42)=0.9912, p=0.4063, stimulus strength: F(3,42)=135.5, p<0.0001, CNO: F(1,14)=0.3137, p=0.5483). **E,** Acoustic startle behavior, males (two-way repeated measures ANOVA, interaction: F(3,42)=1.663, p=0.1895, stimulus strength: F(3,42)=185.3, p<0.0001, CNO: F(1,14)=2.082, p=0.1711). **F,** Acoustic startle behavior, 120 dB only (two-way ANOVA, interaction: F(1,28)=0.7202, p=0.4033, sex: F(1,28)=4.365, p=0.0459, CNO: F(1,28)=0.2143, p=0.6470). **G,** Total distance in open field (two-way ANOVA, interaction: F(1,28)=0.2579, p=0.6156, sex: F(1,28)=2.160, p=0.1528, CNO: F(1,28)=0.8553, p=0.3629). **H,** Mean velocity in open field (two-way ANOVA, interaction: F(1,28)=0.5936, p=0.4475, sex: F(1,28)=2.523, p=0.1445, CNO: F(1,28)=0.6587, p=0.4239). **I,** Center entries in the open field (two-way ANOVA, interaction: F(1,28)=0.07563, p=0.7852, sex: F(1,28)=0.08771, p=0.7693, CNO: F(1,28)=0.02834, p=0.8668). **J,** Time spent in center of open field (two-way ANOVA, interaction: F(1,28)=2.374, p=0.1346, sex: F(1,28)=0.000948, p=0.0.9957, CNO: F(1,28)=0.01903, p=0.8668). **K,** Alcohol intake in DiD (two-way ANOVA, interaction: F(1,28)=0.7706, p=0.3875, sex: F(1,28)=1.775, p=0.1936, CNO: F(1,28)=1.824, p=0.1877). **L,** Alcohol preference in DiD (two-way ANOVA, interaction: F(1,28)=2.293, p=0.1331, sex: F(1,28)=0.08412, p=0.7739, CNO: F(1,28)=5.595, p=0.0212; Holm-Sidak post-hoc male CNO vs. male saline p=0.0174). **M,** Sucrose consumption (two-way ANOVA, interaction: F(1,28)=1.668, p=0.2071, sex: F(1,28)=1.079, p=0.3079, CNO: F(1,28)=0.001453, p=0.9699). For all panels, n=8 mice/group. All data are presented as mean ± SEM. $ denotes effect of DiD, @ denotes effect of sex, * denotes post-hoc effects.

**Figure S2 (accompanies Figure 3).**
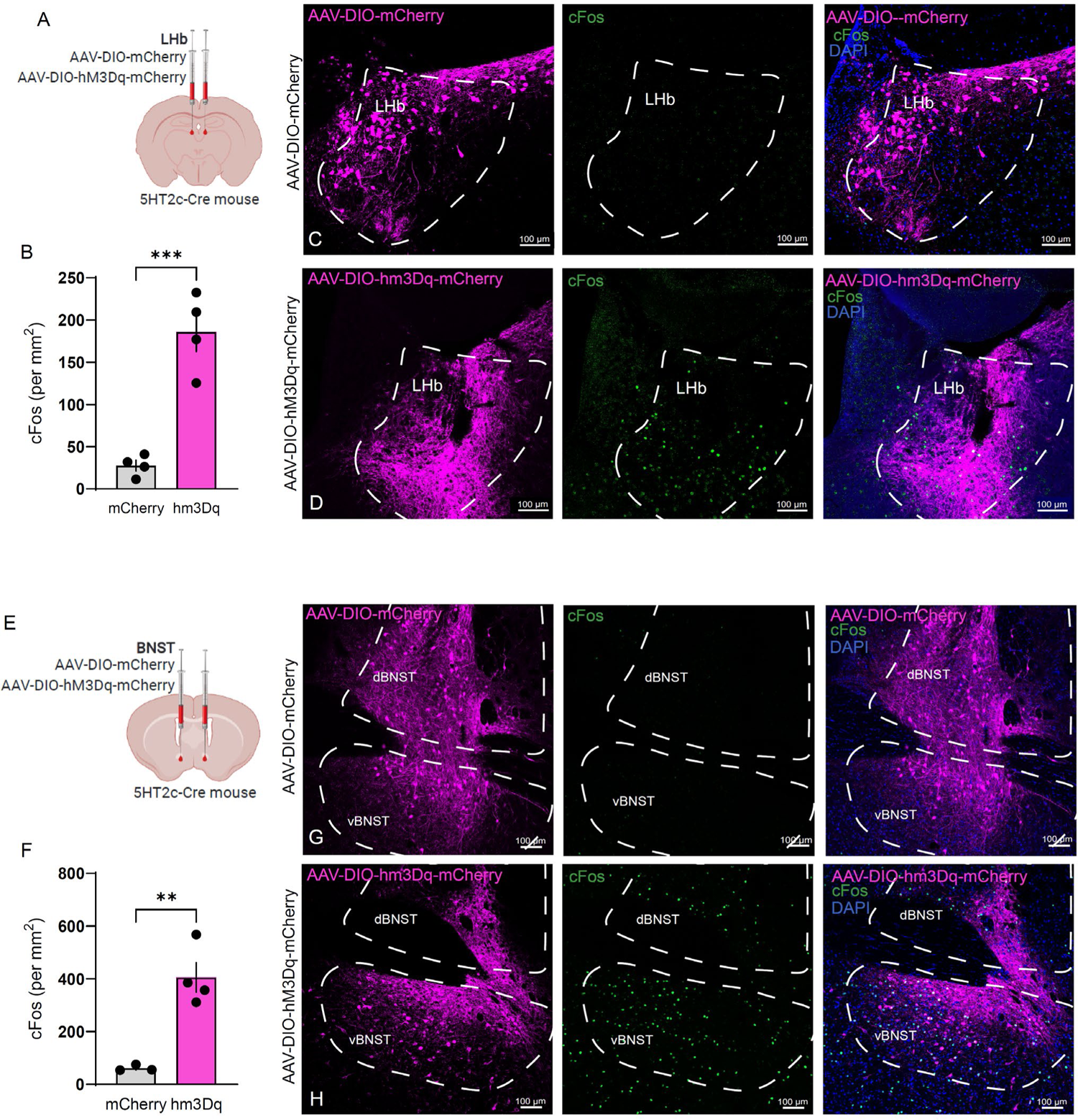
*In-vivo* validation of chemogenetic activation approach. **A,** Surgical schematic for chemogenetic activation of LHb_5HT2c_ neurons. **B,** Quantification of c-Fos following systemic CNO administration (unpaired t-test, t(6)=6.532, p=0.0006). **C,** Representative viral infection (left), c-Fos (middle), and composite (right) for AAV-DIO-mCherry LHb condition. **D,** Representative viral infection (left), c-Fos (middle), and composite (right) for AAV-DIO-hm3Dq-mCherry LHb condition. **E,** Surgical schematic for chemogenetic activation of BNST_5HT2c_ neurons. **F,** Quantificaiton of c-Fos following systemic CNO administration (unpaired t-test, t(5)=5.154, p=0.0036). **G,** Representative viral infection (left), c-Fos (middle), and composite (right) for AAV-DIO-mCherry BNST condition. **H,** Representative viral infection (left), c-Fos (middle), and composite (right) for AAV-DIO-hm3Dq-mCherry BNST condition. For all panels, n=3-4 mice/group, 2-3 slices/mouse. All data presented as mean ± SEM. **p>0.01, ***p<0.001.

**Figure S3 (accompanies Figure 3).**
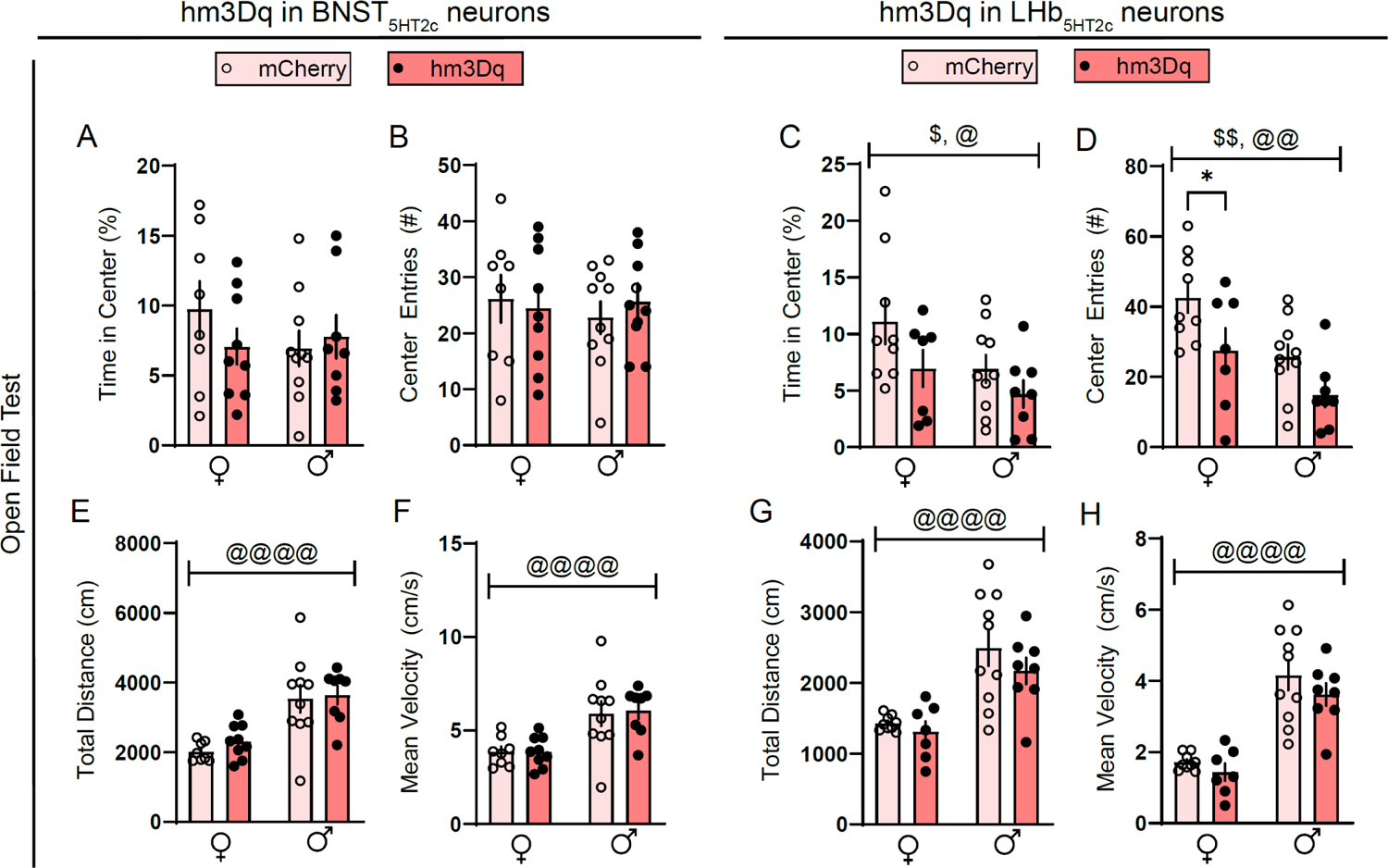
Effects of chemogenetic activation of LHb_5HT2c_ or BNST_5HT2c_ neurons on open field behavior. **A,** Time in center of open field for BNST_5HT2c_ activation (two-way ANOVA, interaction F(1,31)=1.351, p=0.2541, sex: F(1,31)=0.4748, p=0.4959, virus: F(1,31)=0.3689, p=0.584). **B,** Center entries in the open field BNST_5HT2c_ activation (two-way ANOVA, interaction: F(1,31)=0.4112, p=0.5261, sex: F(1,31)=0.09316, p=0.7622, virus: F(1,31)=0.02653, p=0.8717). **C,** Time in center of open field for LHb_5HT2c_ activation (two-way ANOVA, interaction: F(1,30)=0.3754, p=0.5447, sex: F(1,30)=4.229, p=0.0485, virus: F(1,30)=4.191, p=0.0495). **D,** Entries in center of open field for LHb_5HT2c_ activation (two-way ANOVA, interaction: F(1,30)=0.1305, p=0.7204, sex: F(1,30)=10.59, p=0.0029, virus: F(1,30)=9.326, p=0.0047; Holm-Sidak post-hoc female mCherry vs. hM3dq p=0.0489). **E,** Total distance in open field for BNST_5HT2c_ activation (two-way ANOVA, interaction: F(1,31)=0.1347, p=0.7161, sex: F(1,31)=26.89, p<0.0001, virus: F(1,31)=0.5604, p=0.4589). **F,** Mean velocity in open field for BNST_5HT2c_ activation (two-way ANOVA, interaction: F(1,31)=0.0341, p=0.8546, sex: F(1,31)=20.41, p<0.0001, virus: F(1,31)=0.0352, p=0.8523). **G,** Total distance in open field for LHb_5HT2c_ activation (two-way ANOVA, interaction: F(1,30)=0.3027, p=0.5863, sex: F(1,30)=26.49, p<0.0001, virus: F(1,30)=1.413, p=0.2439). **H,** Mean velocity in open field for LHb5HT2c activation (two-way ANOVA, interaction: F(1,30)=0.1694, p=0.6835, sex: 54.73, p<0.0001, virus: F(1,30)=1.745, p=0.1965). For all panels, n=8-10 mice/group. All data presented as mean ± SEM. $ denotes effect of virus, @ denotes effect of sex, and *denote post-hoc effects.

**Figure S4 (accompanies Figure 4).**
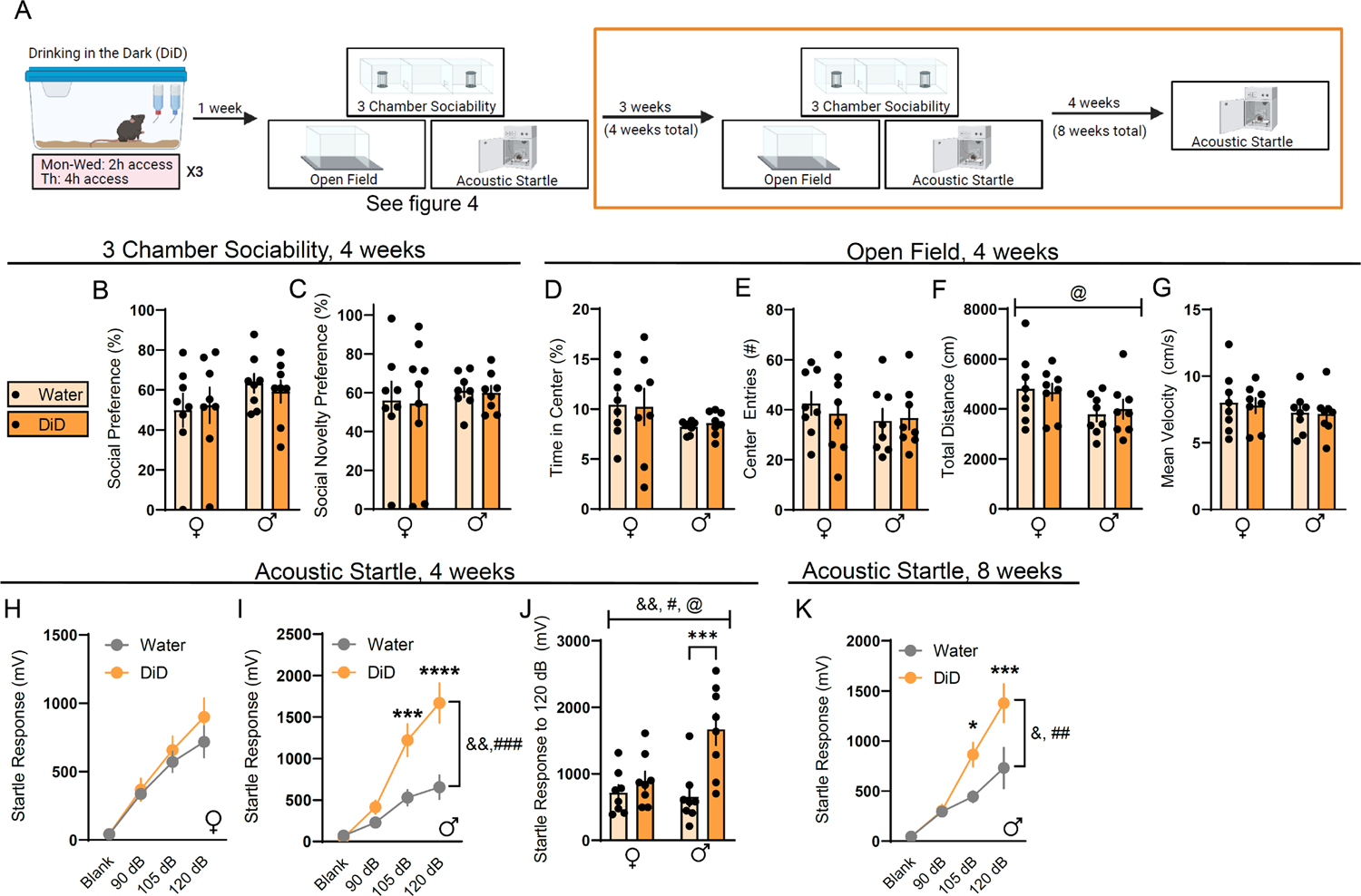
DiD induces long-lasting changes in male acoustic startle behavior. **A,** Experimental timeline for investigating effects of DiD on affective behaviors. This figure is specifically showing behaviors at the time points in the orange box. **B,** Social preference in 3-chamber sociability test (two-way ANOVA, interaction: F(1,27)=0.2612, p=0.6455, sex: F(1,27)=2.057, pp=0.1625, DiD: F(1,27)=0.01356, p=0.9081). **C,** Social novelty preference in the 3-chamber sociability test (two-way ANOVA, interaction: F(1,27)=0.01294, p=0.9176, sex: F(1,27)=0.4284, p=0.5180, DiD: F(1,27)=0.03006, p=0.8636). **D,** Time in center of open field (two-way ANOVA, interaction, F(1,28)=0.0732, p=0.7887, sex: F(1,28)=3.010, p=0.0937, DiD: F(1,28)=0.00598, p=0.9389). **E,** Center entries in open field (two-way ANOVA, interaction: F(1,28)=0.2769, p=0.6029, sex: F(1,28)=0.7692, p=0.3879, DiD: F(1,28)=0.09042, p=0.7659. **F,** Total distance in open field (two-way ANOVA, interaction: F(1,28)=0.2003, p=0.6579, sex: F(1,28)=5.090, p=0.0321, DiD: F(1,28)=0.011, p=0.9172). **G,** Mean velocity in an open field (two-way ANOVA, interaction: F(1,28)=0.01128, p=0.9162, sex: F(1,28)=1.192, p=0.2842, DiD: F(1,28)=0.05686, p=0.8133). **H,** Acoustic startle behavior at four weeks, females (two-way repeated measures ANOVA, interaction: F(3,42)=0.8026, p=0.4995, stimulus strength F(3,42)=55.14, p<0.0001, DiD: F(1,14)=0.7259, p=0.4085). **I,** Acoustic startle behavior at 4 weeks, males (two-way repeated measures ANOVA, interaction: F(3,42)=10.69, p<0.0001, stimulus strength: F(3,42)=48.45, p<0.0001, DiD: F(1,14)=12.83, p=0.0016; Holm-Sidak post-hoc water vs DiD 105 db: p=0.001, 120 dB: p<0.0001. **J,** Acoustic startle behavior at four weeks, 120 dB only (two-way ANOVA, interaction: F(1,28)=6.366, p=0.0176, sex: F(1,28)=4.611, p=0.0406, DiD: F(1,28)=13.16, p=0.0011; Holm-Sidak post-hoc male water vs. male DiD p=0.003). **K,** Acoustic startle behavior at 8 weeks, males (two-way repeated measures ANOVA, interaction: F(3,42)=5.274, p=0.0035, stimulus strength: F(3,42)=39.78, p<0.0001, DiD: F(1,14)=2.207, p=0.0242; Holm-Sidak post-hoc water vs. DiD 105 dB: p=0.0311, 120 dB: p=0.0006). For all panels, n=7-8 mice/group. All data are represented as mean ± SEM. & denotes effects of DiD, @ denotes effect of sex, # denotes interaction, *denote post-hoc effects.

**Figure S5 (accompanies Figure 6).**
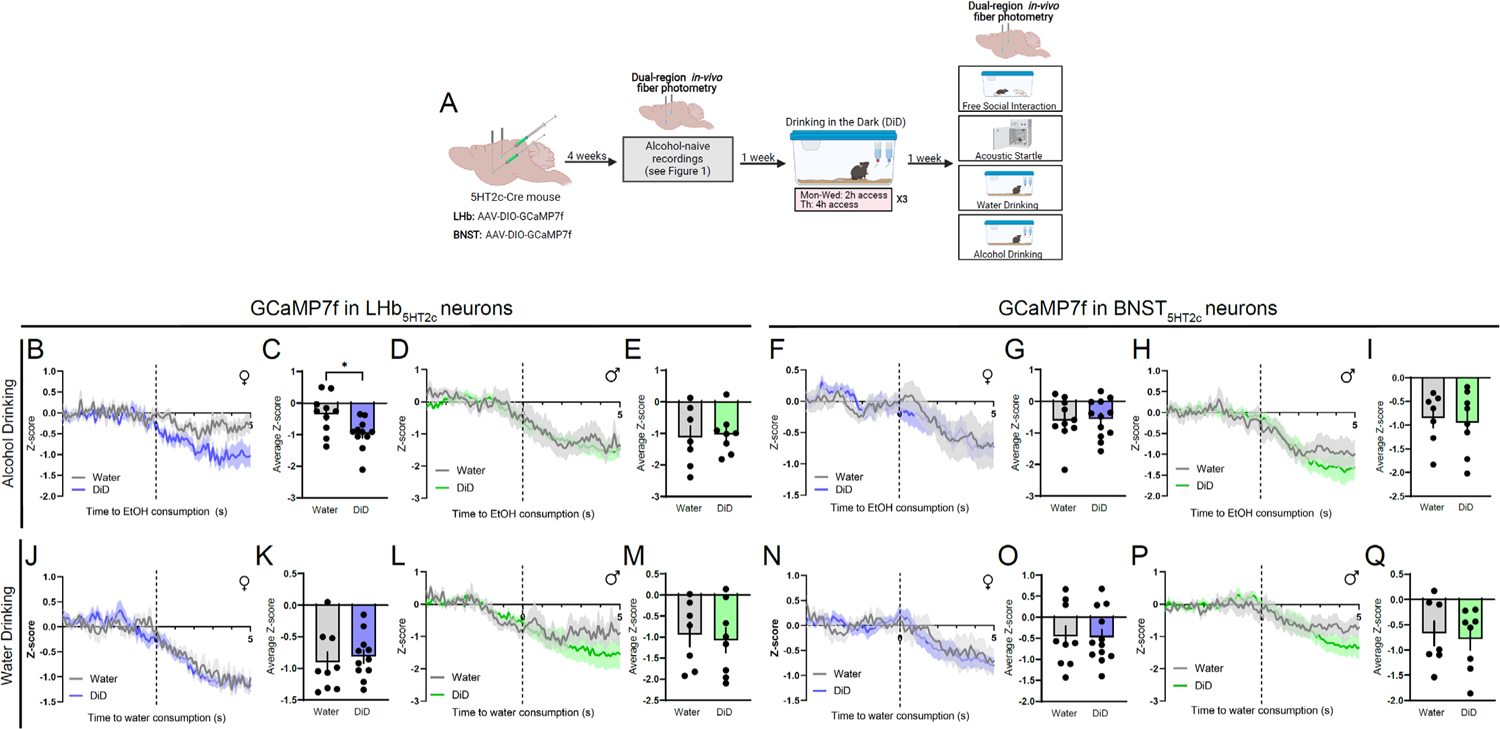
DiD alters calcium (GCaMP) responses of LHb_5HT2c_ neurons in females. **A,** Surgical schematic and experimental timeline for LHb_5HT2c_ and BNST_5HT2c_ GCaMP recordings. **B,** Peri-event plot of LHb_5HT2c_ GCaMP activity during alcohol drinking, females (average of 1-3 bouts/mouse). **C,** Average z-score of LHb_5HT2c_ GCaMP activity for 0-5s post bout start, females (unpaired t-test, t(19)=2.582, p=0.0183). **D,** Peri-event plot of LHb_5HT2c_ GCaMP activity during alcohol drinking, males (average of 1-3 bouts/mouse). **E,** Average z-score of LHb_5HT2c_ GCaMP signal for 0-5s post bout start, males (unpaired t-test, t(13)=0.2215, p=0.8282). **F,** Peri-event plot of BNST_5HT2c_ GCaMP activity during alcohol drinking, females(average of 1-3 bouts/mouse). **G,** Average z-score of BNST_5HT2c_ GCaMP signal for 0-5s post bout start, females (unpaired t-test, t(20)=0.1574, p=0.8765). **H,** Peri-event plot of BNST_5HT2c_ GCaMP signal during alcohol drinking, males (average of 1-3 bouts/mouse). **I,** Average z-score of BNST_5HT2c_ GCaMP signal for 0-5s post bout start, males (unpaired t-test, t(13)=0.2981, p=0.7704). **J,** Peri-event plot of LHb_5HT2c_ GCaMP activity during water drinking, females (average of 1-3 bouts/mouse). **K,** Average z-score of LHb_5HT2c_ signal for 0-5s post bout start, females (unpaired t-test, t(18)=0.4728, p=0.642). **L,** Peri-event plot of LHb_5HT2c_ GCaMP activity during water drinking, males (average of 1-3 bouts/mouse). **M,** Average z-score of LHb_5HT2c_ GCaMP signal for 0-5s post bout start, males (unpaired t-test, t(13)=0.3188, p=0.7549). **N,** Peri-event plot of BNST_5HT2c_ GCaMP activity during water drinking, females (average of 1-3 bouts/mouse). **O,** Average z-score of BNST_5HT2c_ GCaMP signal for 0-5s post bout start (unpaired t-test, t(19)=0.0633, p=0.9502). **P,** Peri-event plot of BNST_5HT2c_ GCaMP activity during water drinking, males (average of 1-3 bouts/mouse). **Q,** Average z-score of BNST_5HT2c_ GCaMP signal for 0-5s post bout start (unpaired t-test, t(13)=0.347, p=0.7342). For all panels, n=7-10 mice, 1-3 bouts/mouse. All data are presented as mean ± SEM. *p<0.05.

**Figure S6 (accompanies Figure 6).**
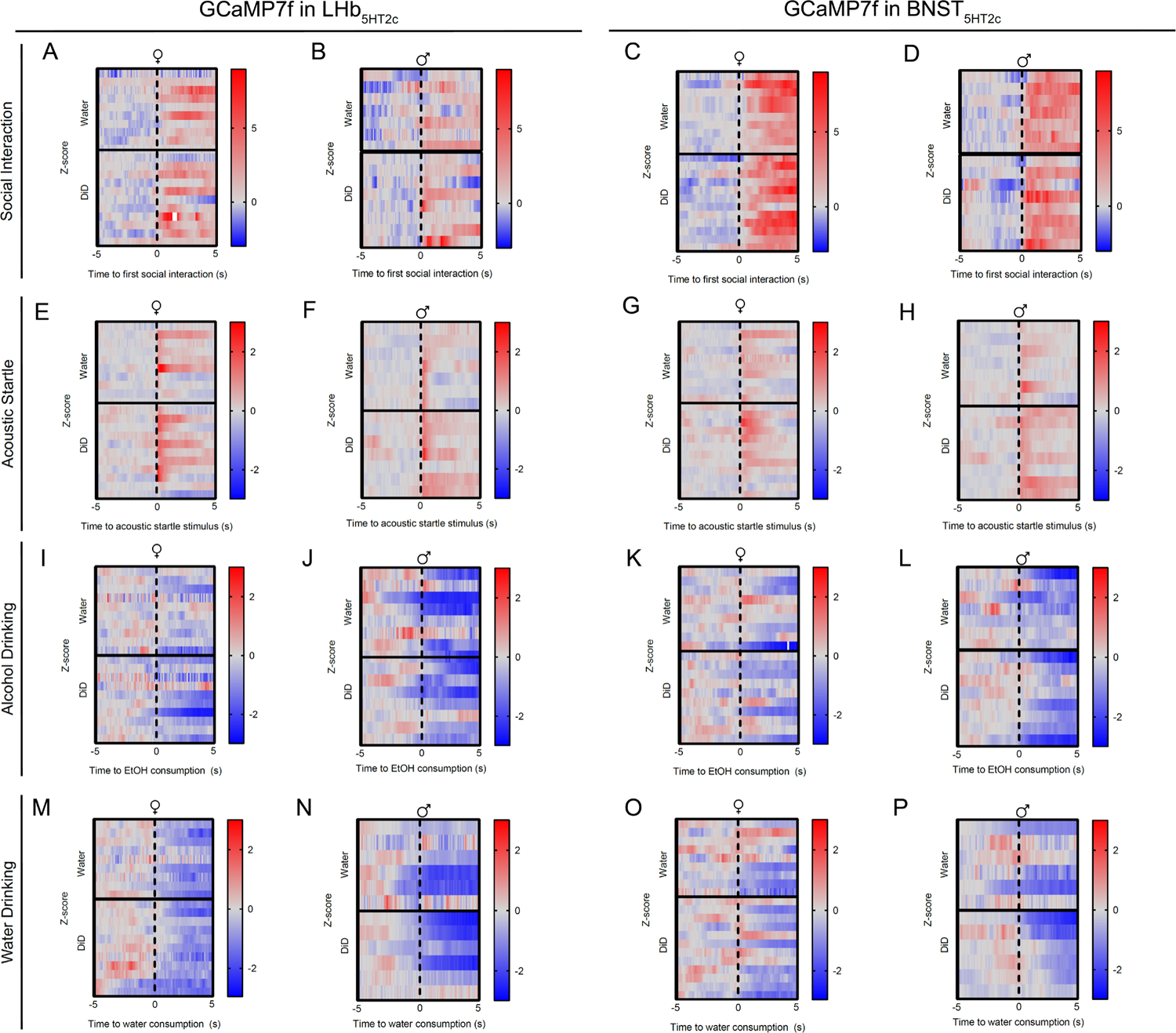
Individual animal responses for GCaMP7f experiments. **A,** Free social interaction, LHb_5HT2c_ females. **B,** Free social interaction, LHb_5HT2c_ males. **C,** Free social interaction, BNST_5HT2c_ females. **D,** Free social interaction, BNST_5HT2c_ males. **E,** Acoustic startle, LHb_5HT2c_ females. **F,** Acoustic startle, LHb_5HT2c_ males. **G,** Acoustic startle, BNST_5HT2c_ females. **H,** Acoustic startle, BNST_5HT2c_ males. **I,** Alcohol drinking, LHb_5HT2c_ females. **J,** Alcohol drinking, LHb_5HT2c_ males. **K,** Alcohol drinking, BNST_5HT2c_ females. **L,** Alcohol drinking, BNST_5HT2c_ males. **M,** Water drinking, LHb_5HT2c_ females. **N,** Water drinking, LHb_5HT2c_ males. **O,** Water drinking, BNST_5HT2c_ males. **P,** Water drinking, BNST_5HT2c_ males. Each horizontal line in the heatmap corresponds to the average of all trials for a single animal.

**Figure S7 (accompanies Figure 6).**
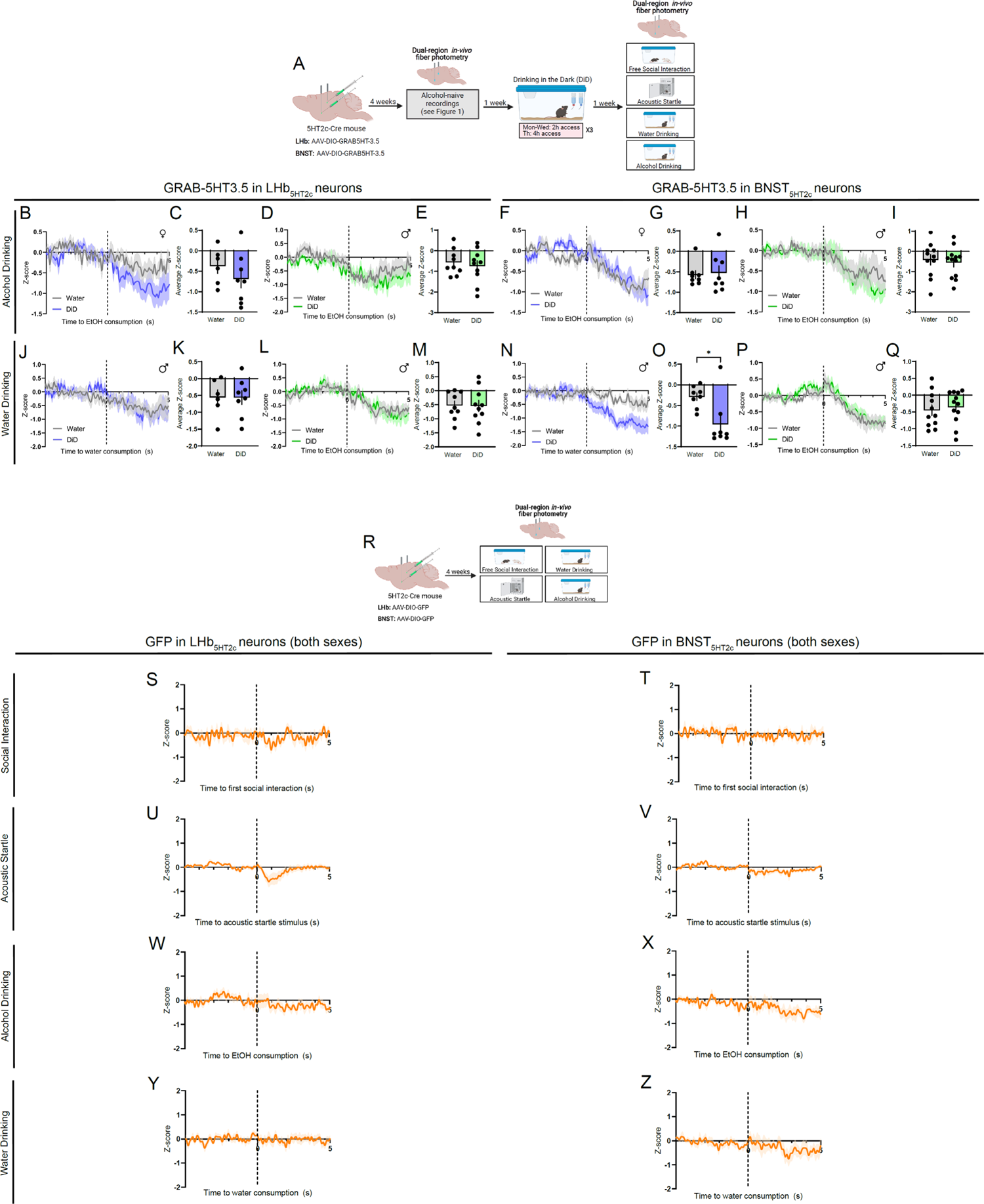
DiD alters 5-HT release onto BNST_5HT2c_ neurons in females during water drinking; GFP motion controls. **A,** Surgical schematic and experimental timeline for LHb_5HT2c_ and BNST_5HT2c_ GRAB-5HT recordings. **B,** Peri-event plot of LHb_5HT2c_ GRAB-5HT activity during alcohol drinking, females (average of 1-3 bouts/mouse). **C,** Average z-score of LHb_5HT2c_ GRAB-5HT activity for 0-5s post bout start, females (unpaired t-test, t(12)=1.058, p=0.3107). **D,** Peri-event plot of LHb_5HT2c_ GRAB-5HT activity during alcohol drinking, males (average of 1-3 bouts/mouse). **E,** Average z-score of LHb_5HT2c_ GRAB-5HT signal for 0-5s post bout start, males (unpaired t-test, t(17)=0.5325, p=0.6013). **F,** Peri-event plot of BNST_5HT2c_ GRAB-5HT activity during alcohol drinking, females (average of 1-3 bouts/mouse). **G,** Average z-score of BNST_5HT2c_ GRAB-5HT signal for 0-5s post bout start, females (unpaired t-test, t(14)=0.2529, p=0.804). **H,** Peri-event plot of BNST_5HT2c_ GRAB-5HT signal during alcohol drinking, males (average of 1-3 bouts/mouse). **I,** Average z-score of BNST_5HT2c_ GRAB-5HT signal for 0-5s post bout start, males (unpaired t-test, t(21)=0.3883, p=0.7017). **J,** Peri-event plot of LHb_5HT2c_ GRAB-5HT activity during water drinking, females (average of 1-3 bouts/mouse). **K,** Average z-score of LHb_5HT2c_ signal for 0-5s post bout start, females (unpaired t-test, t(12)=0.01504, p=0.9882). **L,** Peri-event plot of LHb_5HT2c_ GRAB-5HT activity during water drinking, males (average of 1-3 bouts/mouse). **M,** Average z-score of LHb_5HT2c_ GRAB-5HT signal for 0-5s post bout start, males (unpaired t-test, t(17)=0.08381, p=0.9342). **N,** Peri-event plot of BNST_5HT2c_ GRAB-5HT activity during water drinking, females (average of 1-3 bouts/mouse). **O,** Average z-score of BNST_5HT2c_ GRAB-5HT signal for 0-5s post bout start (unpaired t-test, t(14)=2.917, p=0.0113). **P,** Peri-event plot of BNST_5HT2c_ GRAB-5HT activity during water drinking, males (average of 1-3 bouts/mouse). **Q,** Average z-score of BNST_5HT2c_ GRAB-5HT signal for 0-5s post bout start (unpaired t-test, t(21)=0.4308, p=0.671). For all GRAB-5HT panels, n=7-10 mice. **R,** Surgical schematic and experimental timeline for LHb_5HT2c_ and BNST_5HT2c_ GFP control photometry experiments. **S,** Peri-event plot of LHb_5HT2c_ GFP signal during free social interaction (1 trial/mouse). **T,** Peri-event plot of BNST_5HT2c_ GFP signal during free social interaction (1 trial/mouse). **U,** Peri-event plot of LHb_5HT2c_ GFP signal during the acoustic startle test (average of 10 trials/mouse). **V,** Peri-event plot of BNST_5HT2c_ GFP signal during the acoustic startle test (average of 10 trials/mouse). **W,** Peri-event plot of LHb_5HT2c_ GFP signal during voluntary alcohol drinking (average of 1-3 bouts/mouse). **X,** Peri-event plot of BNST_5HT2c_ GFP signal during voluntary alcohol consumption (average of 1-3 bouts/mouse). **Y,** Peri-event plot of LHb_5HT2c_ GFP signal during voluntary water drinking (average of 1-3 bouts/mouse). **Z,** Peri-event plot of BNST_5HT2c_ GFP signal during voluntary water drinking (average of 1-3 bouts/mouse). For all GFP panels, n=8-9 mice. All data are presented as mean ± SEM. *p<0.05.

**Figure S8 (accompanies Figure 6).**
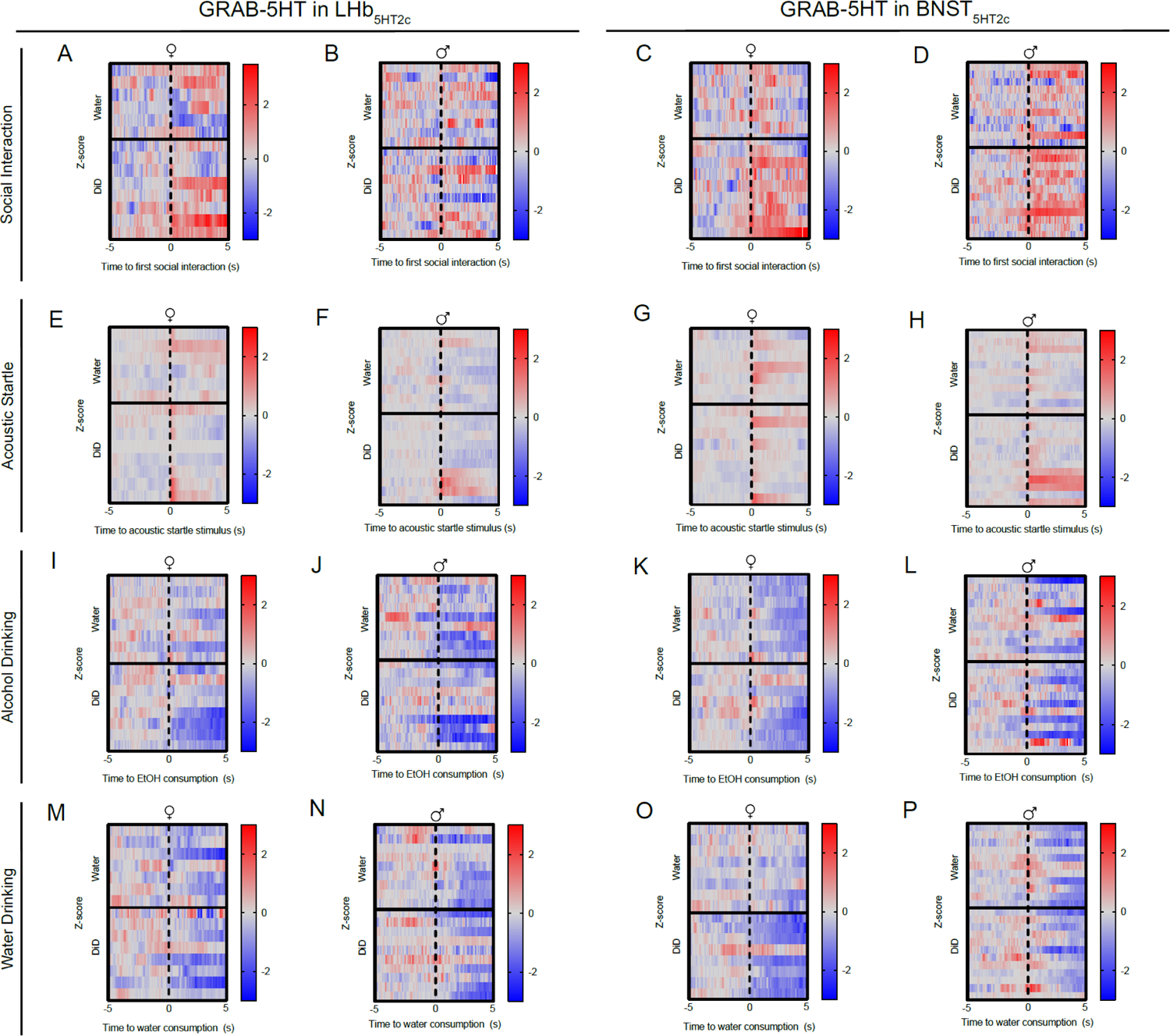
Individual animal responses for GRAB-5HT experiments. **A,** Free social interaction, LHb_5HT2c_ females. **B,** Free social interaction, LHb_5HT2c_ males. **C,** Free social interaction, BNST_5HT2c_ females. **D,** Free social interaction, BNST_5HT2c_ males. **E,** Acoustic startle, LHb_5HT2c_ females. **F,** Acoustic startle, LHb_5HT2c_ males. **G,** Acoustic startle, BNST_5HT2c_ females. **H,** Acoustic startle, BNST_5HT2c_ males. **I,** Alcohol drinking, LHb_5HT2c_ females. **J,** Alcohol drinking, LHb_5HT2c_ males. **K,** Alcohol drinking, BNST_5HT2c_ females. **L,** Alcohol drinking, BNST_5HT2c_ males. **M,** Water drinking, LHb_5HT2c_ females. **N,** Water drinking, LHb_5HT2c_ males. **O,** Water drinking, BNST_5HT2c_ males. **P,** Water drinking, BNST_5HT2c_ males. Each horizontal line in the heatmap corresponds to the average of all trials for a single animal.

**Figure S9 (accompanies Figure 7).**
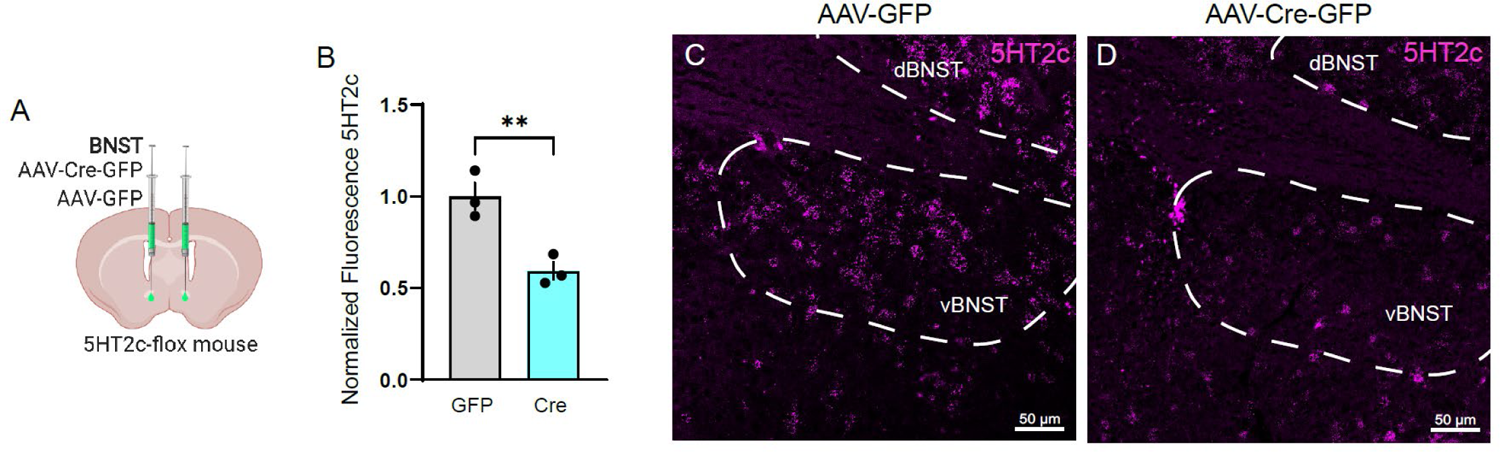
Validation of 5HT_2c_ knockdown in our 5HT_2c_*^lox/lox^* mouse. **A,** Surgical schematic for 5HT_2c_ knockdown. **B,** Quantification of 5HT2c fluorescence (unpaired t-test, n=3/group, t(4)=4.641, p=0.0097). **C,** Representative 5HT2c fluorescence in AAV-GFP condition. **D,** Representative 5HT2c fluorescence in AAV-Cre-GFP condition. All data presented as mean ± SEM. **p<0.01.

**Figure S10 (accompanies Figure 7).**
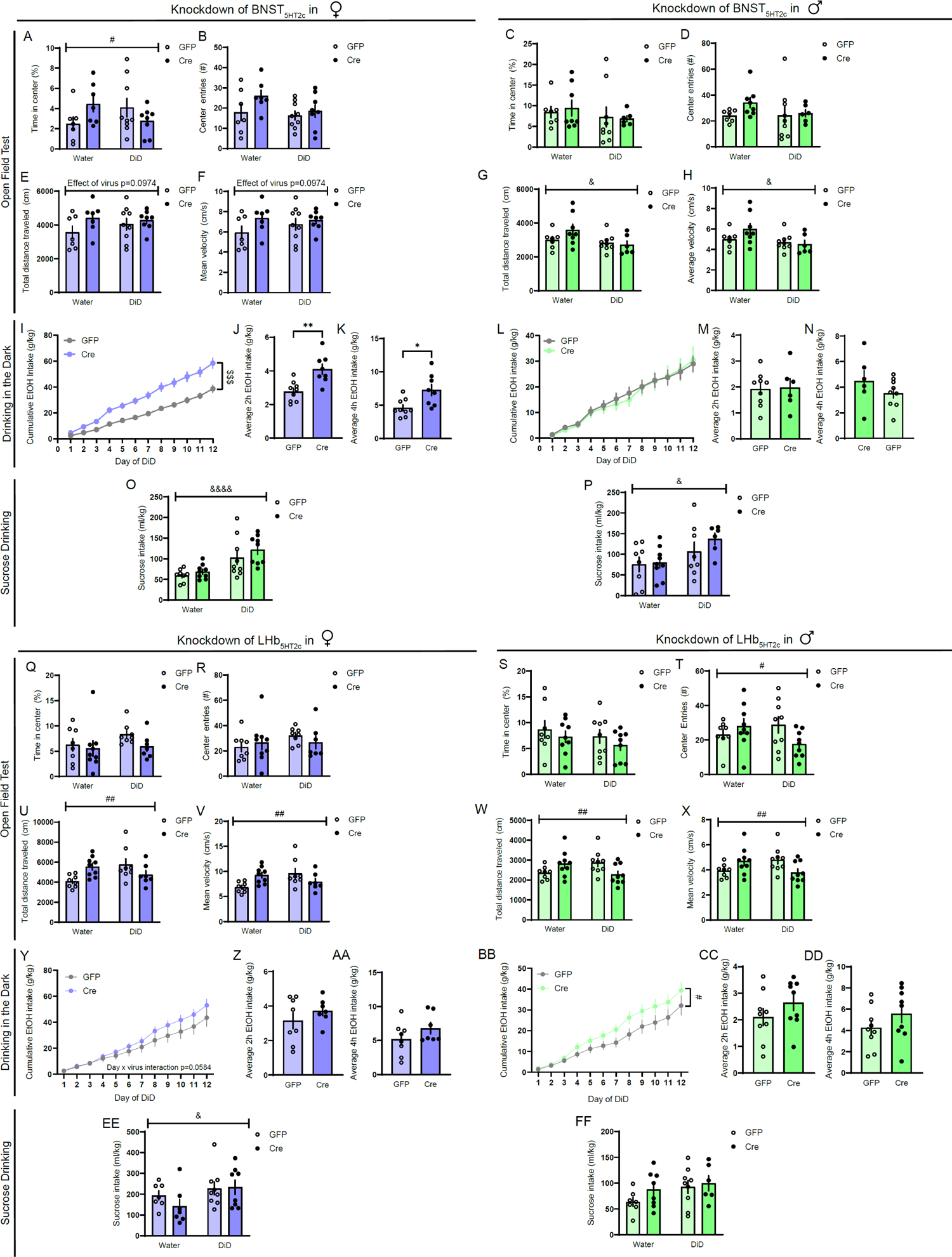
Effects of 5HT_2c_ knockdown on open field behavior, DiD, and sucrose consumption. **A,** Time in center of open field, BNST females (two-way ANOVA, interaction: F(1,27)=4.799, p=0.0373, DiD: F(1,27)=0.0028, p=0.9581, virus: F(1,27)=0.1789, p=0.6757). **B,** Center entries in open field, BNST females (two-way ANOVA, interaction: F(1,27)=0.9661, p=0.334, DiD: F(1,27)=2.203, p=0.1494, virus: F(1,27)=2.875, p=0.1015). **C,** Time in center of open field, BNST males (two-way ANOVA, interaction: F(1,26)=0.1345, p=0.7167, DiD: F(1,26)=0.9586, p=0.3366, virus: F(1,26)=0.0253, p=0.8746). **D,** Center entries in open field, BNST males (two-way ANOVA, interaction: F(1,26)=0.8505, p=0.3649, DiD: F(1,26)=0.6862, p=0.4150, virus: F(1,26)=1.477, p=0.2351). **E,** Total distance in open field, BNST females (two-way ANOVA, interaction: F(1,27)=0.9632, p=0.3351, DiD: F(1,27)=0.3147, p=0.5794, virus: F(1,27)=2.498, p=0.0974). **F,** Mean velocity in open field, BNST females (two-way ANOVA, interaction: F(1,27)=0.9632, p=0.3351, DiD: F(1,27)=0.3147, p=0.5794, virus: F(1,27)=2.498, p=0.0974). **G,** Total distance in open field, BNST males (two-way ANOVA, interaction: F(1,26)=2.166, p=0.1531, DiD: F(1,26)=4.651, p=0.0405, virus: F(1,26)=0.9236, p=0.3435). **H,** Mean velocity in open field, BNST males (two-way ANOVA, interaction: F(1,26)=2.166, p=0.1531, DiD: F(1,26)=4.651, p=0.0405, virus: F(1,26)=0.9236, p=0.3435). **I,** Cumulative alcohol consumption in DiD, BNST females (two-way repeated measures ANOVA, interaction: F(11,165)=13.93, p<0.0001, day: F(11,165)=342.5, p<0.0001, virus: F(1,15)=16.94, p=0.0009; Holm-Sidak post-hoc GFP vs. Cre all days except day 1 p<0.05). **J,** Average 2h alcohol intake in DiD, BNST females (unpaired t-test, t(15)=3.741, p=0.002). **K,** Average 4h alcohol intake in DiD, BNST females (student’s unpaired t-test, t(15)=2.920, p=0.0106). **L,** Cumulative alcohol consumption in DiD, BNST males (two-way repeated measures ANOVA, interaction: F(11,143)=0.3609, p=0.9688, day: F(11,143)=98.92, p<0.0001, virus: F(1,13)=0.02151, p=0.8857). **M,** Average 2h alcohol intake in DiD, BNST males (unpaired t-test, t(13)=0.1509, p=0.8824). **N,** Average 4h alcohol intake in DiD, BNST males (student’s unpaired t-test, t(13)=1.194, p=0.2537). **O,** Sucrose intake, BNST females (two-way ANOVA, interaction: F(1,27)=0.8971, p=0.3520, DiD: F(1,27)=4.022, p=0.0500, virus: F(1,27)=0.5658, p=0.4584). **P,** Sucrose intake, BNST males (two-way ANOVA, interaction: F(1,26)=0.4579, p=0.5046, DiD: F(1,26)=2.754, p=0.1090, virus: F(1,26)=1.598, p=0.2174). **Q,** Time in center of open field, LHb females (two-way ANOVA, interaction: F(1,28)=0.4915, p=0.4891, DiD: F(1,28)=1.060, p=0.3119, virus: F(1,28)=1.671, p=0.2067). **R,** Center entries in open field, LHb females (two-way ANOVA, interaction: F(1,28)=0.9930, p=0.3275, DiD: F(1,28)=1.066, p=0.3108, virus: F(1,28)=0.04981, 0.8250). **S,** Time in center of open field, LHb males (two-way ANOVA, interaction: F(1,31)=0.01133, p=0.9159, DiD: F(1,31)=1.367, p=0.2512, virus: F(1,31)=1.453, p=0.2372). **T,** Center entries in open field, LHb males (two-way ANOVA, interaction: F(1,31)=4.409, p=0.044, DiD: F(1,31)=0.3780, p=0.5432, virus: F(1,31)=0.6709, p=0.4190). **U,** Total distance in open field, LHb females (two-way ANOVA, interaction: F(1,28)=9.334, p=0.0049, DiD: F(1,28)=1.173, p=0.2879, virus: F(1,28)=0.3086, p=0.5830; Holm-Sidak post-hoc water GFP vs DiD GFP p=0.0386). **V,** Mean velocity in open field, LHb females (two-way ANOVA, interaction: F(1,28)=9.334, p=0.0049, DiD: F(1,28)=1.173, p=0.2879, virus: F(1,28)=0.3086, p=0.5830; Holm-Sidak post-hoc water GFP vs DiD GFP p=0.0386). **W,** Total distance in open field, LHb males (two-way ANOVA, interaction: F(1,31)=8.657, p=0.0064, DiD: F(1,31)=0.003814, p=0.9504, virus: F(1,31)=0.1529, p=0.6978). **X,** Mean velocity in open field, LHb males (two-way ANOVA, interaction: F(1,31)=8.657, p=0.0064, DiD: F(1,31)=0.003814, p=0.9504, virus: F(1,31)=0.1529, p=0.6978). **Y,** Cumulative alcohol consumption in DiD, LHb females (two-way repeated measures ANOVA, interaction: F(11,143)=1.804, p=0.0584, day: F(11,143)=114.7, p<0.0001, virus: F(1,13)=1.026, p=0.3295). **Z,** Average 2h alcohol intake in DiD, LHb females (unpaired t-test, t(13)=1.026, p=0.3237). **AA,** Average 4h alcohol intake in DiD, LHb females (student’s unpaired t-test,t(13)=1.335, p=0.2048). **BB,** Cumulative alcohol consumption in DiD, LHb males (two-way repeated measures ANOVA, interaction: F(11,176)=1.970, p=0.0339, day: F(11,176)=111.9, p<0.0001, virus: F(1,16)=2.319, p=0.1473). **CC,** Average 2h alcohol intake in DiD, males (unpaired t-test, t(13)=0.1509, p=0.8824). **DD,** Average 4h alcohol intake in DiD, LHb males (student’s unpaired t-test, t(16)=1.266, p=0.2235). **EE,** Sucrose intake, females (two-way ANOVA, interaction: F(1,27)=0.8971, p=0.3520, DiD: F(1,27)=4.022, p=0.0500, virus: F(1,27)=0.5658, p=0.4584). **FF,** Sucrose intake, males (two-way ANOVA, interaction: F(1,26)=0.4579, p=0.5046, DiD: F(1,26)=2.754, p=0.1090, virus: F(1,26)=1.598, p=0.2174). For all panels, n=6-10 mice/group. All data presented as mean ± SEM. & denotes effect of DiD, $ denotes effect of virus, # denotes interaction (day x virus or DiD x virus), and * denotes post-hoc comparisons.

**Figure S11 (accompanies Figure 8).**
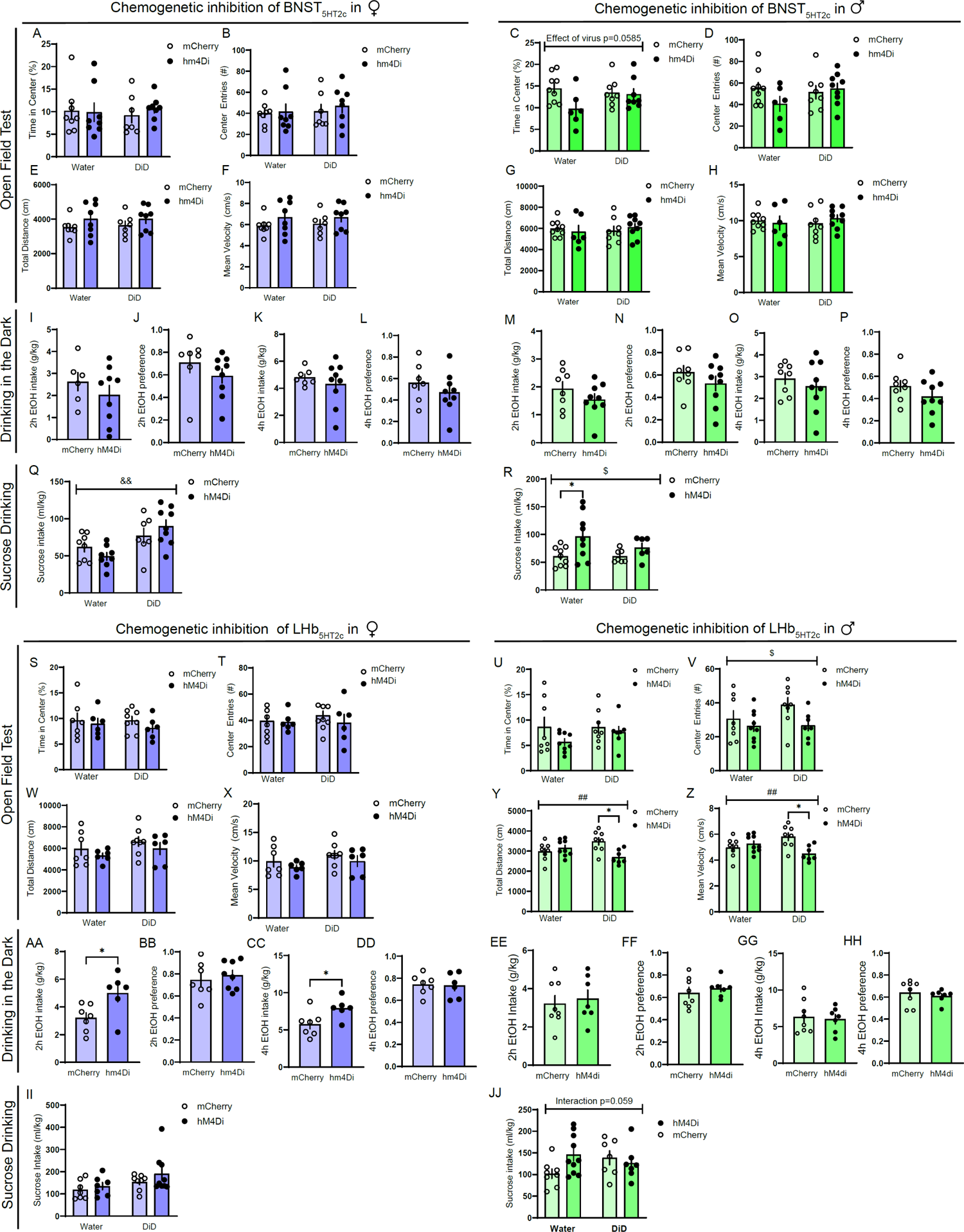
Effects of chemogenetic inhibition of LHb_5HT2c_ and BNST_5H2c_ on open field behavior, DiD, and sucrose consumption. **A,** Time in center of open field (%), BNST females (two-way ANOVA, interaction: F(1,28)=0.2804, p=0.6007, DiD: F(1,28)=0.0029, p=0.9573, F(1,28)=0.1157, p=0.7363). **B,** Center entries in open field, BNST females (two-way ANOVA, interaction: F(1,28)=0.0813, p=0.7776, DiD: F(1,28)=0.3737, p=.5461, virus: F(1,28)=0.3027, p=0.5876). **C,** Time in center of open field (%), BNST males (two-way ANOVA, interaction: F(1,28)=2.801, p=0.1057, DiD: F(1,28)=0.8712, p=0.3589, virus: F(1,28)=3.902, p=0.0585). **D,** Center entries in open field, BNST males (two-way ANOVA, interaction: F(1,28) =0.8397, p=0.3673, virus: F(1,28)=0.9497, p=0.3381). **E,** Total distance traveled in open field, BNST females (two-way ANOVA, interaction: F(1,28)=0.04142, p=0.8403, DiD: F(1,28)=0.04283, p=0.8376, virus: F(1,28)=2.737, p=0.1096). **F,** Mean velocity in open field, BNST females (two-way ANOVA, interaction: F(1,28)=0.0414, p=0.8403, DiD: F(1,27)=0.0428, p=0.1096). **G,** Total distance traveled in open field, BNST males (two-way ANOVA, interaction: F(1,28)=0.6688, p=0.4204, DiD: F(1,28)=0.0761, p=0.7846, virus: F(1,28)=0.0010, p=0.9741). **H,** Mean velocity in open field, BNST males (two-way ANOVA, interaction: F(1,28)=0.7183, p=0.4039, DiD: F(1,28)=0.0539, p=0.8181, virus: F(1,28)=0.0895, p=0.7670). **I,** 2 hr alcohol intake in DiD, BNST females (student’s unpaired t-test, t(14)=1.355, p=0.1969). **J,** 2 hr alcohol preference in DiD, BNST females (student’s unpaired t-test, t(14)=1.114, p=0.2840). **K,** 4h alcohol intake in DiD, BNST females (student’s unpaired t-test, t(14)=0.7428, p=0.4699). **L,** 4h alcohol preference in DiD, BNST females (student’s unpaired t-test, t(14)=0.9671, p=0.3499). **M,** 2h alcohol intake in DiD, BNST males (student’s unpaired t-test, t(15)=1.224, p=0.2398). **N,** 2h alcohol preference in DiD, BNST males (student’s unpaired t-test, t(15)=1.132, p=0.2753). **O,** 4h alcohol intake in DiD, BNST males (student’s unpaired t-test, t(15)=0.7164, p=0.4847). **P,** 4h alcohol preference in DiD, BNST males (student’s unpaired t-test, t(15)=1.318, p=0.2073). **Q,** 4h sucrose intake, BNST females (two-ay ANOVA, interaction: F(1,28)=2.708, p=0.111, DiD: F(1,28)=13.22, p=0.0011, virus: F(1,28)=0.0006, p=0.9801; Holm-Sidak test for multiple comparisons water hm4Di vs. DiD hm4Di p=0.003). **R,** 4h sucrose intake, BNST males (two-way ANOVA, interaction: F(1,28)=1.134, p=0.2964, DiD: F(1,28)=1.119, p=0.2996, virus: F(1,17)=7.093, p=0.0129; Holm-Sidak test for multiple comparisons, water mCherry vs. water hm4Di p=0.0449). **S,** Time in center of open field (%), LHb females (two-way ANOVA, interaction: F(1,23)=0.1562, p=0.6963, DiD: F(1,23)=0.1617, p=0.6913, virus: F(1,23)=1.021, p=0.3229). **T,** Center entries to open field, LHb females (two-way ANOVA, interaction: F(1,23)=0.3261, p=0.5735, DiD: F(1,23)=0.1708, p=0.6832, virus: F(1,23)=0.6076, p=0.4437). **U,** Time in center of open field (%), LHb males (two-way ANOVA, interaction: F(1,28)=-.6047, p=0.4433, DiD: F(1,28)=0.5476, p=0.4655, virus: F(1,28)=2.396, p=0.1329). **V,** Center entries to open field, LHb males (two-way ANOVA, interaction: F(1,28)=1.008, p=0.3239, DiD: F(1,28)=1.216, p=0.2795, virus: F(1,28)=4.482, p=0.0433). **W,** Total distance traveled in open field, LHb females (two-way ANOVA, interaction: F(1,23)=0.00011, p=0.9915, DiD: F(1,23)=1.799, p=0.1930, virus: F(1,23)=1.731, p=0.2013). **X,** Mean velocity in open field, LHb females (two-way ANOVA, interaction: F(1,23)=0.00011, p=0.9915, DiD: F(1,23)=1.799, p=0.1930, virus: F(1,23)=1.731, p=0.2013). **Y,** Total distance traveled in open field, LHb males (two-way ANOVA, interaction: F(1,28)=9.503, p=0.0046, DiD: F(1,28)=0.02741, p=0.8697, virus: F(1,28)=3.813, p=0.0609; Holm-Sidak test for multiple comparisons DiD mCherry vs. DiD hm4Di p=0.0106). **Z,** Mean velocity in open field, LHb males (two-way ANOVA, interaction: F(1,28)=9.505, p=0.0046, DiD: F(1,28)=0.02668, p=0.8714, virus: F(1,28)=3.813, p=0.0613; Holm-Sidak test for multiple comparisons DiD mCherry vs. DiD hm4Di p=0.0106). **AA,** 2h alcohol intake in DiD, LHb females (student’s unpaired t-test, t(11)=2.561, p=0.0265). **BB,** 2h alcohol preference in DiD, LHb females (student’s unpaired t-test, t(11)=0.09569, p=0.9255). **CC,** 4h alcohol intake in DiD, LHb females (student’s unpaired t-test, t(11)=2,488, p=0.0301). **DD,** 4h alcohol preference in DiD, LHb females (student’s unpaired t-test, t(11)=0.1693, p=0.8686). **EE,** 2h alcohol intake in DiD, LHb males (student’s unpaired t-test, t(13)=0.4194, p=0.6817). **FF,** 2h alcohol preference in DiD, LHb males (student’s unpaired t-test, t(13)=0.6729, p=0.5128). **GG,** 4h alcohol intake in DiD, LHb males (student’s unpaired t-test, t(13)=0.3153, p=0.7576). **HH,** 4h alcohol preference in DiD, LHb males (student’s unpaired t-test, t(13)=0.6684, p=0.5156). **II,** 4h sucrose intake, LHb females (two-way ANOVA, interaction: F(1,23)=0.2698, p=0.6084, DiD: F(1,23)=4.057, p=0.0558, virus: F(1,23)=2.422, p=0.1333). **JJ,** 4h sucrose intake, LHb males (two-way ANOVA, interaction: F(1,28)=3.886, p=0.0586, F(1,28)=0.3643, p=0.5510, virus: F(1,28)=1.249, p=-0.2733).

**Figure S12:**
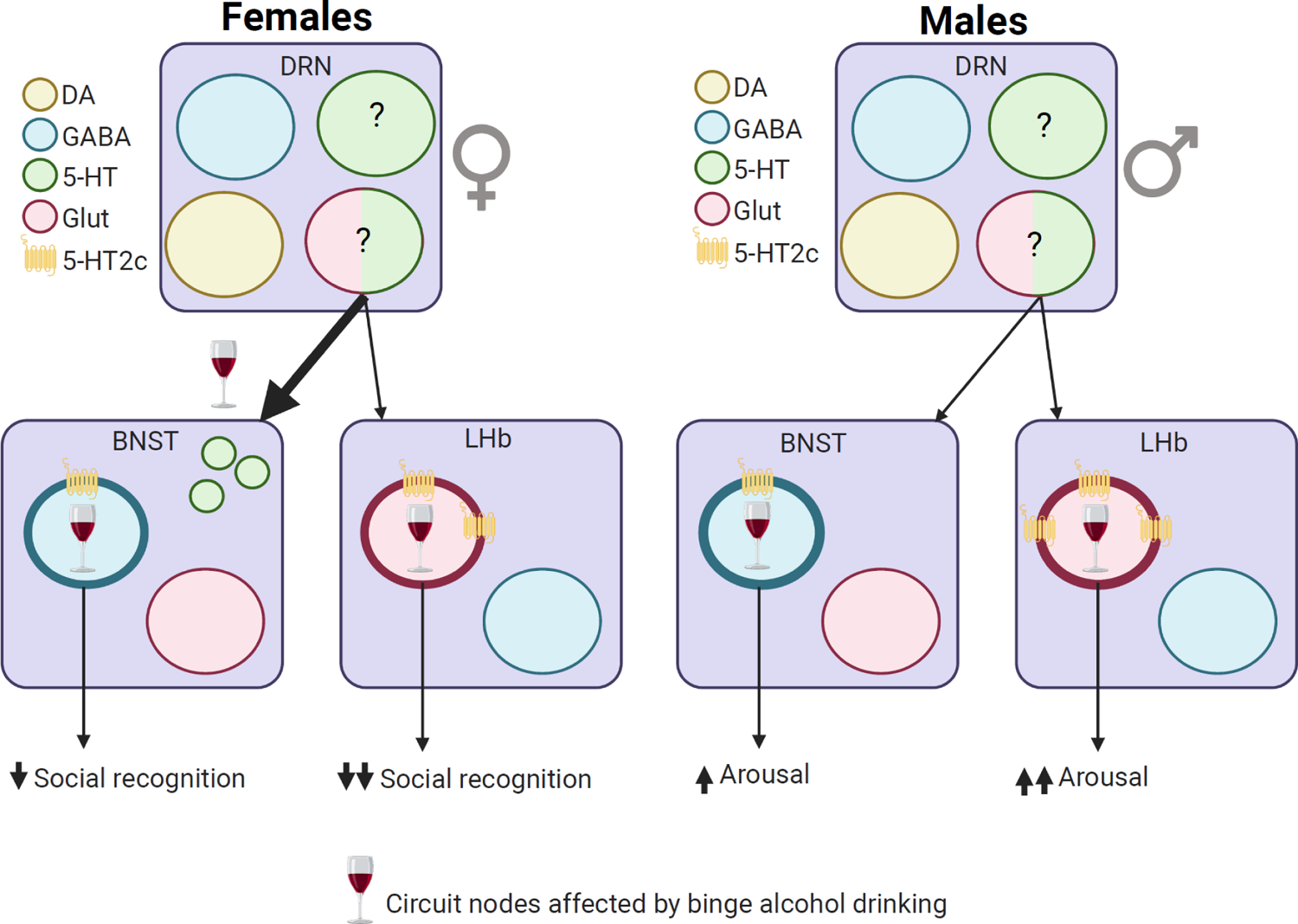
Summary of binge alcohol effects on DRN-BNST and DRN-LHb circuitry. Single 5-HT neurons in the DRN project to both BNST and LHb. In females, binge alcohol consumption increases 5-HT release in the BNST upon social interaction, decreases BNST_5HT2c_ excitability in slice, and increases medial LHb_5HT2c_ excitability in slice. 5HT_2c_ in the BNST and the LHb partially regulate social recognition deficits induced by alcohol, but excessive activity of LHb_5HT2c_ predominantly drives alcohol-induced social recognition deficits in females. In males, binge alcohol consumption increases LHb_5HT2c_ and BNST_5HT2c_ activity upon acoustic startle delivery, increases LHb_5HT2c_ excitability in slice, and increases 5HT_2c_ expression in the LHb. 5HT_2c_ in the BNST partially regulates arousal disturbances induced by alcohol, but excessive activity of LHb_5HT2c_ is the primary causal mechanism driving this effect.

